# Multiplexed perturbation of yew reveals cryptic proteins that enable a total biosynthesis of baccatin III and Taxol precursors

**DOI:** 10.1101/2024.11.06.622305

**Authors:** Conor James McClune, Jack Chun-Ting Liu, Chloe Wick, Ricardo De La Peña, Bernd Markus Lange, Polly M. Fordyce, Elizabeth S. Sattely

## Abstract

Plants make complex and potent therapeutic molecules, but difficulties in sourcing from natural producers or chemical synthesis can challenge their use in the clinic. A prominent example is the anti-cancer therapeutic paclitaxel (Taxol^®^). Identification of the full paclitaxel biosynthetic pathway would enable heterologous drug production, but it has eluded discovery despite a half century of intensive research. Within the search space of *Taxus’* large, enzyme-rich genome, we suspected the complex paclitaxel pathway would be difficult to resolve using conventional gene co-expression analysis and small sample sets. To improve the resolution of gene set identification, we developed a multiplexed perturbation strategy to transcriptionally profile cell states spanning tissues, cell types, developmental stages, and elicitation conditions. This approach revealed a set of paclitaxel biosynthetic genes that segregate into expression modules that suggest consecutive biosynthetic sub-pathways. These modules resolved seven new genes that, when combined with previously known enzymes, are sufficient for the *de novo* biosynthesis and isolation of baccatin III, an industrial precursor for Taxol, in *Nicotiana benthamiana* leaves at levels comparable to the natural abundance in *Taxus* needles. Included are taxane 1β-hydroxylase (T1βH), taxane 9α-hydroxylase (T9αH), taxane 7β-*O*-acyltransferase (T7ΑΤ), taxane 7β-*O*-deacetylase (T7dA), taxane 9α-*O*-deacetylase (T9dA), and taxane 9-oxidase (T9ox). Importantly, the T9αH we discovered is distinct and independently evolved from those recently reported, which failed to yield baccatin III with downstream enzymes. Unexpectedly, we also found a nuclear transport factor 2 (NTF2)-like protein (FoTO1) crucial for high yields of taxanes; this gene promotes the formation of the desired product during the first taxane oxidation step, resolving a longstanding bottleneck in paclitaxel pathway reconstitution. Together with a new β-phenylalanine-CoA-ligase, the eight genes discovered in this study enables the complete reconstitution of 3’-*N*-debenzoyl-2’-deoxy-paclitaxel with a 20-enzyme pathway in *Nicotiana* plants. More broadly, we establish a generalizable approach for pathway discovery that scales the power of co-expression studies to match the complexity of specialized metabolism, enabling discovery of gene sets responsible for high-value biological functions.

## Introduction

Plants defend themselves with complex chemical arsenals that have been an essential source of therapeutics^1,2^. Paclitaxel (trade name Taxol^®^) is a potent microtubule-stabilizing agent discovered from *Taxus* plants during the 1960s-1970s National Cancer Institute screening campaign and one of the most valuable chemotherapeutics used in the clinic^3^. Shortly after its approval by the FDA in 1993 for the treatment of ovarian cancer, this diterpenoid became a best-selling pharmaceutical and remains the active component of diverse formulations, derivatizations, and biological conjugates^4^. This extensive use, combined with the chemical complexity and low natural abundance (0.007% DW in *Taxus* bark)^4,5^ of Taxol have made it one of the most sought-after molecules for synthesis. While many elegant synthetic routes have been developed^6^, none are economically viable; drug supply still relies on the extraction from yew tissue, either from plant cell cultures or farming. The promise of a biomanufacturing strategy has made the discovery of the complete *Taxus* enzyme set for heterologous Taxol biosynthesis a grand challenge for natural product chemistry.

The search for the complete Taxol biosynthetic gene set, which was originally proposed to involve 19 enzymes [including 14 enzymes to a key intermediate, baccatin III (**16**)], began in the late 1990s. By 2006, The Croteau lab and others had discovered 12 enzymes, including the scaffold-forming enzyme, taxadiene synthase (TDS), as well as multiple tailoring oxidases and acyltransferases (**Fig. 1a, Table S1**). Progress largely stalled for two decades, until recent reports identified a taxane oxetanase (TOT) that installs Taxol’s unique oxetane moiety, as well as several additional enzymes proposed to act in the pathway^7–11^. However, several of the critical functional groups on Taxol, such as the C-1β hydroxyl, still do not yet have an assigned enzyme with direct biochemical evidence.

**Figure 1.**
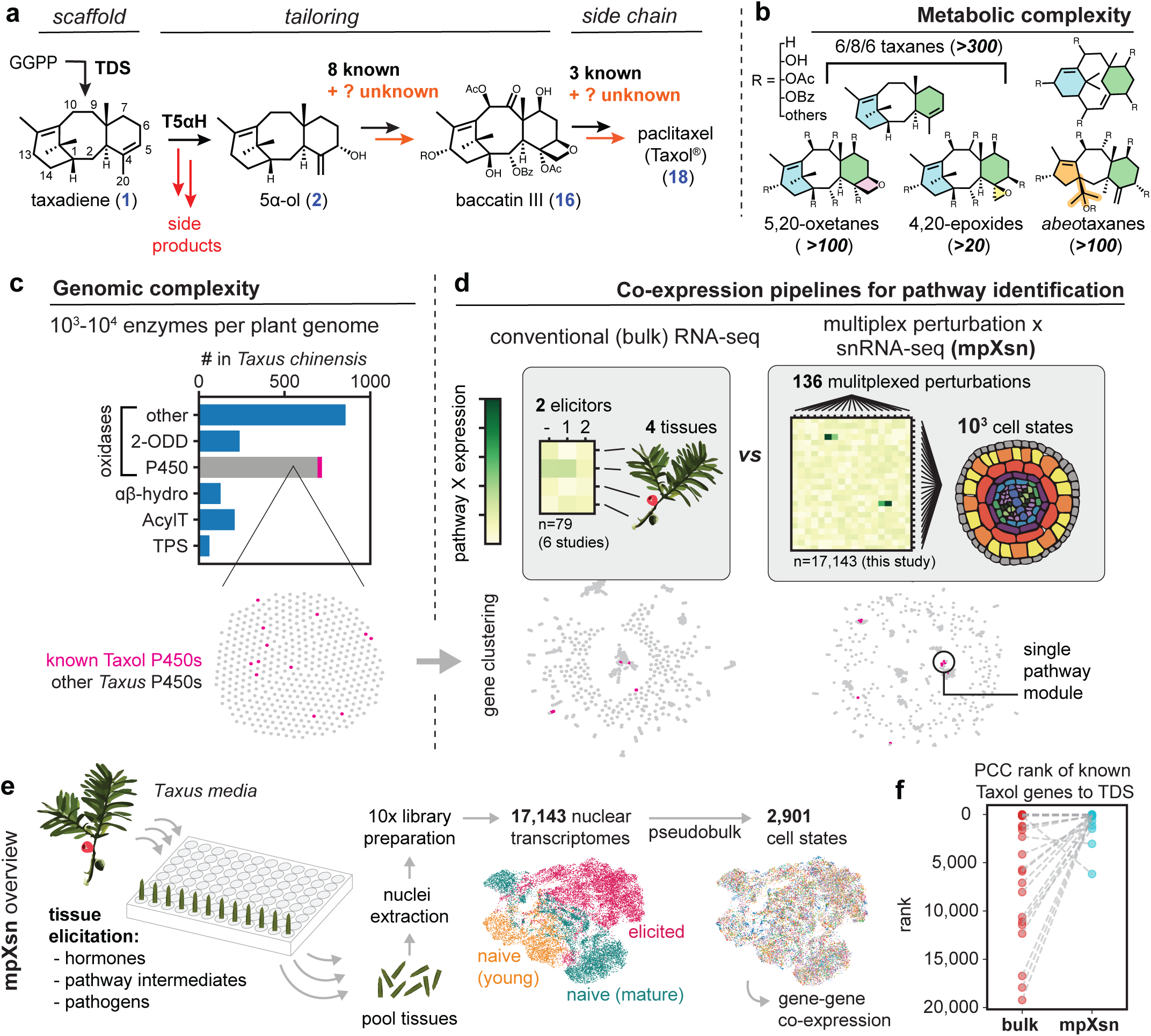
A platform combining multiplexed perturbation and single nuclei RNA-seq (mpXsn) to overcome the challenges of Taxol biosynthetic gene discovery. **(a)** Proposed Taxol biosynthesis pathway with gaps highlighted in orange. In addition to our incomplete knowledge of the biosynthetic gene set, significant inefficiencies (red arrows) of the first oxidase, taxadiene 5α-hydroxylase (T5αΗ), prevent Taxol pathway reconstitution and discovery. TDS, taxadiene synthase. **(b)** Prominent classes of taxane metabolites that have been isolated from Taxus species. The tailoring acyl groups on taxanes include acetyl, benzoyl, small chain fatty acid residues, or phenylisoserine derivatives. **(c)** Gene counts in *Taxus chinensis* genome of the enzyme families known to be involved in taxane biosynthesis: 2-oxoglutarate dependent dioxygenase (2-ODD), cytochrome P450 (P450), ɑβ-hydrolase (αβ-hydro), acyltransferase (AcylT), and terpene synthase (TPS). **(d)** Overview of the differences between conventional co-expression approaches and the mpXsn methodology in this manuscript. Dot networks are visualizations of the co-expression network, where nodes are linked when mutual rank <20, using either bulk RNA-seq or our mpXsn data. For visual clarity, only P450s are displayed. **(e)** Experimental overview for mpXsn with UMAP of single nuclei transcriptomes. **(f)** Rank of each known Taxol gene by Pearson correlation coefficient (PCC) to TDS using either bulk or mpXsn data.

In addition to missing pathway enzymes, heterologous reconstitution of the Taxol pathway has been stymied by inefficiency of the first proposed Taxol oxidation. Despite extensive troubleshooting and optimization efforts in a variety of heterologous systems,^12^ the first Taxol oxidase, taxadiene 5α-hydroxylase, (T5αH), primarily produces side products with rearranged carbon bonds instead of the proposed “on-pathway” intermediate, taxadien-5α-ol (**2**)^13–16^. These two challenges indicate a major gap in our understanding of the endogenous biochemistry (**Fig. 1a**). Both a complete enzyme set and resolution of early oxidation channeling into productive pathway intermediates will be crucial for realizing a heterologous biosynthetic route to Taxol.

The missing components of the Taxol pathway have likely eluded scientists due to the metabolic and genomic complexity of *Taxus.* Taxol is one of almost 600 of taxanes that have been isolated from *Taxus* species^4,17^, including hundreds of 6/8/6 taxanes that differ from Taxol by only subtle tailoring modifications^4^ (**Fig. 1b)**. Within the *Taxus* genome, Taxol pathway genes are just a minute fraction of the hundreds of oxidases, acyltransferases and other enzymes involved in the biosynthesis of primary and secondary metabolites, including numerous taxanes (**Fig. 1c**)^18^.

Transcriptional co-expression approaches that have illuminated other complex plant biosynthetic pathways^19,20^ have not identified the complete Taxol pathway, despite extensive transcriptional profiling of *Taxus*^18,21,22^. Diverse taxanes appear to be co-produced with similar timing, tissue specificity and conditions, suggesting bulk tissue analysis does not provide sufficient resolution to differentiate Taxol enzymes from the thousands of other *Taxus* specialized metabolism genes (**Fig. 1d)**. Improving gene-association specificity would also be necessary if this pathway requires unanticipated genes. To parse large lists of candidate genes, previous efforts have made assumptions about which genes are involved in Taxol biosynthesis, filtering candidates to gene families where previous Taxol enzymes were found, such as the P450 subclass CYP725A^7–11^.

To improve our resolution for identifying gene associations within the taxane metabolic network, we developed an approach to efficiently profile *Taxus* cell states across a vast set of cell types and perturbations. Differential transcriptional activation of *Taxus*’s diverse biosynthetic processes enabled the discovery of multiple Taxol transcriptional modules, from which we identified eight new genes in Taxol biosynthesis. Highlighting the importance of this approach, most identified genes do not belong to previously proposed Taxol gene families. None of the four oxidases identified here derive from the CYP725A family that has been the focus of previous search efforts^7–11^. Three other enzymes are proposed to catalyze the addition and removal of cryptic acetylations that are absent from Taxol, but the inclusion of these enzymes is essential in our reconstituted pathway. Finally, we identified a protein from the NTF2-like family, not previously implicated in plant metabolism, that is crucial for high yields of the productive intermediate during the first taxane oxidation step; this protein resolves the previous challenge of efficient reconstitution the first steps of Taxol biosynthesis^13–16^. With this set of eight genes plus nine previously described, we constructed a pathway in *Nicotiana benthamiana* that produces the prevalent Taxol precursor baccatin III (**16**) at levels comparable to the natural abundance in yew.

## RESULTS

### Multiplexed chemical elicitation produces cells with high expression of known Taxol biosynthetic enzymes

Initially, we developed traditional single-cell transcriptomic methods for *Taxus* and tested whether natural cell type heterogeneity would be sufficient to identify differentially expressed pathways and new Taxol enzyme candidates. The cells of *Taxus* species, like many plants, are often 2–3x larger than the 35 µm diameter limit for the standard 10x Genomics Chromium single-cell library devices. Consequently, we could not use single-cell isolation approaches (e.g. protoplasting) without introducing severe cell-type bias and instead adapted recent nuclei isolation methods^23^ into a conifer-compatible single-nuclei RNA-seqencing (snRNA-seq) protocol (**Methods**). *Taxus media var. hicksii* aerial tissues (needles, stems, and bud scales) were manually disrupted by razorblade and detergent treatment, followed by DNA staining, fluorescence-activated cell sorting (FACS) purification, and library synthesis in the 10x Chromium platform. Initially, we profiled 6,077 cells from unelicited (“naive”) mature tissues. Scanning for Taxol pathway expression revealed that several Taxol biosynthetic enzymes, including TDS, T5αH, and 10-deacetylbaccatin III-10-*O*-acetyl transferase (DBAT) were not highly expressed in any cell (**Fig. S1**). This exemplified one of the core challenges of using transcriptomes to find specialized metabolism genes: tissues must be in a state of active biosynthesis – the conditions for which can be difficult to determine – to capture pathway-associated transcripts^24^. The search for such a tissue state can be risky and challenging, as it may require a specific developmental age^25^ or exposure to a specific biotic stress^24^.

To mitigate the difficulty of identifying biosynthetic cell states by screening large panels of perturbations individually, we leveraged the scale of single-cell transcriptomics into a method we term multiplexed perturbation x single nuclei (mpXsn) that simultaneously test large numbers of perturbations. While single cell transcriptomics have been developed into parallelized screens in mammalian systems^26–28^, we were unable to find such an experimental platform for plants. When developing mpXsn, we also considered the value of a method that requires no genetic tools and would therefore be generalizable to diverse, non-model species. The fact that many plant tissues can be kept alive in liquid media for extended periods of time enables tissues to be subjected, in parallel, to large panels of chemical or biological challenges prior to transcriptome profiling. By combining all tissues into a single snRNA-seq library assembly, individual sample processing is no longer limiting, enabling us to test a large number of samples (272), spanning conditions and time points, in a single experiment (**Fig. 1e**). To maximize the probability of activating biosynthetic states, we compiled a panel of plant hormones (associated with growth, abiotic stress, and biotic stress), pathogen associated molecular patterns (PAMPs), taxane pathway intermediates, and select bacterial and fungal species (**Table S2**). We subjected both young and mature *T. media* needles to this panel for 1, 2, 3, or 4 days before washing and pooling all tissues, extracting nuclei, and generating snRNA-seq libraries. Compared to the naive cell states initially collected, a subset of the elicited cell states now displayed high expression of the early Taxol pathway (**Fig. S1**).

To determine if this single cell transcriptional data provides new information for identifying Taxol enzymes, we directly compared co-expression analysis using either our mpXsn data or bulk RNA-seq data from six previous studies spanning tissues and elicitation conditions (**Methods**). mpXsn data was produced by integrating 8,039 elicited transcriptomes, 3,027 naive transcriptomes from young tissues, and 6,077 naive transcriptomes from mature tissues, clustering these 17,143 profiles into 2,901 cell states using scVI^29^ Leiden clustering, and pooling the reads for each cell state to yield 2,901 pseudobulk RNA-seq profiles. We then determined which dataset showed a stronger correlation between known Taxol genes. Using either bulk RNA-seq data set (79 samples) or the mpXsn (2,901 cell states), we ranked each gene in the *Taxus* genome by Pearson correlation coefficient (PCC) to TDS (**Fig. 1f**). When examining the 14 genes previously associated with Taxol biosynthesis (**Table S1**), all but two genes were prioritized higher using the mpXsn data compared to bulk RNA-seq data (**Fig. 1f, S2-S3**). This analysis suggested the Taxol pathway was better resolved in the mpXsn dataset compared to compiled bulk RNA-seq datasets.

### Identification of three Taxol biosynthetic gene modules

The Taxol pathway has been hypothesized to involve 19 transformations, and previously at least 13 enzymes have been characterized (**Fig. 2a**, **Table S1**). While the first enzyme in the pathway, TDS, does show correlated expression to most Taxol genes (**Fig. 1f, S3**), some known Taxol genes are much more co-expressed than others, forming different co-expressed subsets of the pathway (**Fig. 2b**). These subclusters motivated us to take an untargeted approach to identify gene co-expression modules across the *T. media* transcriptome to determine if the Taxol genes fell into one or more co-expressed gene sets and which other genes were highly co-expressed.

**Figure 2.**
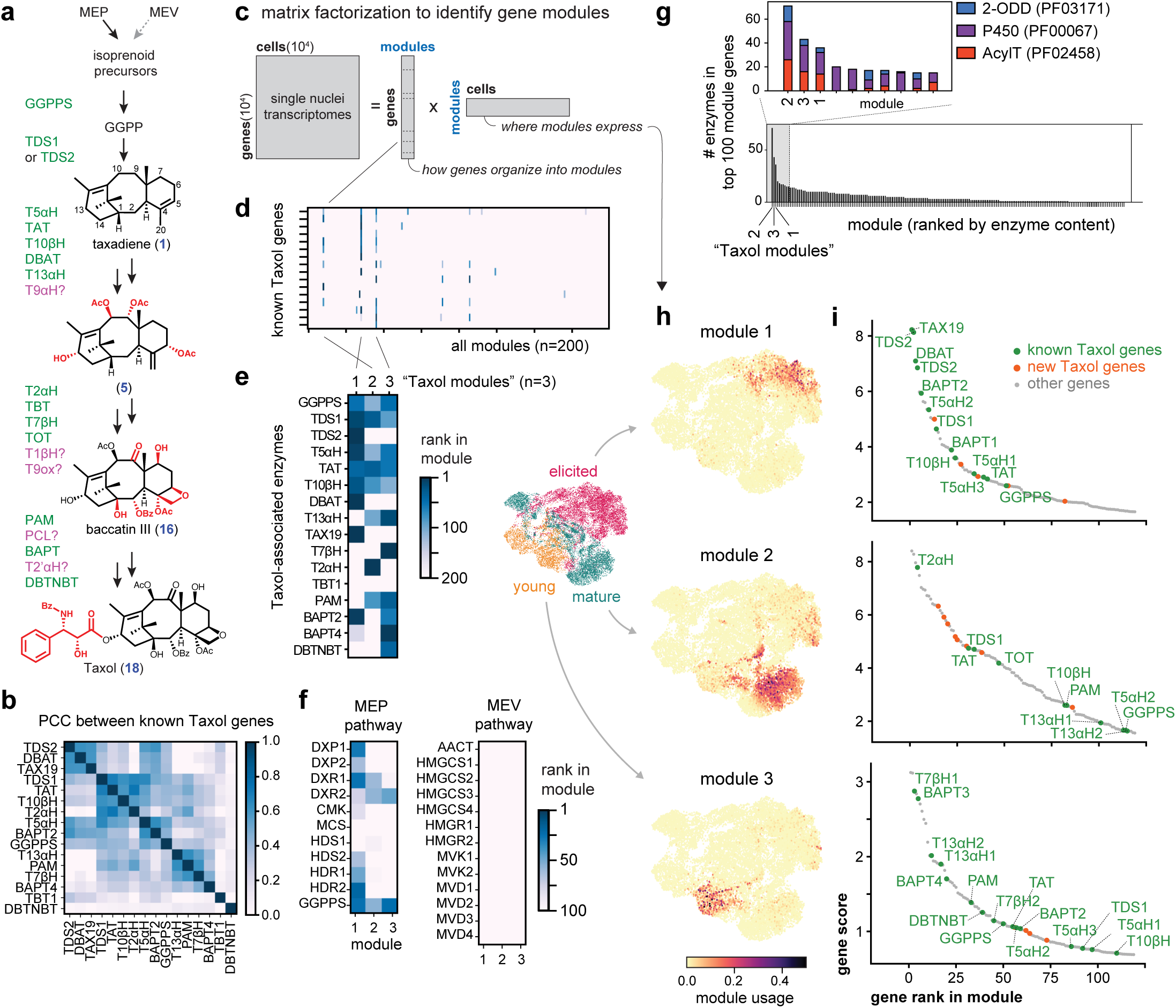
Identification of taxane biosynthetic gene modules. (**a**) Schematic of Taxol biosynthesis and previously hypothesized gene order. (**b**) Correlation between known Taxol genes using mpXsn data. To identify substructures, genes were hierarchically clustered (scipy fcluster, euclidean distance) on both axes. (**c**) Schematic for matrix factorization. mpXsn data factorized using consensus non-negative matrix factorization (cNMF)^30^. (**d**) Heatmap showing known Taxol biosynthetic genes’ rank in each of the modules produced by matrix factorization. (**e**) Same as d, but displaying only the three modules enriched in Taxol genes (hereafter referred to as modules 1, 2, 3). (**f**) Heatmap of Taxol modules showing module rankings for the two isoprenoid pathways in the primary metabolism potentially upstream of the Taxol biosynthesis. Only the MEP pathway is co-expressed with the first Taxol module, supporting its role in synthesizing Taxol precursors. (**g**) All gene modules ranked by total number of 2-ODDs, P450s and acetyltransferases in the top 100 genes of each module. (**h**) Module usage of each cell, which is analogous to gene expression, plotted onto the single nuclei transcriptomic UMAP. Taxol modules 1-3 are expressed in non-overlapping cell states, and were primarily identified in different experiments. (**i**) Unfiltered list of top genes in each module, plotted as module rank and score. Green: previously identified genes associated with Taxol biosynthesis, orange: new biosynthetic genes identified in this study.

Using consensus non-negative matrix factorization^30^ we factored the large gene-by-cell matrix from the mpXsn dataset into submatrices that describe how genes are organized into modules (gene-module score), and how these modules are expressed across cells in the dataset (module-cell score) (**Fig. 2c**). The top scoring genes in each module are coordinately expressed and likely part of the same molecular processes. Of 200 total gene expression modules, we found three where the top genes included many known Taxol enzymes (subsequently called Taxol modules 1, 2, and 3) (**Fig. 2d-e, i**). Interestingly, no single gene module has all the known Taxol enzymes ranked in the top 200 (**Fig. 2e**), suggesting that the full Taxol biosynthesis is regulated by separate transcriptional programs. Furthermore, Taxol module 1 is enriched in genes of the methylerythritol phosphate (MEP) pathway, but not the mevalonate (MEV) pathway, highlighting a linkage of primary and secondary metabolism (**Fig. 2f**). This finding is consistent with current consensus that the MEP pathway supplies precursors for diterpenoids in gymnosperms^31^.

The Taxol modules are highly enriched in enzymes, indicating that these are dominant genetic programs in specialized metabolism. When modules are ranked by the number of enzymes associated with specialized metabolism within their top 100 genes, the three Taxol modules rank first (**Fig. 2g**). Furthermore, these three modules were expressed in different subsets of cells (**Fig. 2h**). Elicitation was crucial for activating module 1, which consists of the early portion of Taxol pathway, as it was not strongly expressed in the naive young and mature *Taxus* tissues we profiled (**Fig. 2h**). An unfiltered analysis of the top genes of the Taxol module 1 revealed all of the genes we had previously used to reconstitute the early Taxol pathway^14^, including TDS, T5αH, taxadien-5α-ol-O-acetyltransferase (TAT), taxane 10β-hydroxylase (T10βH), and DBAT (**Fig. 2i**). We therefore initiated our search for the missing components of the Taxol pathway by examining the uncharacterized genes in this highly co-expressed gene module.

### Discovery of Facilitator of Taxane Oxidization (FoTO1)

One of our primary goals was to determine if there are missing pathway components that could account for the inefficiency of the early Taxol pathway. The first taxane oxidation by T5αH yields a large set of closely related products, most of which appear to not be productive pathway intermediates. Dozens of studies, including from our group, have tried to dissect and troubleshoot this enzyme in a variety of contexts^12–16^. Previously, we reported the first *de novo* reconstitution of the early six Taxol biosynthetic steps, consisting of TDS, T5αH, TAT, T10βΗ, DBAT, and taxane 13α-hydroxylase (T13αΗ), in *Nicotiana benthamiana*, that resulted in the production of 5α,10β-diacetoxytaxadien-13α-ol (**4**) (**Fig. 3a**)^14^. This required extensive expression tuning of T5αΗ that increased yields of the on-pathway intermediate taxadien-5α-ol (**2**)^32^ relative to 5(12)-oxa-3(11)-cyclotaxane (OCT, **2’a**), iso-OCT (**2’b**), and other rearranged side products (**2’c**). Despite our optimization efforts, these side products still accumulated as the dominant products of T5αH^14^. Notably, OCT-derived products are not known to naturally accumulate to significant levels in *Taxus* plants. We hypothesized that Taxol module 1 might contain previously undiscovered proteins that could facilitate this initial oxidation and eliminate the formation of side-products. We tested gene candidates highly ranked in Taxol module 1 (**Fig. 3b**) by expressing them with the six-enzyme early pathway in *N. benthamiana* via *Agrobacterium* mediated transient expression.

**Figure 3.**
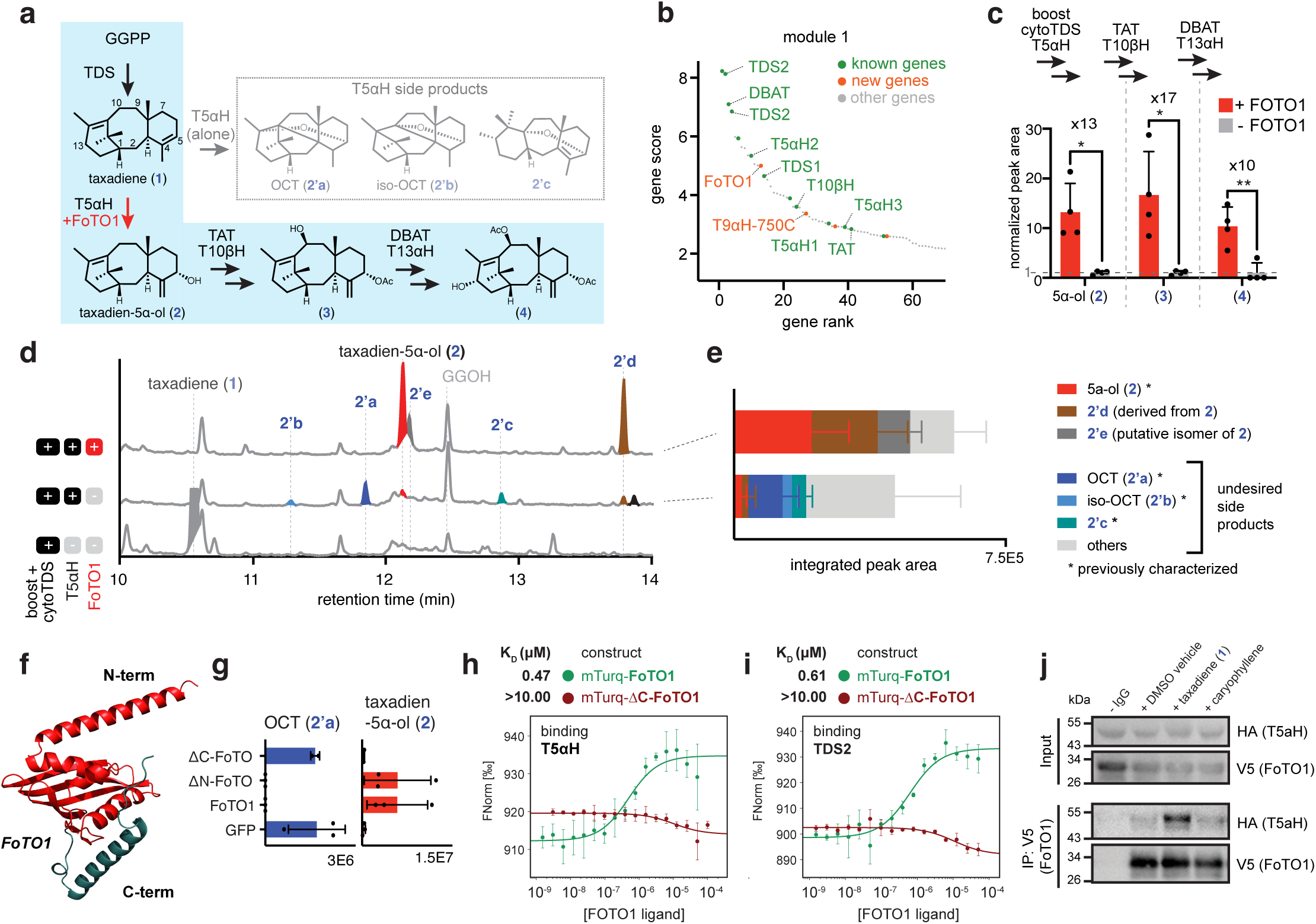
Characterization of FoTO1 (Facilitator of Taxane Oxidization 1). (**a**) Early Taxol biosynthetic pathway and the divergence by T5αH to form multiple side-products, including previously characterized **2’a**, **2’b**, and **2’c**. Blue shade highlights the biosynthetic pathway toward Taxol via **1-4**. (**b**) Rank and score of genes in Taxol module 1 as well as known Taxol genes. (**c**) Bar graph showing the FoTO1-induced fold change in end products’ peak area of subpathways when transiently expressed in N. benthamiana leaves. Fold change is calculated by quantifying **2∼4** GCMS total ion chromatogram (TIC) peak area and normalized to the -FoTO1 condition. Data are shown as the mean ± standard deviation. n = 3 biological, independent leaf samples. Statistical analyses were performed using a two-sided, unpaired Welch’s t-test. * indicates p-value < 0.05, ** indicates p-value < 0.01. (**d**) GCMS TIC of *N. benthamiana* leaves transiently expressing combinations of diterpenoid boost enzymes and cytosolic TDS as the background, T5αΗ, and FoTO. Peaks **2’a-2’e** are side-products of T5αH; **2’d**, while structurally uncharacterized, can be derived by feeding taxadien-5α-ol (**2**) to leaves expressing T5αH.13 (**e**) Bar graph of total oxidized taxanes for +T5αH and +T5αΗ+FoTO1 condition. Data are shown as the mean ± standard deviation. n = 6 biological, independent leaf samples. (**f**) Structure of FoTO1 generated by AlphaFold3. (**g**) Bar graphs showing integrated peak area of taxadien-5α-ol (**2**) and side product OCT (**2’a**) when N- or C-terminal truncated FoTO1 is transiently expressed in *N. benthamiana* leaves together with boost and cytosolic TDS. Expression of FoTO1 and GFP in the same condition are used as positive and negative control, respectively. Data are shown as the mean ± standard deviation, n = 3 biological replicates. (**h**) Quantification of binding between purified T5αΗ and FoTO1 or ΔC-FoTO1 using microscale thermophoresis. (**i**) Quantification of binding between purified TDS2 and FoTO1 or ΔC-FoTO1, as in (**h**). N-terminal transmembrane tags of T5αΗ and TDS2 are removed for purification purpose. Data are shown as the mean ± standard deviation, n = 3. (**j**) Immunoblot analysis of the co-immunoprecipitation (co-IP) of T5αH-HA (prey) by V5-FoTO1 (bait) in *N. benthamiana* leaves expressing both proteins. Incubating lyates with 400 uM taxadiene (**1**) for 15 minutes prior to co-IP increases the pulldown of T5αH by FoTO1 while incubating with a DMSO vehicle or a mock terpene, caryophyllene, does not.

Surprisingly, we observed that gene #13 from module 1, a NTF2-like protein we later named FoTO1 (Facilitator of Taxane Oxidation), resulted in a 10–17 fold increase in yields from early Taxol subpathways (**Fig. 3c**). We hypothesized that FoTO1 may be involved in altering product flux of the early oxidative steps.To determine if FoTO1 ameliorates side product formation, we compared gas chromatography mass spectrometry (GCMS) analysis of *N. benthamiana* leaf extracts expressing TDS and T5αΗ with and without FoTO1. Without FoTO1, T5αΗ yields primarily undesired side products, including **2’a-c** and many uncharacterized compounds, and very little of the desired product, taxadien-5α-ol (**2**) (**Fig. 3d-e**). However, inclusion of FoTO1 drastically alters the product profile: side products **2’a-c** are no longer produced, and taxadien-5α-ol (**2**) and its derivatives (**2’d**) become the major products (**Fig. 3d-e**).

FoTO1 is a 197 amino-acid protein in the NTF2 (nuclear transport factor 2)-like family (**Fig. 3f**), which has not previously been implicated in plant metabolism. While some fungal NTF2 proteins have evolved catalytic activity^33,34^, plant and animal NTF2 proteins have primarily been studied for their capacity to mediate protein transport to the nucleus^35^. Consequently, we anticipated that FoTO1 could be operating via a multitude of mechanisms including (i) scaffolding or allosteric support of Taxol enzymes, (ii) transport or positioning of taxane intermediates, or (iii) enzymatic resolution of an unstable intermediate. FoTO1 homologs from *T. media* and *Arabidopsis thalaiana* did not produce the same metabolomic change for early Taxol pathway reconstitution as FoTO1, suggesting FoTO1 may have a function specific to taxane biosynthesis (**Fig. S4**). When co-expressed *in planta* with either TDS1 or TDS2, FoTO1 has no effect on the production of taxadiene (**1**) and iso-taxadiene, nor does it yield new products (**Fig. S5**), suggesting it has no enzymatic activity on taxadiene (**1**). To further test for potential active sites that could be important for catalysis or substrate binding, we generated mutations of residues within the protein’s cavity based on the predicted AlphaFold structure of FoTO1 (**Fig. 3f, S6**). While none of the amino acid substitutions tested caused FoTO1 to lose its capacity to suppress oxidation side products (**Fig. S6**), deletion of the C-terminal alpha helix, but not the N-terminal helix, eliminated FoTO1’s *in planta* phenotype (**Fig. 3f-g**).

To determine if FoTO1’s function involved a direct interaction with Taxol enzymes and perhaps a scaffolding role, we purified FoTO1 (fused to mTurquoise2) and the soluble portions of TDS2 and T5αΗ. Using microstate thermophoresis, we found that FoTO1 binds both T5αΗ and TDS2 with high nanomolar K_D_ values (**Fig. 3h-i**). Deletion of the C-terminal helix, which disrupts FoTO1’s *in planta* metabolic phenotype (**Fig. 3g**), also eliminated binding affinity with both proteins (**Fig. 3h-i**), suggesting that this region may be involved in protein-protein interactions. To determine if this physical interaction was physiologically relevant, we conducted co-immunoprecipitation (co-IP) of epitope-tagged proteins, V5-FoTO1 and T5αΗ-HA, co-expressed in *N. benthamiana* leaves (**Fig. 3j**). Immunoprecipitation of the bait, V5-FoTO1, using V5 antibody was able to capture T5αΗ-HA (**Fig. 3j, S7**). Furthermore, the T5αΗ-HA co-IP signal increased four-fold when leaf lysates were incubated with 400 μM taxadiene (**1**), but not a mock terpene (caryophyllene) (**Fig. 3j, S7**).

Taken together, these data suggest a mechanism involving a direct interaction between FoTO1, T5αΗ, and possibly TDS. The influence of taxadiene (**1**) on the efficiency of co-IP of T5αΗ and FoTO1 could indicate that the metabolite plays a direct role in such an interaction. While TDS, T5αΗ, and FoTO1 localizes to different subcellular locations – plastid, endoplasmic reticulum (ER), and cytoplasm, respectively (**Fig. S8**) – direct contacts between the outer lamina of these membranes are known to be involved in lipid trafficking^36^ and the biosynthesis of other diterpenoids like gibberellin^37^.

### Independently evolved taxane 9α-hydroxylases (T9αHs) for different taxane substrates

The presence of FoTO1 and all biosynthetic enzymes for (**4**) in gene module 1 suggested coordinated regulation of early Taxol biosynthetic enzymes and an opportunity to discover missing Taxol enzymes from this module (**Fig. 3b**). The next oxidation after C-5α, C-10β, and C-13α hydroxylation is proposed to be the C-9α hydroxylation. Therefore, we screened the activities of the oxidases, including both 2-oxoglutarate dependent dioxygenases (2-ODDs) and P450s, within the top 50 genes of module 1 by co-expressing candidate genes in batches together with the upstream pathway to **4**, a taxane with three oxidation and two acylation modifications (3O2A), in *N. benthamiana*. Through untargeted metabolomic analysis, we found a P450 in the CYP750C family (T9αH-750C) that resulted in depletion of 3O2A (**4**) and, surprisingly, concurrent production of a mass corresponding to a 4O3A (**5**) intermediate that includes both an additional oxidation and an acetylation on **4** (**Fig. 4b**). The additional acetylation was unexpected given that the reconstitution included only two acetyltransferases, TAT and DBAT, which were known mainly to acetylate the C-5 and C-10 hydroxyls, respectively. To confirm the function of T9αH-750C and to provide support for the structural assignment of **5**, we further co-expressed TAX19, the previously characterized C-13α-*O*-acetyltransferase^38^, with the biosynthetic genes to **4** and T9αH-750C at large scale and isolated the resulting acetylated 4O4A products from *N. benthamiana l*eaves. Through extensive 1D- and 2D-nuclear magnetic resonance (NMR) and MS/MS comparison to a standard, we structurally characterized the products as 5α,9α,10β,13α-tetraacetoxytaxadiene, also known as taxusin (**6**), and its isomer 13β-taxusin (**6’**) (**Fig. 4b**, **Table S3**, **Fig. S9**). This structural analysis supports the role of T9αH-750C as a taxane 9α-hydroxylase (T9αH).

**Figure 4.**
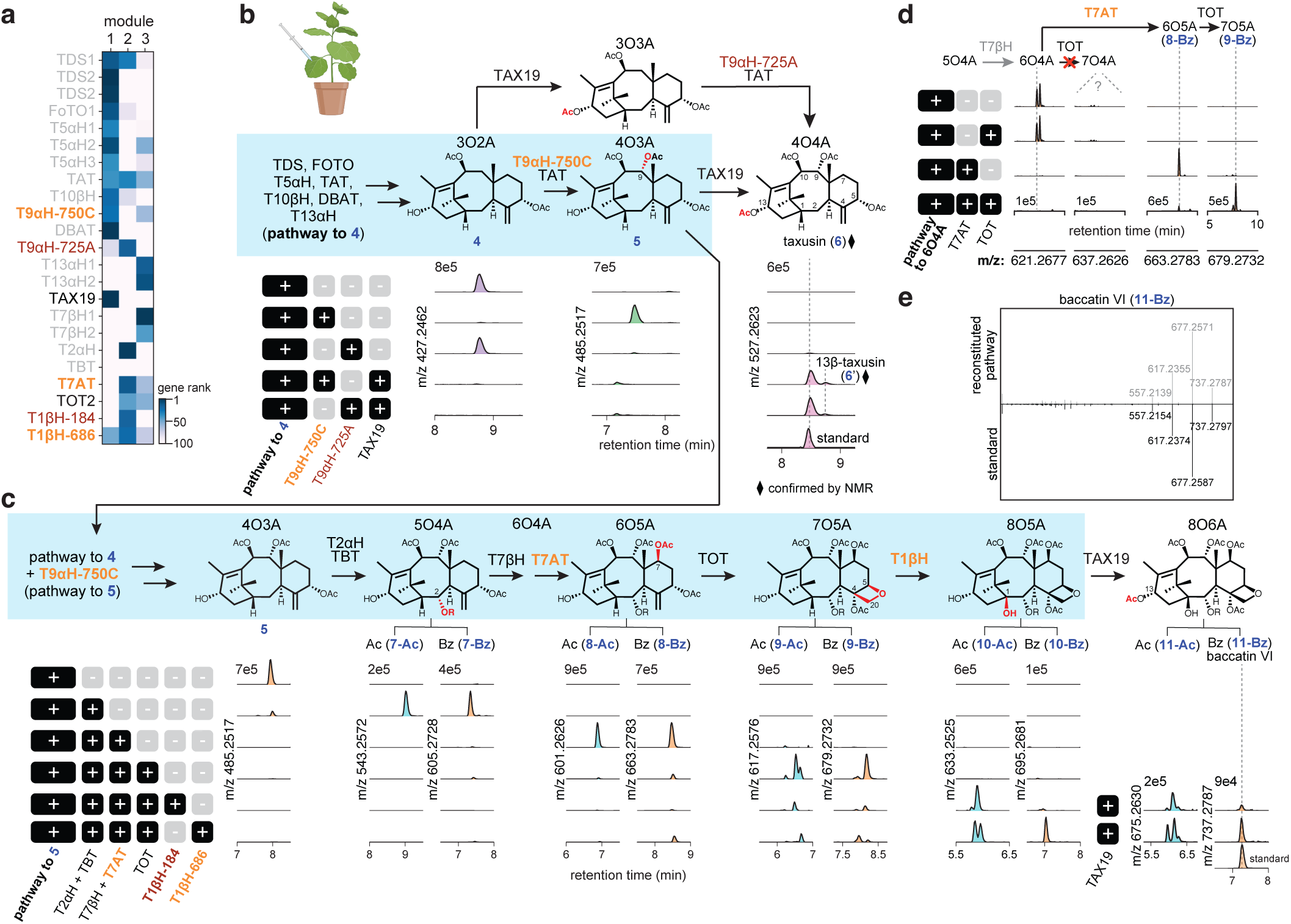
Discovery and characterization of T9αH, T7AT, and two T1βΗs. (**a**) Heat map showing the ranks of new T9α H and T1βHs and other Taxol biosynthetic genes in the three modules. T9αH-725A is the T9αH independently reported by other groups.^8–10^ (**b**) Proposed biosynthetic pathway from **4**, the latest intermediate we reported recently,13 to taxusin (**6**) and the corresponding extracted ion chromatograms (EICs) of products **4-6** when the indicated sets of genes were expressed in *N. benthamiana* leaves. Blue shade highlights the biosynthetic pathway toward Taxol, which only involves **4** and **5**. (**c**) Proposed biosynthetic pathway from **5** to baccatin VI (**11-Bz**) and the corresponding EICs of intermediates when the indicated sets of genes were expressed in N. benthamiana leaves. Blue shade highlights the biosynthetic pathway toward Taxol. Structures of **5**, **7-Ac**, **7-Bz**, **8-Ac**, **8-Bz**, **9-Ac**, **9-Bz**, **10-Ac**, **10-Bz**, **11-Ac** are proposed based on functions of enzymes previously characterized (TAT, TAX19, T2αH, TBT, T7βH, and TOT) and described in this study (Τ9αH-750C, T7AT, and T1βH). TAX19 are used to generate known 13-O-acetylated products, including taxusin (**6**) and baccatin VI (**11-Bz**), for structural analysis. (**d**) EICs of expected products when the pathway to 6O4A is expressed with TOT, T7AT, or both in *N. benthamiana*. In the absence of T7AT, no significant 6O4A depletion or product formation by TOT is observed. (**e**) MSMS fragmentation patterns of heterologously produced baccatin VI (**11-Bz**) in N. benthamiana compared to that of **11-Bz** standard. MSMS fragmentations were generated using [M+Na]+ (m/z = 737.2788) as the precursor ion and fragmented with a collision energy of 30 eV.

Recently, multiple groups have independently reported a distantly related CYP725A P450, with <20% identity at the protein level to T9αH-750C, as the taxane T9αH (referred to here as T9αΗ-725A for distinction), through transcriptome-informed screening of the CYP725A genes, a *Taxus*-specific enzyme family that includes all previously known Taxol P450s^8–10^. Interestingly, while both T9αH-750C and T9αΗ-725A can act as T9αH to produce taxusin (**6**) when TAX19 is present, only T9αΗ-750C can deplete 3O2A to yield 4O3A (**Fig. 4b**, **S10**). In contrast, T9αΗ-725A is unable to produce 4O3A and appears to require a substrate with C-13α acetoxy group (**Fig. 4b, S10**). Extremely low sequence conservation between these enzymes suggests the independent evolution of T9αH activities in two distinct P450 families (CYP725A and CYP750C), and their different substrate specificities suggest that T9αΗ-725A is involved in the biosynthesis of C-13α-acetoxyl taxanes while T9αH-750C is involved in the biosynthesis of C-13α-hydroxyl taxanes. As Taxol and its precursor baccatin III (**16**) lack a C-13α-acetoxy moiety, we used the expression of biosynthetic pathway to **4** and T9αH-750C (i.e., pathway to **5**) in *N. benthamiana* as the foundation for all subsequent taxane pathway reconstructions (**Fig. 4c**).

It is worth noting that the reconstructed pathway to taxusin (**6**) required three acetyltransferases despite four acetoxyl groups on this molecule; these data suggest that one of the acetyltransferases is bi-functional. Experiments involving the feeding of **4** to *N. benthamiana* leaves expressing T9αH-750C with either TAT, or DBAT revealed that TAT is likely a bi-functional enzyme that introduces the acetyl groups on both the C-10β and C9ɑ hydroxyls (**Fig. S11**). This is consistent with previously reported acetylation activities of TAT on C-9 and C-10 hydroxyls.^38^

### Discovery of a taxane 7β-acyltransferase (T7AT) and two taxane 1β-hydroxylases (T1βHs)

The missing C-1β hydroxylation in Taxol is proposed to occur after functionalization by several known mid-pathway enzymes: taxoid 2α-hydroxylase (T2αH) and 7β-hydroxylase (T7βH) have both been shown to independently oxidize various tetra-ol taxane derivatives,^39,40^ taxane 2α-*O*-benzoyltransferase (TBT) has been characterized using 2-debenzoyl baccatin III as the substrate,^41^ and taxane oxetanase (TOT) that installs the signature oxetane ring has been independently reported by multiple groups recently using hexa-acyl taxane (6O6A) as the substrate^7–9^. We hypothesized that these characterized enzymes can function coordinately to reach a hepta-oxidized (7O) taxane intermediate before tailoring of the C-1β hydroxylation, and thus began reconstituting the downstream pathway starting from 4O3A (**5**). Upon co-expression of T2αΗ and TBT with the pathway to **5**, we detected a mass feature corresponding to the expected hydroxylated and benzoylated product (**7-Bz**) (**Fig. 4c**). Intriguingly, we also detected a mass feature corresponding to the acetylated product (**Fig. 4c**), suggesting that TBT is also able to catalyze acetylation. Further testing of TBT with 1-hydroxybaccatin I and baccatin VI showed that TBT is able to mediate the interconversion between C-2α-benzoyl and acetyl group (**Fig. S12**). This bifunctionality is consistent with recent report^42^ and might explain the prevalence of C-2α-acetoxy taxanes in nature.^4,17^

Subsequent expression of T7βH with the 5O4A gene sets yielded the expected benzoylated and acetylated 6O4A. However, adding TOT to the 6O4A pathway did not result in any expected oxidized product (**Fig. 4d**). Given that many of the highly oxygenated taxanes with an oxetane or epoxide moiety originating from TOT activity are also C-7β-*Ο*-acetylated (e.g. baccatin I, baccatin IV, and baccatin VI; **Fig. S13**)^17^, we reasoned that the installation of a C-7β-acetoxy group might be a prerequisite for TOT function. Therefore, we screened acyltransferase candidates and found the 15th gene in module 2 to be capable of various C-7β-*O*-acylations, including acetylation (**Fig. S14**), which we named taxane C-7β-*O*-acyltransferase (T7AT). Expression of T7AT with the upstream 6O4A pathway resulted in the acetylated and benzoylated 6O5A products, **8-Ac** and **8-Bz**, respectively (**Fig. 4c**), and most importantly, further addition of TOT to the 6O5A pathway led to the appearance of dominant mass features corresponding to oxidized products **9-Ac** and **9-Bz** (**Fig. 4c-d**). As TOT has been shown to generate non-interchangeable epoxide and oxetane products^7–9^, it is likely that the two major peaks associated with **9-Ac** and **9-Bz** EIC correspond to those two products. T7AT was independently reported in a recent publication but its importance for TOT function has not been described.^9^

After reconstituting a pathway to **9-Ac** and **9-Bz** (7O5A), we screened the oxidases (2-ODDs and P450s) of module 2 to identify the missing C-1β hydroxylase. This revealed two 2-ODDs (2-ODD184 and 2-ODD686) that yielded multiple mono-oxidized products when expressed with various upstream pathways (**Fig. 4c**, **S15**). To determine if any of these mono-oxidation activities was a C-1β-hydroxylation, we tested whether these enzymes would enable reconstitution of the pathway to baccatin VI (**11-Bz**), an octa-oxygenated taxane (8O6A) containing the C-1β-hydroxyl of interest, which is available as a standard. Expression of the 7O5A pathway and TAX19 (C-13α-*O*-acetyltransferase) with either 2-ODD184 and 2-ODD686 resulted in the production of baccatin VI (**11-Bz**), as confirmed by MSMS comparison to the standard (**Fig. 4c, 4e**) as well as several 1β-hydroxybaccatin I isomers (**11-Ac** peaks, **Fig. S16**). This result suggests that either 2-ODD can function as the missing taxane 1β-hydroxylase (T1βH) since all other functional groups in baccatin VI (**11-Bz**) can be explained by other enzymes included in the reconstitution. Intriguingly, both 2-ODDs resulted in two major products with the 2α-*O*-acetylated pathways while only one major product with the 2α-*O*-benzoylated pathways (**Fig. 4c**, **S15**). Among the two major products in the 2α-*O*-acetylated pathways, one is presumably the 1β-hydroxylated product, but the other remains unidentified. Therefore, we isolated the two products from 2-ODD184 co-expressed with taxusin (**6**) pathway in *N. benthamiana* at large scale and structurally characterized them to be 1β-hydroxytaxusin (**6-Ο1**) and its structural isomer, 15-hydroxy-11(15→1)*abeo*-taxusin (**6-Ο2**) (**Table S4-6, Fig. S17**). We propose that the non-classical 11(15→1)*abeo*taxane scaffold arises from radical rearrangement associated with 2-ODD-mediated 1β-hydroxylation (**Fig. S17**). These data support a role for these 2-ODDs in the C-1β hydroxylation and thus they are referred to as T1βH-184 and T1βH-686 hereafter.

Taken together, these results reveal the discovery of an independently evolved T9αΗ, a T7AT important for the function of TOT, and two T1βHs, which allows us to reconstitute the biosynthesis of highly oxygenated taxanes (**10-Αc** and **10-Bz**) and their C-13α-acetoxy counterparts (**11-Αc** and **11-Bz**). We found that T1βH-686 results in significantly higher levels of 2α-*O*-benzoylated product **10-Bz** compared to T1βH-184, which is desirable for Taxol production, and carried out all subsequent pathway reconstitution with T1βH-686.

### Discovery of two taxane deacetylases and the taxane C-9 oxidase (T9ox) that complete baccatin III biosynthesis

The structure of baccatin III (**16**), the direct precursor to Taxol before side-chain installation, would suggest that it requires nine oxidations (seven hydroxylations, one oxetane formation, and one ketone formation) and three acylations (two acetylations and one benzoylation) on the taxadiene (**1**) scaffold. Using a simplified, two-dimensional oxidation-acylation biosynthesis model (**Fig. 5a**), we can see that our latest intermediate **10-Bz** has two additional acetylations not found in baccatin III (**16**) and is lacking a C-9 ketone oxidation. The two additional acetylations come from (i) TAT that promiscuously *O*-acetylates the C-9-hydroxyl (**Fig. 4b, Fig. S11**), and (ii) T7AT that *O*-acetylates the C-7-hydroxyl, which is vital for the function of TOT (**Fig. 4c-d**). While the roles of these additional acetylations is unknown, we considered that they might serve as protecting groups during the biosynthesis, a strategy utilized in the biosynthesis of other plant terpenoids^43^. We hypothesized that, from **10-Bz**, one oxidase and two deacetylases would then be required for the biosynthesis of baccatin III (**16**) (**Fig. 5a**). To identify such enzymes, we fed substrate baccatin VI (**11-Bz**) or 9-dihydro-13-acetylbaccatin-III (9DHAB, **13**) which structurally resemble our latest intermediate **10-Bz** to *N. benthamiana* leaves expressing top candidate genes in the gene modules to rapidly screen for desired deacetylation and oxidation activities.

**Figure 5.**
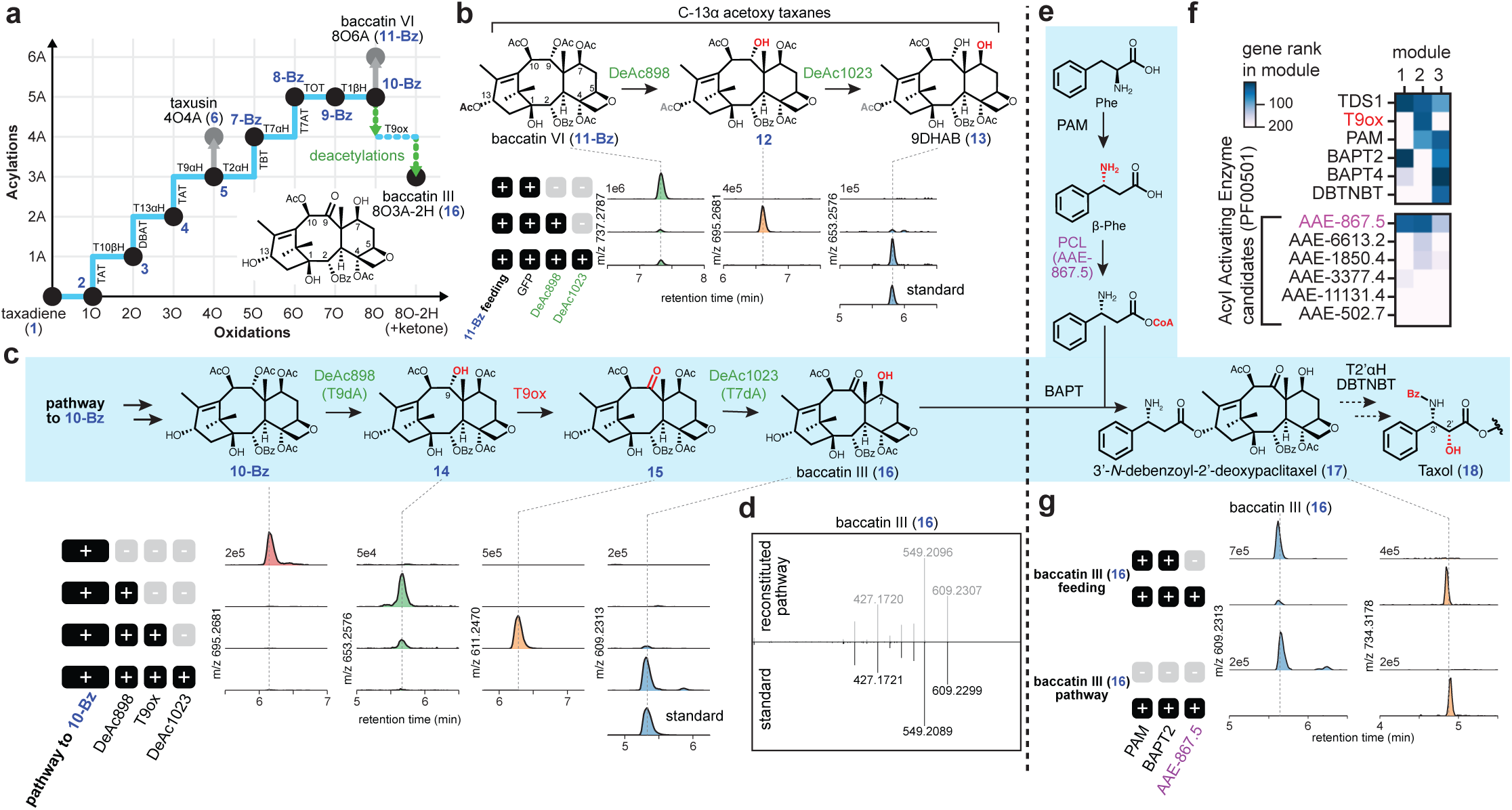
Discovery and characterization of two deacetylases, taxane 9-oxidase (T9ox), and β-phenylalanine-CoA ligase (PCL) that complete the total biosynthesis of 3’-N-debenzoyl-2’-deoxypaclitaxel (17) in N. benthamiana. (**a**) Simplified representation of biosynthetic transformations from taxadiene (**1**) to baccatin III (**16**) with coordinates showing the decoration state (i.e. acylation and oxidation) on the taxadiene (**1**) scaffold. Importantly, two deacetylations and one oxidation are required to go from intermediate 10-Bz to baccatin III (16). Taxanes with C-13 acetoxy, including taxusin (**6**) and baccatin VI (**11-Bz**), are not biosynthetic intermediates to baccatin III (**16**). (**b**) Characterization of the two deacetylases, DeAc 898 and 1023, by feeding baccatin VI (**11-Bz**) to N. benthamiana leaves expressing GFP, DeAc 898 and DeAc 1023. EICs of 11-Bz, the corresponding mono-deacetylated product **12** and di-deacetylated product 9-dihydro-13-acetyl-baccatin III (9DHAB, **13**) are shown. (**c**) Proposed biosynthetic pathway from 10-Bz to baccatin III (**16**) via taxane 9α-O-deacetylase (T9dA; DeAc 898), T9ox, and taxane 7β-O-deacetylase (T7dA; DeAc 1023) and the EICs of intermediates when the indicated sets of genes were expressed in N. benthamiana leaves. (**d**) MSMS fragmentation patterns of heterologously produced baccatin III (16) in N. benthamiana compared to that of 16 standard. MSMS fragmentations were generated using [M+Na]+ (m/z = 609.2313) as the precursor ion and fragmented with a collision energy of 30 eV. (**e**) Proposed side-chain formation and biosynthetic pathway from baccatin III (**16**) to Taxol. Phenylalanine aminomutase (PAM) ‘and PCL convert phenylalanine (Phe) to β-Phe-CoA, which is installed onto baccatin III by baccatin III:3-amino-3-phenylpropanoyl transferase (BAPT) to yield 3’-N-debenzoyl-2’-deoxypaclitaxel (**17**). While taxane 2’α-hydroxylase (T2’αΗ)^46^ and 3′-N-debenzoyl-2′-deoxypaclitaxel-N-benzoyl transferase (DBTNBT) have been previously reported, we were not able to recapitulate their enzymatic activities to complete biosynthetic pathway to Taxol. Only DBTNBT showed activity, and low efficiency suggests that the 2’α-hydroxylation may need to occur first. (**f**) Heatmap showing the ranks of T9ox, side-chain enzymes and PCL candidates in gene modules. The active PCL (867.5) is indicated in magenta. (**g**) 3’-N-debenzoyl-2’-deoxypaclitaxel (**17**) can be produced either by feeding baccatin III (**16**) to N. benthamiana leaves expressing the three side-chain biosynthetic genes (PAM, PCL, and BAPT) or by heterologously expressing the complete biosynthetic gene set to baccatin III (16) plus the three side-chain biosynthetic genes. EICs of 16 and 17 are shown.

Screening of alpha/beta-hydrolase candidates revealed two of these deacetylases (DeAc898 and DeAc1023) were capable of stepwise removal of two *O*-acetylations from baccatin VI (**11-Bz**) to yield a product matching a 9DHAB (**13**) standard, which lacks the C-7 and C-9 *O*-acetyl groups, as supported by MS/MS analysis (**Fig. 5b, Fig. S18**). In addition, expressing the full baccatin VI (**11-Bz**) pathway (rather than fed substrate) with the two deacetylases also resulted in the production of 9DHAB (**Fig. S19**). Screening of oxidase candidates identified a putative taxane C9-oxidase (T9ox) in the 2-ODD family that was capable of oxidizing 9DHAB to the corresponding oxidized product with loss of two protons, presumably via ketone formation at C-9 (**Fig. S20**). This gene has since been independently reported by another group^10^. When we combined T9ox, DeAc898, and DeAc1023 with our 14-enzyme pathway to **10-Bz**, an abundant and significant new taxane product formed (**Fig. S21**), which was subsequently isolated and confirmed as baccatin III (**16**) based on NMR and MS/MS analysis compared to a standard (**Fig. 5c-d, S22)**. Stepwise assembly of the final steps of the pathway revealed DeAc898 to be a prerequisite for T9ox, suggesting this enzyme hydrolyses the C-9 acetyl group (**Fig. 5c**). Consequently, we renamed the DeAc898 and DeAc1023 as taxane 9α-*O*-deacetylase (T9dA) and taxane 7β-*O*-deacetylase (T7dA), respectively.

### Side-chain biosynthetic enzymes enable total biosynthesis of 3’-*N*-debenzoyl-2’-deoxypaclitaxel in *Nicotiana* leaves

The side chain installation and maturation of Taxol have been extensively studied: it is proposed to involve phenylalanine aminomutase (PAM) and β-phenylalanine CoA-ligase (PCL) that convert phenylalanine to β-phenylalanine CoA, which serves as the substrate for baccatin III:3-amino-3-phenylpropanoyl transferase (BAPT). BAPT installs the β-phenylalanine side-chain onto baccatin III (**16**), and ultimately 2’α-hydroxylation and 3’-*N*-benzoylation by taxane 2’α-hydroxylase (T2’αΗ) and 3′-*N*-debenzoyl-2′-deoxypaclitaxel-*N*-benzoyl transferase (DBTNBT), respectively, complete the biosynthesis to Taxol (**18**) (**Fig. 5e, Table S1**)^21,44^. While all five enzymes have been separately reported, we failed to observe the expected product when attempting to reconstitute Taxol biosynthesis by expressing these reported enzymes in combination in *Nicotiana* leaves. Specifically, despite previous reports that two separate *Taxus* acyl activating enzymes (AAE) can act as PCL,^10,21,45^, neither resulted in the production of the 3’-*N*-debenzoyl-2’-deoxypaclitaxel (**17**) when expressed with PAM and BAPT with fed-in baccatin III (**16**) in our system. To identify the missing PCL, we examined the top AAEs in the Taxol expression modules (**Fig. 5f**), revealing a single prominent candidate: AAE-867.5. Co-expression of this PCL with PAM and BAPT *in planta* enabled conversion of fed baccatin III (**16**) to a mass corresponding to 3’-*N*-debenzoyl-2’-deoxypaclitaxel (**17**) (**Fig. 5g**). Furthermore, addition of PCL, PAM and BAPT to our 17-enzyme baccatin III pathway yielded the same dominant product (**Fig. 5g**), which constituted the first *de novo* biosynthetic production of the late-stage paclitaxel precursor 3’-*N*-debenzoyl-2’-deoxypaclitaxel (**17**).

Similarly, while T2’αΗ and DBTNBT have been previously reported^44,46^, we were not able to produce Taxol (**18**) when adding them to the our pathway to **17**. However, when DBTNBT was expressed with PCL, PAM, and DBAT with fed-in baccatin III (**16**) we detected a mass corresponding to 2’-deoxypaclitaxel at very low level (**Fig. S23**). This result suggests that DBTNBT might be functional but requires upstream 2’α-hydroxylation by T2’αH to react efficiently, consistent with previous specificity measurements^47^.

### Expression profile and essentiality of the new Taxol biosynthetic enzymes

Of the eight new Taxol enzymes discovered in this work (**Table S7**), few were clear candidates for the pathway when conducting co-expression analysis using bulk RNA-seq datasets (**Fig. 6a**). Correlation to TDS ranked these genes as 1,000th-10,000th priority with bulk RNAseq, but within the top 10s-100s if using the mpXsn data (**Fig. 6a**), demonstrating that mpXsn strategy was crucial for efficient gene discovery.

**Figure 6.**
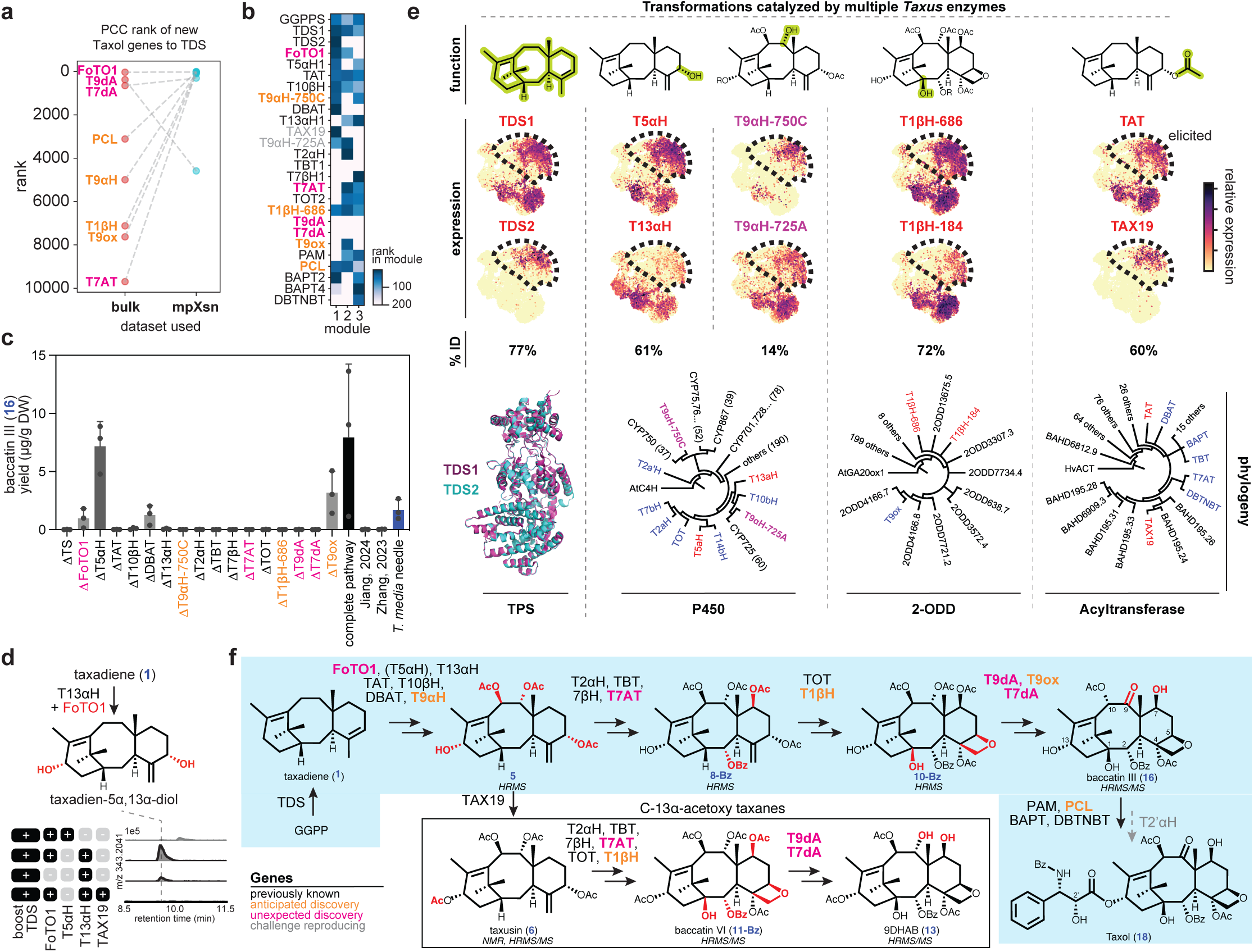
An updated understanding of Taxol biosynthesis with newly discovered genes. (**a**) Ranks of new Taxol biosynthetic gene discovered in this paper by Pearson correlation coefficient (PCC) to TDS using either bulk or mpXsn data. New data from mpXsn was crucial for prioritizing all new Taxol genes except T9dA. Newly discovered genes are indicated in orange (anticipated discovery) and pink (unexpected discovery). (**b**) Heatmap showing the ranks of updated Taxol biosynthetic pathway in the models. (**c**) Baccatin III (**16**) yields in μg/g dried weight (DW), quantified with standards on a triple quadrupole mass spectrometer (**Methods**), of our 17-gene baccatin III pathway with each single-gene dropout tested in *N. benthamiana*. All genes are crucial for significant baccatin III (**16**) yield except T5αH. Baccatin III (**16**) yields from replicating published gene sets^8,10^ as well as from T. media needle are shown for comparison. Data are shown as the mean ± standard deviation, n = 3 biological replicates. (**d**) EIC showing the di-oxidized taxadiene product when T13αΗ is expressed with TDS, and its production is boosted by FoTO1 (**Fig. S26**). Adding TAX19, acyltransferase previously characterized to be specific for taxane C-5α- and C-13α-O-acetylation,^38^ results in the production of a diacetoxytaxadiene (**Fig. S27**). This leads us to propose that T13αH is able to produce taxadien-5α,13α-diol, thus complementing T5αH. (**e**) Five pairs of functional redundant enzymes in the Taxol biosynthetic pathway with their single-cell expression patterns, percentage identity (% ID) in protein sequence, and protein structures or phylogenetic trees shown. Crystal structure of TDS1 (PDB 3P5P) and the AlphaFold3 predicted structure of TDS2 are shown. Phylogenetic trees of the three main tailoring enzyme families involved in Taxol biosynthesis — P450s, 2-ODDs, and acyltransferases. Protein sequences from each family were aligned using Clustal Omega, and the phylogenetic trees were constructed using the neighbor-joining method in Geneious Prime with 1,000 bootstrap replicates. AtC4H, AtGA20ox1, and HvACT were selected as outgroups for each family. (**f**) Summary of the Taxol biosynthesis and the reconstituted C-13α-acetoxy taxane biosynthesis in this study. Blue shade indicates our reconstituted pathway to baccatin III (**16**) and the proposed transformations to Taxol. Co-expression of TAX19 with partial pathways leads to C-13α-acetoxy taxanes including taxusin (**6**), baccatin VI (**11-Bz**), and 9DHAB (**13**) that can be confirmed with chemical standards (Fig. 4e, S9, S19). Structures are proposed based on high resolution MS (HRMS), confirmed by NMR, and/or confirmed by HRMS/MS comparison to standards. Dotted arrow indicated previously reported biosynthetic transformations that we failed to reproduce.

These genes provide new insights into the organization of the biosynthetic pathway by separate transcriptional modules. When ordered by presumed pathway reaction order, the early pathway forms an especially discrete cluster in Taxol module 1 (**Fig. 6b**), suggesting that the pathway up to 4O3A (**5)** may be controlled by separate transcriptional regulation than the later pathway. The latter pathway is not as clustered by order, which could be for multiple reasons: our presumed enzyme order may not reflect the biological pathway or the branching of taxane metabolism towards multiple products may be regulated at various post-transcriptional stages, such as RNA stability or protein localization. Interestingly, some enzymes, such as TDS1, TAT, T1βH-686, are top ranking genes in all three Taxol modules.

For our 17-gene reconstituted baccatin III (**16**) pathway, dropping most of the genes completely abolishes baccatin III (**16**) production, except for FoTO, T5αΗ, DBAT, and T9ox (**Fig. 6c, Fig. S24-25**). Furthermore, in our system, this 17-gene pathway yields levels of baccatin III (**16**) that are significantly greater than those obtained with previously published gene sets (**Fig. 6c, Fig. S24-25**)^8,10^: our attempts to replicate baccatin III (**16**) production with these published gene sets did not yield an amount above the limit of detection in our system (**Fig. S24**). Because these published gene sets lacked several biosynthetic enzymes (e.g. T1βH, T7AT, T7dA, and T9dA), it is possible these findings resulted from multifunctionality of *Taxus* enzymes^11^ or endogenous *N. benthamiana* enzymes. Furthermore, exchange of the T9αH-750C discovered in this work with the alternative T9αH-725A reported in recent publications also yielded no detectable baccatin III (**16**) in our pathway (**Fig. S25-26**), as would be anticipated based on T9αH-750C’s specificity requirement of a C-13α acetoxy that is absent from Taxol and our pathway (**Fig. 4b**).

### Widespread functional overlap of *Taxus* enzymes

Surprisingly, our dropout experiment revealed that T5αH appeared to be nonessential in our reconstituted 17-gene baccatin III (**16**) biosynthesis (**Fig. 6c, Fig. S24-25**). After examining oxidases in the pathway, we found T13αH is able to compensate for T5αΗ in the early pathway and is the only oxidase besides T5αH that can oxidize taxadiene (**1**). Like T5αΗ, T13αΗ yields multiple oxidized taxadiene products including OCT (**2’a)** and iso-OCT (**2’c**) when expressed alone with the taxadiene (**1**) pathway, but not after the introduction of FoTO1 (**Fig. S27**). When co-expressed with FoTO, T13αH produces a di-oxidized taxadiene, which we propose is taxadien-5α,13α-diol (**19**) based on comparison to published MS spectra^48^ and the fact that it can be converted to a diacetoxy-taxadiene by TAX19 (**Fig. 6d, Fig. S28**). This is consistent with previously reported multifunctionality of T13αΗ [converting taxadiene (**1**) to taxadien-5α,10β,13α-triol]^8^ and exemplifies a repeated observation of enzyme functional redundancy during our dissection of the Taxol biosynthetic pathway (**Fig. 6e**).

Beginning with the first committed step of taxane biosynthesis, which can be catalyzed by either of two TDS paralogs,^18^ we found that many taxane functional groups could be installed by multiple genes with different sequences and expression patterns (**Fig. 6e**). In the extreme case of the T9αHs, highly divergent sequences imply convergent evolution. For the T9αH and T1βH variants we found, different substrate and product specificity suggests that these enzymes may be involved in different branches of taxane metabolism, like C-13α-acetoxy taxanes and 11(15→1)*abeo*taxanes. For others, this may just be accidental promiscuity that lacks any evolutionary pressure to correct. A third possibility is that redundant copies of taxane pathway genes act during different conditions, such as at specific developmental stages or after pathogen exposure. Regardless of the mechanism, the prevalence of functional redundancy across the four major enzyme classes in taxane biosynthesis complicates both the dissection of various branches of endogenous taxane metabolism and the identification of optimal enzyme sets for heterologous taxane production.

## DISCUSSION

In this investigation we identified eight new genes in the Taxol biosynthetic pathway, and use them to build 17-gene and 20-gene pathways for *de novo* biosynthesis of baccatin III (**16**) and 3’-*N*-debenzoyl-2’-deoxypaclitaxel (**17**), respectively (**Fig. 6f**), in *Nicotiana benthamiana*.

Future engineering or discovery of the final oxidase, T2’αΗ, would enable *de novo* total biosynthesis of Taxol. Two of the new enzymes, T7AT and T9ox, have recently been independently reported by other groups (**Table S1**).^9,10^ Without optimization, our reconstituted 17-gene pathway yields 10-30 ug/g baccatin III (**16**) in *N. benthamiana* leaves, equivalent to its natural abundance in *Taxus media* needles (**Fig. S29**). Because extraction of baccatin III (**16**) as a semi-synthesis precursor is a dominant industrial strategy for manufacturing Taxol and its derivatives, we believe this work represents a major step towards sustainable production of Taxol and other taxane-based therapeutics.

Our discovery of FoTO1 breaks a long-standing assumption that cytochrome P450s are the sole actors in the first oxidative steps of the Taxol pathway. While all P450s require a cytochrome P450 reductase (CPR) partner, this is the first example, to our knowledge, of a plant P450 that acts in concert with an additional protein to guide product specificity. It is also the first example of NTF2-like proteins being implicated in plant metabolism. Interestingly, a cellulose synthase like protein, GAME15, was recently discovered to be essential for the biosynthesis of *Solanaceae* steroidal molecules and is thought to help orchestrate the function of early pathway enzymes, including two P450s and 2-ODD, on the ER membrane.^49^ Together with our work, this hints at a potentially broader role of protein facilitators in plant secondary metabolism.

Further investigation is needed to dissect how FoTO1 leads to such profound improvements in the activities of multiple *Taxus* oxidases. Co-immunoprecipitation and *in vitro* binding data (**Fig. 3h-j**) suggest the role of direct protein-protein interactions with pathway enzymes, analogous to protein scaffolding role of NTF2-like domains in Ras GTPase activating protein (RasGAP) signaling^50^. While the few characterized NTF2-like proteins in plants also have roles in protein shuttling unrelated to specialized metabolism^35,51^, these proteins differ from FoTO1 in sequence and domain architecture. Thus, we should be cautious with functional extrapolation, given the propensity for this small, conical fold to adapt diverse roles. Beyond the plant kingdom, NTF2-like proteins have independently evolved catalytic activity in fungi and bacteria^33,52^ and have also been computationally engineered into synthetic luciferases^53^. Regardless of its mechanism, FoTO1 has uncharacterized homologs throughout land plants and may represent a conserved family of plant proteins with metabolic roles (**Fig. S30**). It is likely these proteins will be critical for the engineering of complex pathways such as Taxol into heterologous hosts such as yeast for molecule manufacture.

Beyond the unexpected requirement of FoTO1 and the sheer scale of *Taxus* specialized metabolism, multiple obstacles slowed the search for the Taxol biosynthetic enzymes. The number of *Taxus* enzymes that can decorate the taxane scaffold, but not in a way that appears to lead to Taxol biosynthesis, repeatedly lead to dead ends. For example, the T1βH functional homolog T1βH-184 appears to have different substrate and product specificity than the more efficient T1βH-686, suggesting it may act in a different branch of taxane metabolism (**Fig 4c, S15**). Similar expression patterns of these and other taxane enzymes suggest that the branches of the taxane metabolic network may be regulated through posttranslational mechanisms such as protein interactions or metabolons; FoTO1 could be the first of several proteins with scaffolding roles. Furthermore, our long-held model of Taxol biosynthesis requires an update: two additional acetylations appear necessary for the functions of intermediate oxidases, and downstream deacetylations by two deacetylases furnish the baccatin III (**16**) end product (**Fig. 6f**).

While the gene set we identified here heterologously produces baccatin III (**16**) efficiently in *N. benthamiana* via C-13α-hydroxyl intermediates (**Fig. 6f**), it is to be determined if this is the same biosynthetic route adopted in the *Taxus* plants. The prevalence of C-13α-acetoxy taxanes like taxusin (**6**), baccatin VI (**11-Bz**), and 1β-hydroxybaccatin I in *Taxus* plants indicates it is difficult to rule out that the natural biosynthetic pathway might proceed through C-13α-acetoxy intermediates. We were able to access these C-13α-acetoxy taxanes by incorporating the C-13α-*O*-acetyltransferase, TAX19, to our gene set (**Fig. 6f**); however, we were unable to identify a corresponding C-13α-*O*-deacetylase, which prevented us from accessing baccatin III (**16**) via this alternative biosynthetic route.

We overcame the myriad hurdles of Taxol gene discovery by developing a transcriptomic strategy, mpXsn, to more specifically infer gene-gene associations. Scalable profiling of diverse cell states with differentially perturbed biosynthetic processes enabled us to discriminate different enzyme sets and was crucial for prioritizing candidate genes that conflicted with our initial hypothetical biochemical models. In addition to gene discovery, mpXsn data provided biological context, including linkage to primary metabolism (**Fig. 1f**) and the partitioning of Taxol enzymes into modules that appear separately regulated. Beyond Taxol biosynthesis, we anticipate mpXsn to be useful for studying gene sets of interest in other non-model organisms. In mammalian systems, single-cell techniques with parallelized genetic^26^ or chemical^27,28^ perturbation experiments have enabled the de-orphaning of genes and dissecting gene networks. However, most organisms and biological systems lack genetic interrogation tools and rely heavily on observational experiments, such as transcriptomics and other “-omics”. Eukaryotes, especially, face steep challenges for functional genomics and gene-guided discovery as they generally lack the comprehensive gene clusters found in prokaryotes. The advent of methods like mpXsn, which affordably capture precise gene covariance across diverse transcriptional states, may finally overcome this long standing challenge in functional genomics.

## DATA AVAILABILITY

The raw NMR free induction decay (FID) data of individual compounds have been deposited in the Natural Products Magnetic Resonance Database (np-mrd.org) with the following ID: taxusin (**6**, NP0341906), 13β-taxusin (**6’**, NP0341907), 1β-hydroxytaxusin (**6-O1**, NP0341908), 15-hydroxy-11(15→1)abeo-taxusin (**6-O2**, NP0341909); the processed NMR data are shown in **Fig. S32-56** and **Table S3-6.**

## AUTHOR CONTRIBUTION

C.J.M., P.M.F. and E.S.S. conceived of the mpXsn approach used to identify pathway candidates. C.J.M. developed and conducted single-cell procedures, performed transcriptome analysis, selected and cloned gene candidates, and performed initial activity screens. C.J.M. and J.C-T.L. characterized *Taxus* enzymes and analyzed the Taxol pathway products. J.C-T.L. isolated and structurally characterized taxane intermediates by NMR as well as MSMS and performed enzyme phylogenetic analysis. C.W. and C.J.M. performed experiments to characterize the role of FoTO1. B.M.L. shared *Taxus* resources. R.D.L.P. conducted preliminary experiments to establish procedures to characterize *Taxus* enzymes, which B.M.L. helped analyze. E.S.S. and P.M.F. helped analyze the data; C.J.M., J.C-T.L. and E.S.S. wrote the manuscript.

## ACKNOWLEDGEMENTS

We thank all Sattely lab members (2016-2024) for constructive feedback for this project, specifically, Catherine Liou for the help with QQQ data acquisition. We also thank Patrick Almhjell (Stanford University) for helpful comments on this manuscript. We thank Hudson Alpha and the Joint Genome Institute for their work on the *Taxus media cultivar:Hicksii* genome (PRJNA651763), which we used to identify complete versions of some genes. The assembly of the *Taxus media* genome (proposal: 10.46936/10.25585/60001097 led by Joerg Bohlmann) was conducted by the U.S. Department of Energy Joint Genome Institute (https://ror.org/04xm1d337), a DOE Office of Science User Facility, and is supported by the Office of Science of the U.S. Department of Energy operated under Contract No. DE-AC02-05CH11231. We thank Joerg Bohlmann, Jay Keasling, Philipp Zerbe and Anne Osbourn for helpful discussions. We would like to acknowledge Stanford Genomics Core, the Macromolecular Structure Knowledge Center at ChEMH, and Stanford Chemistry NMR facility for use of their instruments. This work is supported by NIH R01 AT010593 (E.S.S.), NIH K99 1K99AT012787 (C.J.M.), and Damon Runyon Cancer Research Foundation DRG: 2421-21 (C.J.M.).

## METHODS

### Chemical and biological materials

Chemical standards are purchased from the following vendors (with catalog number listed): taxusin (TargetMol; TN6763), 1-hydroxybaccatin I (LKT Labs; T0092), baccatin VI (Santa Cruz Biotechnology; sc-503244), 10-deacetylbaccatin III (Sigma-Aldrich; D3676), baccatin III (MedChemExpress; HY-N6985), 9-dihydro-13-acetylbaccatin III (TargetMol; T5132). Taxadien-5α-ol is synthesized as previously described.^14^ *Taxus media var. hicksii* was obtained from FastGrowingTrees.

### Preparation of tissue and single-nuclei sequencing

Nuclei extraction buffer (NIB) consisted of 5 mM MgCl2, 10 mM HEPES pH 7.6, 0.8 M sucrose, 0.1% Triton X and (for density matching to prevent nuclei settling during flow sorting) 1% Dextran T40 and 2% Ficoll. On the day of use, NIB was supplemented with 1 mM dithiothreitol. All nuclei extraction steps were conducted at 4°C and wide bore pipette tips were used when handling nuclei. Steps between tissue harvesting and loading into the Chromium device were completed within 90 minutes to avoid RNA loss. To isolate nuclei, approximately 1g of *Taxus media* tissue was removed from the plant and immediately placed in a petri dish with 10mL NIB. Tissue was chopped at ∼200 rpm with a fresh razor blade for 5 minutes until most large tissue was broken down, and then gently rocked at 4°C for 15 minutes. To remove large debris, disrupted tissue was then passed through a pre-wet 100μm cell strainer stacked atop a 40μm cell strainer. Nuclei were gently pelleted at 300g at 4°C for 5 minutes and resuspended in 1 mL NIB with 5ng/uL 4,6-diamidino-2-phenylindole (DAPI, Thermo Fischer) and 5ng/uL propidium iodide. Using a Sony SH800 cell sorter with a 70μm chip, 140,000-200,000 nuclei were sorted into a tube containing 1mL PBS+ (PBS, 0.1% bovine serum albumin, 20 U/mL Invitrogen Ribonuclease Inhibitor). Gating strategy illustrated in **Fig. S31**. Nuclei were centrifuged at 300g at 4°C for 5 minutes, then gently resuspended in 40uL PBS+. Nuclei were immediately loaded onto a 10x Genomics Chromium controller and libraries were generated using v3 chemistry. Libraries were sequenced on an illumina NextSeq 3000.

### Multiplexed tissue elicitation

For multiplexed elicitation experiment, *Taxus* needles were subjected to perturbation in deep-well 96-well plates with 200 uL MS media [7.5 g/L Murashige and Skoog macronutrients (Fischer), 3 g/L sucrose, pH 5.7]. Two needles (biological replicates) each from two developmental stages (young and mature) were treated with each elicitation condition (17 conditions listed in **Table S2**) for each time point (1, 2, 3 and 4 days), resulting in 272 tissue samples (two replicates of 136 perturbations). To minimize contamination, needles were washed thoroughly in sterile water before moving to MS plates, which were sealed with breathable rayon film (VWR) and placed under 18 hour light cycles.

### Analysis of single-cell data

Reads were mapped to the genomes of *Taxus chinensis*^18^ with STARsolo^54^ (STAR … --runThreadN 32 --alignIntronMa× 10000 --soloUMIlen 12 --soloCellFilter EmptyDrops_CR --soloFeatures GeneFull --soloMultiMappers EM --soloType CB_UMI_Simple). Ambient RNA was removed with cellbender^55^. Using the doubletdetection library^56^, doublets were removed, as well as cells with outlier numbers of reads or where most reads were the most expressed genes (pct_counts_in_top_20_genes < 30). Genes were removed from analysis if expressed in fewer than 50 cells. For integrated UMAP plots, scVI was used to integrate cells from multiple single-cell experiments^29^. Scanpy^57^ was used for processing and plotting post-filtered nuclear transcriptomes. For co-expression analysis and gene-gene correlation calculations, scVI-normalized transcriptomes were clustered into by leiden clustering ^57^ of the, and then raw reads from each cluster were pooled to yield pseudo-bulk transcriptomes. These pseudobulk transcriptomes were used to calculate gene-gene correlations. For module analysis, raw reads were analyzed by consensus non-negative matrix factorization to yield gene modules and their usage across cells.^30^

### Bulk RNA-seq analysis

Raw fastq files for six previous studies^18,58–62^ were downloaded from NCBI, cleaned with trimmomatic^63^ and aligned to the *T. chinensis* genome^18^ (STARmap^64^). Gene-gene correlation was conducted with numpy. Mutual rank (mr), used to calculate the gene linkage maps (**Fig. 1d**) is defined as:

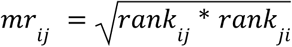

Where rank_ij_ indicates the Pearson correlation rank of gene *i* to gene *j*.

### Cloning of *Taxus* genes

The cloning of cytosolic diterpenoid boost genes (tHMGR and GGPPS), cytosolic TDS1 and TDS2, T5αH, TAT, T10βΗ, DBAT, T13αΗ, and TAX19 genes are described in previous studies.^14,65^ Candidate genes are amplified from *T. media* gDNA or cDNA (generated with SuperScript IV, ThermoFischer) via PCR (PrimeStar, Takara Bio #R045B, primers in **Table S9**), and the PCR products were ligated with AgeI- and XhoI-(New England BioLabs) linearized pEAQ-HT vector^66^ using HiFi DNA assembly mix (New England BioLabs). Gene annotations used for cloning were taken from the *T. chinensis* genome^18^ by default, but were BLAST-searched against the *T. media* genome (NCBI PRJNA1136025) to determine if alternative gene models were available. Constructs were transformed into 10-beta competent *E. coli* cells (New England BioLabs). Plasmid DNA was isolated using the QIAprep Spin Miniprep Kit (Qiagen) and sequence verified by whole-plasmid sequencing (Plasmidsaurus).

### Transient expression of *Taxus* genes in *N. benthamiana* via *Agrobacterium*-mediated infiltration

pEAQ-HT plasmids containing the *Taxus* gene were transformed into *Agrobacterium tumefaciens* (strain GV3101) cells using the freeze-thaw method. Transformed cells were grown on bacteria screening medium 523-agar plates containing kanamycin and gentamicin (50 and 30 μg/mL, respectively; same for the 523 media below), at 30 °C for 2 days. Single colonies were then picked and grew overnight at 30 °C in 523-Kan/Gen liquid media. The overnight cultures were used to make DMSO stocks (7% DMSO) for long-term storage in the −80 °C fridge. For routine *N. benthamiana* infiltration experiments, individual *Agrobacterium* DMSO stocks were streaked out on 523-agar containing kanamycin and gentamicin and grew for 1∼2 days at 30 °C. Patches of cells were scraped off from individual plates using 10 μL inoculation loops and resuspended in 1∼2 mL of *Agrobacterium* induction buffer (10 mM MES pH 5.6, 10 mM MgCl_2_ and 150 μM acetosyringone; Acros Organics) in individual 2 mL safe-lock tubes (Eppendorf). The suspensions were briefly vortexed to homogeneity and incubated at room temperature for 2 h. OD600 of the individual *Agrobacterium* suspensions were measured, and the final infiltration solution where OD600 = 0.2 for each strain (except for TDS, T7AT, and T7dA, whose OD600 were 0.6, 0.4, and 0.1, respectively) was prepared by mixing individual strains and diluting with the induction buffer. Leaves of 4-week old *N. benthamiana* were infiltrated using needleless 1 mL syringes from the abaxial side. Each experiment was tested on leaf-6, 7, and 8 (numbered by counting from the bottom) of the same *N. benthamiana* plant as three biological replicates.

For the reconstitution of pathways that involve TBT, the following modifications were made to the procedure above to increase production of the desired benzoylated products: *N. benthamiana* plants were watered with 2 mM benzoic acid in water (buffered to pH 5.6) a day prior to *Agrobacterium* infiltration, 1 mM benzoic acid was added to the induction buffer and pH adjusted to 5.6 before being used for the resuspension of *Agrobacterium* and preparation of the final infiltration solution.

### Metabolite extraction of *N. benthamiana* leaves

*N. benthamiana* leaf tissue 5-days post *Agrobacterium* infiltration was collected using a leaf disc cutter 1 cm in diameter and placed inside a 2 mL safe-lock tube (Eppendorf). Each biological replicate consisted of 4 leaf discs from the same leaf (approximately 40 mg fresh weight). The leaf discs were flash-frozen and lyophilized overnight. Analysis of the more hydrophobic metabolites, e.g. compound **1**∼**6**, are done by GCMS while analysis of the more hydrophilic metabolites, e.g. **4**∼**18**, are done by LCMS. To extract metabolites, ethyl acetate (ACS reagent grade; J.T. Baker) or 75% acetonitrile (HPLC grade; Fischer Chemical) in water 500 μL was added to each sample along with one 5 mm stainless steel bead for GCMS or LCMS analysis, respectively. The samples were homogenized in a ball mill (Retsch MM 400) at 25 Hz for 2 min. After homogenization, the samples were centrifuged at 18,200 × *g* for 10 min. For GCMS samples, the supernatants were transferred to 50 μL glass inserts placed in 2 mL vials and subjected to GCMS instrument. For LCMS samples, the supernatants were filtered using 96-well hydrophilic PTFE filters with 0.45 μm pore size (Millipore) and subjected to LCMS instrument.

### GCMS analysis

GCMS samples were analyzed using an Agilent 7820 A gas chromatography system coupled to an Agilent 5977B single quadrupole mass spectrometer. Data were collected with Agilent Enhanced MassHunter and analyzed by MassHunter Qualitative Analysis B.07.00. Separation was carried out using an Agilent VF-5HT column (30 m × 0.25 mm × 0.1 μm) with a constant flow rate of 1 ml/min of helium. The inlet was set at 280 °C in split mode with a 10:1 split ratio. The injection volume was 1 μl. Oven conditions were as follows: start and hold at 130 °C for 2 min, ramp to 250 °C at 8 °C/min, ramp to 310 °C at 10 °C/min and hold at 310 °C for 5 min. Post-run condition was set to 320 °C for 3 min. MS data were collected with a mass range 50–550 m/z and a scan speed of 1562 u/s after a 4-min solvent delay. The MSD transfer line was set to 250 °C, the MS source was set to 230 °C and the MS Quad was set to 150 °C.

### LCMS analysis

LCMS samples were analyzed on either or both of our two instruments: (1) an Agilent 1260 HPLC system coupled to an Agilent 6520 Q-TOF mass spectrometer or (2) and Agilent 1290 HPLC system coupled to an Agilent 6546 Q-TOF mass spectrometer. Typically, the 6520 system shows better sensitivity for the more hydrophobic metabolites, e.g. **4**∼**6**, while the 6546 system works better for the more hydrophilic, highly modified taxanes. Data were collected with Agilent MassHunter Workstation Data Acquisition and analyzed by MassHunter Qualitative Analysis 10.0. Separation was carried out using a Gemini 5 μm NX-C18 110 Å column (2 × 100 mm; Phenomenex) with a mixture of 0.1% formic acid in water (A) and 0.1% formic acid in acetonitrile (B) at a constant flow rate of 400 μL/min at room temperature. The injection volume was 2 μl or 1 uL for the 6520 or 6546 system, respectively. The following gradient of solvent B was used: 3% 0–1 min, 3%–50% 1–2 min, 50%–97% 2–12 min, 97% 12–14 min, 97%–3% 14–14.5 min and 3% 14.5–21 min (6520 system) and 3% 0–1 min, 3%–50% 1–5 min, 50%–97% 5–10 min, 97% 10–12 min, 97%–3% 12–12.5 min and 3% 12.5–15 min (6546 system). MS data were collected using electrospray ionization (ESI) on positive mode with a mass range 50–1200 m/z and a rate of 1 spectrum/s (6520 system) or Dual AJS ESI on positive mode with a mass range 100-1700 m/z and a rate of 1 spectrum/s (6546 system). The ionization source was set as follows: 325 °C gas temperature, 10 L/min drying gas, 35 psi nebulizer, 3500 V VCap, 150 V fragmentor, 65 V skimmer, and 750 V octupole 1 RF Vpp (6520 system) or 325 °C gas temperature, 10 L/min drying gas, 20 psi nebulizer, 3500 V VCap, 150 V fragmentor, 65 V skimmer, and 750 V octupole 1 RF Vpp (6546 system).

### Quantification of baccatin III (16)

Samples in **Fig. 6c** was analyzed by an Agilent 1290 HPLC system coupled to an Agilent 6470 triple quadrupole (QQQ) mass spectrometer to accurately quantify the concentration of baccatin III. Data were collected with Agilent MassHunter Workstation Data Acquisition and analyzed by MassHunter Quantitative Analysis 10.1. Separation was carried out using a ZORBAX RRHD Eclipse Plus C18 Column (2.1 x 50 mm, 1.8 µm; Agilent) with a mixture of 0.1% formic acid in water (A) and 0.1% formic acid in acetonitrile (B) at a constant flow rate of 600 μL/min at 30°C. The injection volume was 0.5 μL. The following gradient of solvent B was used: 30% 0–1 min, 30%–100% 1–5 min, 100% 5–6.5 min, 100%–30% 6.5–7 min and 30% 7–8 min. MS data were collected using AJS ESI on positive mode. Multiple reaction monitoring (MRM) scan was used to monitor 609.2 to 549.2 ion transition at 24 eV collision energy as quantifier and 609.2 to 427.1 ion transition at 32 eV collision energy as qualifier. The ionization source was set as follows: 250 °C gas temperature, 12 L/min drying gas, 25 psi nebulizer, 300 °C sheath gas temperature, 12 L/min sheath gas flow, 3500 V VCap, 0 V nozzle voltage.

### Extraction and purification of taxanes from *N. benthamiana*

*N. benthamiana* plants were infiltrated with combinations of biosynthetic genes indicated in **Table S8** for the purification of taxusin (**6**), taxusin (**6’**), 1β-hydroxytaxusin (**6-O1**), and 15-hydroxy-11(15→1)*abeo*-taxusin (**6-O2**). Lyophilied *N. benthamiana* materials were cut into small pieces and extracted with 1 L ethyl acetate (ACS reagent grade; J.T. Baker) in a 2 L flask for 48 h at room temperature with constant stirring. Extracts were filtered using vacuum filtration and dried using rotary evaporation. Two rounds of chromatography were used to isolate compounds of interest. Chromatography conditions for each compound are summarized in **Table S8**. In short, the first chromatography was performed using a 7-cm-diameter column loaded with P60 silica gel (SiliCycle) and using hexane (HPLC grade; VWR) and ethyl acetate as the mobile phases. The second chromatography was carried out on an automated Biotage Selekt system with a Biotage Sfar C18 Duo 6 g column using Milli-Q water and acetonitrile as the mobile phases. Fractions were analyzed by LCMS to identify those containing the compound of interest. Desired fractions were pooled and dried using rotary evaporation (first round) or lyophilization (second round). Purified products were analyzed by NMR.

### NMR analysis of purified compound

CDCl_3_ (Acros Organics) was used as the solvent for all NMR samples. 1H, 13C, and 2D-NMR spectra were acquired on a Varian Inova 600 MHz spectrometer at room temperature using VNMRJ 4.2, and the data were processed and visualized on MestReNova v14.3.1-31739. Chemical shifts were reported in ppm downfield from Me4Si by using the residual solvent (CDCl_3_) peak as an internal standard (7.26 ppm for 1H and 77.16 ppm for 13C chemical shift). Spectra were analyzed and processed using MestReNova version 14.3.1-31739.

### Taxane feeding experiments

*Taxus* genes were expressed in *N. benthamiana* leaves using the *Agrobacterium*-mediated infiltration method described above. At 3-days post *Agrobacterium* infiltration, taxanes [purified 3O2A (**4**), taxusin (**6**), 10-deacetylbaccatin III, or 9-dihydro-13-acetylbaccatin III (**13**); unless otherwise specified, 100 uM solution after diluting 10 mM DMSO stock was used] were fed into the leaves. Approximate 150 μL solution was used per leaf to yield a circle with a diameter around 3 cm, which was marked for reference. After 18–24 h, four leaf discs were harvested within the marked area with a 1-cm diameter cutter and LCMS samples were prepared following the aforementioned methods.

### Construction of phylogenetic tree

Sequences from *Taxus chinensis* genome were selected based on PFAM to identify 672 P450s (PF00067), 218 2-ODDs (PF03171), and 195 acyltransferases (PF02458). P450s are further filtered to those longer than 300 aa (467 P450s). Multiple sequence alignment for each family was performed using Clustal Omega, and the phylogenetic trees were constructed using the neighbor-joining method in Geneious Prime (v2024.0.4) with 100 bootstrap replicates for initial analysis. *Arabidposis thaliana* cinnamate 4-hydroxylase (*At*C4H, accession NP_180607.1), *Arabidposis thaliana* gibberellin 20-oxidase1 (*At*GA20ox1, accession NP_194272.1), and *Hordeum vulgare* agmatine coumaroyltransferase (*Hv*ACT, accession AAO73071.1) are used as outgroup for the P450, 2-ODD, and acyltransferase family, respectively. All analyses were conducted with default settings unless otherwise specified. Representative genes from major clades of the initial analyses and the Taxol biosynthetic genes were then selected to construct the final phylogenetic trees (**Fig. 6e**) using the neighbor-joining method with 1,000 bootstrap replicates.

### Purification of proteins and binding assays

All proteins were purified from standard pET28a vectors expressed in BL21DE3 cells (NEB, C2527H). FoTO1 and FoTO1ΔCterm were purified as C terminal fusions: His6-3xFLAG-TEV-mTurq2-GSG-FoTO1. T5αH and TDS were purified with N-terminal purification tags(His6-3xFLAG-TEV-enzyme) with N-terminal signal peptides removed (T5αH: 47aa removed, TDS2: 60aa removed). Proteins were purified as previously described^67^, with post-lysis steps conducted at 4°C. Briefly, 1 L of cells were grown to OD600 of 0.4-0.5, induced with 0.3 mM IPTG and expressed for 16hr overnight at 18°C. Cell pellets were lysed in lysis buffer (1mg/mL lysozyme, HALT protease cocktail (Thermo Scientific), 1 μL/mL NEB DNAse I) by sonication, clarified by centrifugation for 1hr at 8000g. Proteins were purified on pre-equilibrated Ni-NTA beads (NEB) and exchanged into a protein storage buffer (10 mM HEPES-KOH pH 8.0, 50 mM KCL, 10% glycerol, 1 mM DTT, 1 mM EDTA). Purified proteins were quantified by Bradford assay, and SDS-PAGE gels were used to verify protein size and correct protein concentration.

For each multiscale thermophoresis (MST) experiment, one protein was first labeled with Nanotemper His-tag labeling kit (RED-tris-NTA v2, # MO-L018) for 30 minutes at room temperature according to reagent protocols. MST experiments were performed in PBS with 0.05 % Tween-20 with labeled query protein (T5αH or TDS labeled) at 100 nM and a titration series of target protein.

### Co-Immunoprecipitation

*Nicotiana benthiana* leaves were harvested four days post-infiltration. Leaf tissue was homogenized in liquid nitrogen and resuspended in extraction buffer (50 mM Tris pH 7.5, 150 mM NaCl, 0.6% NP-40, 0.6% CHAPS, 1 mM β-mercaptoethanol [citation below]). Lysates were kept on ice and centrifuged at 20,000g for 10 minutes at 4 °C. The protein content of the clarified extract was determined via Bradford Assay (Abcam #119216). 10 uL of protein G coated magnetic beads (Invitrogen #10003d) were washed twice in binding buffer (50 mM Na2HPO4, 25 mM citric acid, pH 5.0) prior to an hour long room temperature incubation under agitation with 1 uL antibody. Lysates were incubated with the indicated compounds for 15 minutes under agitation. Antibody-bound beads were then washed twice in extraction buffer and incubated for 15 minutes under agitation at room temperature with lysate corresponding to 100 ug of total protein content. Following incubation, bead complexes were washed three times in extraction buffer and mixed with LDS sample buffer (Invitrogen #NP0007) for subsequent analysis by immunoblotting.

### Immunoblotting

Lysates were separated for 1.5 hours at 80 V on a NuPAGE gel (Invitrogen #NP0321) prior to transfer onto a PVDF membrane using a BioRad Trans-Blot Turbo system (BioRad #1704150). Immunoblots were incubated with indicated antibodies for three hours at room temperature under agitation. Blots were subsequently washed and incubated with HRP-Protein G (Genscript #M00090) for one hour, then imaged on the iBright FL1500 Imaging System (Invitrogen #a44241). Extraction buffer was adapted from previously published procedure^68^.

