## Supplementary Figures and Information for "Multiplexed perturbation of yew reveals cryptic proteins that enable a total biosynthesis of baccatin III and Taxol precursors"

[1]Department of Chemical Engineering, Stanford University, California 94305, [2]Department of Chemistry, Stanford University, California 94305, [3]Institute of Biological Chemistry, Washington State University, Pullman, Washington 99164, [4]Department of Bioengineering, Stanford University, California 94305, [5]Department of Genetics, Stanford University, California 94305, [6]Howard Hughes Medical Institute, Stanford University, Stanford, California 94305

†These authors contributed equally

#### Supplementary Table

**Table S1. List of genes used in this study.**

| | Acronym | Full Name | Protein family | Ref. |
|---|---|---|---|---|
| 1 | TDS | taxadiene synthase | terpene cyclase | [1] |
| 2 | T5αH | taxadiene 5α-hydroxylase | P450 | [2] |
| 3 | FoTO1 | facilitator of taxane oxidation | NTF2-like | This work |
| 4 | TAT | taxadien-5α-ol-*O*-acetyltransferase | BAHD acyltransferase | [3] |
| 5 | T10βH | taxane 10β-hydroxylase | P450 | [4] |
| 6 | DBAT | 10-deacetylbaccatin III-10β-*O*-acetyltransferase | BAHD acyltransferase | [5] |
| 7 | T13αH | taxane 13α-hydroxylase | P450 | [6] |
| 8 | T9αH-750C | taxane 9α-hydroxylase-750C | P450 | This work |
| | T9αH-725A | taxane 9α-hydroxylase-725A | P450 | [7–9] |
| 9 | T2αH | taxoid 2α-hydroxylase | P450 | [10] |
| 10 | TBT | taxane 2α-*O*-benzoyltransferase | BAHD acyltransferase | [11] |
| 11 | T7βH | taxoid 7β-hydroxylase | P450 | [12] |
| 12 | T7AT | taxane 7β-*O*-acetyltransferase | BAHD acyltransferase | This work [8] |
| 13 | TOT | taxane oxetanase | P450 | [7,8,13] |
| 14 | T1βH-184 | taxane 1β-hydroxylase-184 | 2-ODD | This work |
| | T1βH-686 | taxane 1β-hydroxylase-686 | 2-ODD | This work |
| 15 | T9dA | taxane 9α-*O*-deacetylase | Alpha/beta hydrolase | This work |
| 16 | T9ox | taxane C9-oxidase | 2-ODD | This work [9] |
| 17 | T7dA | taxane 7β-*O*-deacetylase | Alpha/beta hydrolase | This work |
| 18 | PAM | phenylalanine aminomutase | ammonia-lyases | [14] |
| 19 | PCL | phenylalanine-CoA ligase | AMP binding / acyl-activating | This work |
| 20 | BAPT | baccatin III:3-amino-3-phenylpropanoyl transferase | BAHD acyltransferase | [15] |
| 21 | T2’αH | taxane 2’α-hydroxylase | P450 | [16*] |
| 22 | DBTNBT | 3′-*N*-debenzoyl-2′-deoxypaclitaxel-*N*-benzoyl transferase | BAHD acyltransferase | [17] |

P450: cytochrome P450; 2-ODD: 2-oxoglutarate-dependent dioxygenase; NTF2: nuclear transport factor 2

*Enzymatic activity not observed with our reconstituted pathway.

**Table S2.** Panel of elicitation conditions

| condition # | class | perturbation | concentration |
|---|---|---|---|
| 1 | vehicle | MS media | - |
| 2 | biotic stress hormone | methyl jasmonate | 10 µM |
| 3 | biotic stress hormone | methyl jasmonate | 100 µM |
| 4 | biotic stress hormone | methyl jasmonate | 1 mM |
| 5 | biotic stress hormone | methyl jasmonate | 10 mM |
| 8 | biotic stress hormone | N-hydroxypipecolic acid | 2 mM |
| 9 | abiotic stress hormone | abscisic acid | 1 mM |
| 10 | biotic stress hormone | salicylic acid | 2 mM |
| 12 | growth hormone | trans-zeatin | 10 µM |
| 11 | PAMP (pathogen associated molecular pattern) | chitosan | 1% |
| 6 | PAMP | Flg22 (Stanford PAN facility) | 1 µM |
| 7 | PAMP | Flg22 | 10 µM |
| 13 | taxane | paclitaxel | 5 µM |
| 14 | taxane | baccatin III (MedChemExpress) | 100 µM |
| 15 | microbial challenge | Aspergillus ssp. (**Supplemental Note 1**) | OD600 0.1 |
| 16 | microbial challenge | Aspergillus ssp. | OD600 0.1 |
| | | methyl jasmonate | 100 µM |
| 17 | microbial challenge | *Agrobacterium tumefaciens* GV3101 | OD600 0.01 |

**Table S3.** $^{13}$C and $^{1}$H δ assignments of taxusin (**6**) and 13β-taxusin (**6'**) recorded in $CDCl_3$.

| | 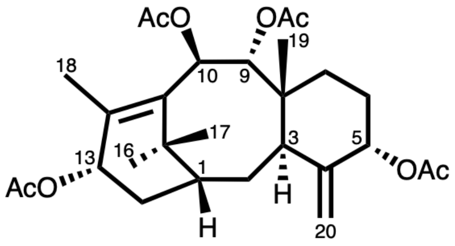  taxusin (**6**) | | 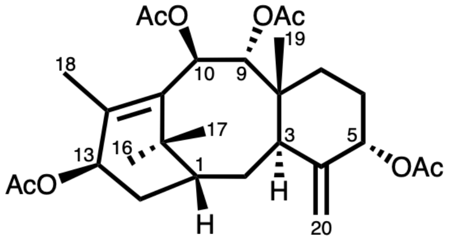  13β-taxusin (**6'**) | |
|---|---|---|---|---|
| **C-#** | **δ $^{13}$C (ppm)** | **δ $^{1}$H (mult.; *J* in Hz)** | **δ $^{13}$C (ppm)** | **δ $^{1}$H (mult.; *J* in Hz)** |
| 1 | 40.3 | 1.84 (m) | 38? | 2.00 (m) |
| 2a | 27.3 | 1.79 (m) | N.A. | 1.73 (m) |
| 2b | 27.3 | 1.69 (m) | N.A. | 1.84 (m) |
| 3 | 37.9 | 3.00 (d; 6.5) | 38.7 | 2.80 (m) |
| 4 | 148.3 | -- | N.A. | -- |
| 5 | 76.3 | 5.36 (t; 2.5) | 76 | 5.38 (br, s) |
| 6a | 27.3 | 1.84 (m) | N.A. | 1.84 (m) |
| 6b | 27.3 | 1.68 (m) | N.A. | 1.69 (m) |
| 7 | 27.3 | 1.76 (m) | 27.2 | 1.74 (m) |
| 8 | 42.8 | -- | 43.4 | -- |
| 9 | 70.8 | 5.88 (d; 10.7) | 76.9 | 5.83 (d; 10.5) |
| 10 | 72.5 | 6.08 (d; 10.7) | 73 | 6.03 (d; 10.5) |
| 11 | 134.3 | -- | 140.8 | -- |
| 12 | 136.5 | -- | 136.3 | -- |
| 13 | 77.5 | 5.87 (m) | 72.7 | 5.33 (dd; 4.3, 9.6) |
| 14a | 31.9 | 1.06 (dd; 7.5, 14.5) | N.A. | 1.94 (dd; 9.6, 15.4) |
| 14b | 31.9 | 2.69 (dt; 14.6, 9.8) | N.A. | 2.10 (m) |
| 15 | 39.2 | -- | 47.8 | -- |
| 16 | 30.6 | 1.11 (s) | 35.6 | 1.25 (s) |

| | | | | |
|---|---|---|---|---|
| 17 | 26.7 | 1.62 (s) | 26.1 | 1.55 (s) |
| 18 | 14.9 | 2.11 (s) | 18.7 | 2.05 (s) |
| 19 | 17.7 | 0.75 (s) | 17.2 | 0.74 (s) |
| 20a-(E) | 114 | 4.85 (s) | 113.4 | 4.84 (s) |
| 20b-(Z) | 114 | 5.21 (s) | 113.4 | 5.20 (s) |
| -OCOMe | 21 | 2.01 (s) | 21.1 | 2.03 (s) |
| -OCOMe | 20.7 | 2.05 (s) | 20.7 | 2.05 (s) |
| -OCOMe | 21.6 | 2.06 (s) | 21.5 | 2.08 (s) |
| -OCOMe | 21.7 | 2.16 (s) | 21.5 | 2.14 (s) |
| -OCOMe | 169.4 | -- | 169.9 | -- |
| -OCOMe | 169.4 | -- | 170.2 | -- |
| -OCOMe | 169.9 | -- | 170.4 | -- |
| -OCOMe | 169.9 | -- | 171.1 | -- |

NMR spectra are shown in **Fig. S32-37** (taxusin) and **Fig. S38-43** (13β-taxusin).
s = singlet, d = doublet, dd = doublet of doublets, dt = doublet of triplets, t = triplet, q = quartet, quint = quintet, m = multiplet

**Table S4.** $^{13}C$ and $^{1}H$ δ assignments as well as 2D-NMR correlations of 1β-hydroxytaxusin (**6-O1**) recorded in $CDCl_3$.

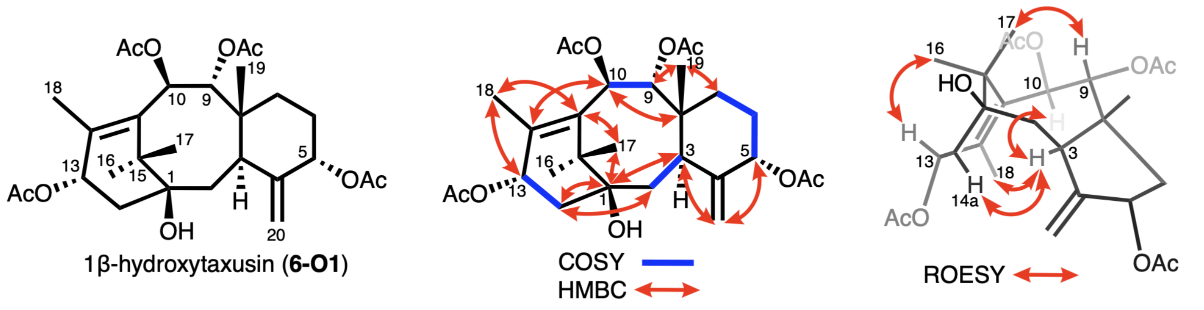


1β-hydroxytaxusin (**6-O1**)

| C-# | δ $^{13}C$ (ppm) | δ $^{1}H$ (mult.; *J* in Hz) | COSY | HMBC | key ROESY |
|---|---|---|---|---|---|
| 1 | 76.3 | -- | -- | -- | -- |
| 2a | 38.3 | 1.79 (d; 15.2) | 2b | 1, 3, 4, 8 | -- |
| 2b | 38.3 | 1.91 (dd; 6.7, 15.2) | 2a,3 | 1, 3, 8 | -- |
| 3 | 41.7 | 2.94 (d; 6.7) | 2b | 1, 2, 4, 5, 8, 19, 20 | 6a, 10, 14a, 18 |
| 4 | 148.0 | -- | -- | -- | -- |
| 5 | 76.4 | 5.38 (t; 2.6) | 6b | 3, 4, 6, 7, 20 | 6b, 7 |
| 6a | 27.4 | 1.74 (dd; 4.4, 11.0) | 7 | 19 | -- |
| 6b | 27.4 | 1.85 (m) | 7 | -- | -- |
| 7 | 27.4 | 1.70 (m) | 6a, 6b | -- | -- |
| 8 | 43.4 | -- | -- | -- | -- |
| 9 | 77.0 | 5.88 (d; 10.7) | 10 | 7, 8, 10, 19 | 2b, 17, 19 |
| 10 | 72.2 | 6.08 (d; 10.7) | 9 | 8, 9, 11, 12, 15 | 3, 6a, 18 |
| 11 | 134.7 | -- | -- | -- | -- |
| 12 | 139.1 | -- | -- | -- | -- |
| 13 | 71.1 | 6.03 (t; 8.5) | 14a, 14b, 18 | 11, 12, 14 | 14b, 16 |
| 14a | 41.6 | 1.60 (dd; 7.1, 14.8) | 13, 14b | 1, 2, 13, 15 | -- |
| 14b | 41.6 | 2.53 (dd; 9.8, 14.8) | 13, 14a | 1, 2, 12, 13 | -- |
| 15 | 43.8 | -- | -- | -- | -- |
| 16 | 27.4 | 1.2 (s) | -- | 1, 11, 15, 17 | -- |
| 17 | 22.1 | 1.62 (s) | -- | 1, 11, 15, 16 | -- |
| 18 | 14.9 | 2.10 (s) | -- | 11, 12, 13 | -- |
| 19 | 17.9 | 0.76 (s) | -- | 3, 7, 8, 9 | -- |
| 20a-(E) | 114.4 | 4.92 (s) | 20b | 3, 4, 5 | 2a, 20b |
| 20b-(Z) | 114.4 | 5.25 (s) | 20a | 3, 5 | 20a |
| -OCOMe | 20.7 | 2.05 (s) | -- | -- | -- |

| | | | | | |
|---|---|---|---|---|---|
| -OCOMe | 21.1 | 2.02 (s) | -- | -- | -- |
| -OCOMe | 21.7 | 2.08 (s) | -- | -- | -- |
| -OCOMe | 21.7 | 2.16 (s) | -- | -- | -- |
| -OCOMe | 169.8 | -- | -- | -- | -- |
| -OCOMe | 169.8 | -- | -- | -- | -- |
| -OCOMe | 170.2 | -- | -- | -- | -- |
| -OCOMe | 170.3 | -- | -- | -- | -- |
| C1-OH | -- | 1.42 (s) | -- | 1, 2, 14 | -- |

NMR spectra are shown in **Fig. S44-49**.

s = singlet, d = doublet, dd = doublet of doublets, dt = doublet of triplets, t = triplet, q = quartet, quint = quintet, m = multiplet

**Table S5.** $^{13}$C and $^{1}$H δ assignments as well as 2D-NMR correlations of 15-hydroxy-11(15→1)*abeo*-taxusin (**6-O2**) recorded in $CDCl_3$.

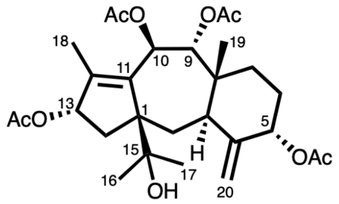


5α,9α,10β,13α-tetraacetoxy-
15-hydroxy-11(15→1)-*abeo*-texa-4(20),11-diene
[15-hydroxy-11(15→1)*abeo*-taxusin, **6-O2**]

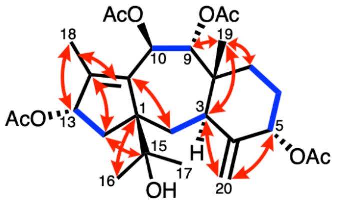


COSY
HMBC

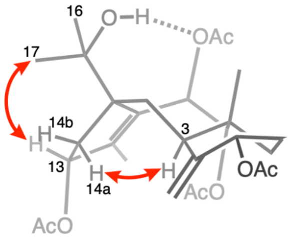


ROESY

| C-# | δ $^{13}$C (ppm) | δ $^{1}$H (mult.; *J* in Hz) | COSY | HMBC | key ROESY |
|---|---|---|---|---|---|
| 1 | 62.8 | -- | -- | -- | -- |
| 2a | N.A. | 1.38 (d; 14.4) | 2b | 1, 8, 4, 11 | -- |
| 2b | N.A. | 2.17 (dd; 8.8, 14.4) | 2a, 3 | 3 | -- |
| 3 | 39.7 | 2.7 (d; 8.6) | 2b | 4, 20 | 14a |
| 4 | 146.6 | -- | -- | -- | -- |
| 5 | 74.7 | 5.31 (t; 2.7) | 6a, 6b | -- | -- |
| 6a | N.A. | 1.79 (m) | 5, 6b | -- | -- |
| 6b | N.A. | 1.88 (m) | 5, 6a | -- | -- |
| 7 | 27.4 | 1.69 (m) | 6a, 6b | -- | -- |
| 8 | 42.1 | -- | -- | -- | -- |
| 9 | 77.6 | 5.78 (d; 10.2) | 10 | -- | 2b, 19, C15-OH |
| 10 | 69.2 | 6.16 (d; 10.2) | 9 | -- | 18 |
| 11 | 137.4 | -- | -- | -- | -- |
| 12 | 144.8 | -- | -- | -- | -- |
| 13 | 79.3 | 5.53 (t; 7.5) | 14a, 14b, 18 | -- | 14b, 17 |
| 14a | 44.3 | 1.22 (dd; 7.7, 13.9) | 13, 14b | 1, 13, 15 | -- |
| 14b | 44.3 | 2.49 (dd; 7.3, 13.9) | 13, 14a | 12, 15 | -- |
| 15 | 75.2 | -- | -- | -- | -- |
| 16 | 24.6 | 1.32 (s) | -- | 1, 15, 17 | -- |
| 17 | 27 | 1.15 (s) | -- | 1, 15, 16 | -- |
| 18 | 11.5 | 1.83 (s) | -- | 11, 12, 13 | -- |
| 19 | 16.5 | 0.78 (s) | -- | 3, 7, 8, 9 | -- |
| 20a-(E) | 111.9 | 4.76 (s) | 20b | -- | 2a, 20b |
| 20b-(Z) | 111.9 | 5.17 (s) | 20a | 3, 5 | 20a |
| -OCOMe | 20.8 | 2.00 (s) | -- | -- | -- |

| | | | | | |
|---|---|---|---|---|---|
| -OCOMe | 20.8 | 2.01 (s) | -- | -- | -- |
| -OCOMe | 20.8 | 2.02 (s) | -- | -- | -- |
| -OCOMe | 21.3 | 2.04 (s) | -- | -- | -- |
| -OCOMe | 168.4 | -- | -- | -- | -- |
| -OCOMe | 169.4 | -- | -- | -- | -- |
| -OCOMe | 170.9 | -- | -- | -- | -- |
| -OCOMe | 170.9 | -- | -- | -- | -- |
| C15-OH | -- | 2.42 (s) | -- | -- | 9, 17 |

NMR spectra are shown in **Fig. S50-56**.
N.A.: chemical shift not assigned due to overlapping signals.
s = singlet, d = doublet, dd = doublet of doublets, dt = doublet of triplets, t = triplet, q = quartet, quint = quintet, m = multiplet
This assignment is consistent with previously reported assignment of the same compound in $CD_3OD$.[18]

**Table S6.** Comparison of the $^{1}H$ δ assignments of taxusin (**6**), 1β-hydroxytaxusin (**6-O1**), and 15-hydroxy-11(15→1)*abeo*-taxusin (**6-O2**).

| | 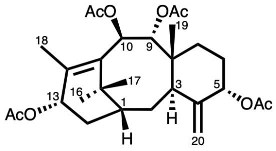 taxusin (**6**) | 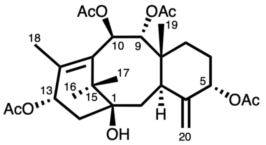 1β-hydroxytaxusin (**6-O1**) | 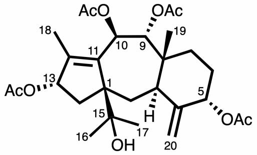 5α,9α,10β,13α-tetraacetoxy-15-hydroxy-11(15→1)-*abeo*-texa-4(20),11-diene [15-hydroxy-11(15→1)*abeo*-taxusin, **6-O2**] |
|---|---|---|---|
| **C-#** | **δ $^{1}$H (mult.; *J* in Hz)** | | |
| 1 | 1.84 (m) | -- | -- |
| 2a | 1.79 (m) | 1.79 (d; 15.2) | 1.38 (d; 14.4) |
| 2b | 1.69 (m) | 1.91 (dd; 6.7, 15.2) | 2.17 (dd; 8.8, 14.4) |
| 3 | 3.00 (d; 6.5) | 2.94 (d; 6.7) | 2.7 (d; 8.6) |
| 4 | -- | -- | -- |
| 5 | 5.36 (t; 2.5) | 5.38 (t; 2.6) | 5.31 (t; 2.7) |
| 6a | 1.84 (m) | 1.74 (dd; 4.4, 11.0) | 1.79 (m) |
| 6b | 1.68 (m) | 1.85 (m) | 1.88 (m) |
| 7 | 1.76 (m) | 1.70 (m) | 1.69 (m) |
| 8 | -- | -- | -- |
| 9 | 5.88 (d; 10.7) | 5.88 (d; 10.7) | 5.78 (d; 10.2) |
| 10 | 6.08 (d; 10.7) | 6.08 (d; 10.7) | 6.16 (d; 10.2) |
| 11 | -- | -- | -- |
| 12 | -- | -- | -- |
| 13 | 5.87 (m) | 6.03 (t; 8.5) | 5.53 (t; 7.5) |
| 14a | 1.06 (dd; 7.5, 14.5) | 1.60 (dd; 7.1, 14.8) | 1.22 (dd; 7.7, 13.9) |
| 14b | 2.69 (dt; 14.6, 9.8) | 2.53 (dd; 9.8, 14.8) | 2.49 (dd; 7.3, 13.9) |
| 15 | -- | -- | -- |
| 16 | 1.11 (s) | 1.2 (s) | 1.32 (s) |
| 17 | 1.62 (s) | 1.62 (s) | 1.15 (s) |
| 18 | 2.11 (s) | 2.10 (s) | 1.83 (s) |
| 19 | 0.75 (s) | 0.76 (s) | 0.78 (s) |
| 20a-(E) | 4.85 (s) | 4.92 (s) | 4.76 (s) |
| 20b-(Z) | 5.21 (s) | 5.25 (s) | 5.17 (s) |
| -OCOMe | 2.01 (s) | 2.05 (s) | 2.00 (s) |
| -OCOMe | 2.05 (s) | 2.02 (s) | 2.01 (s) |
| -OCOMe | 2.06 (s) | 2.08 (s) | 2.02 (s) |
| -OCOMe | 2.16 (s) | 2.16 (s) | 2.04 (s) |
| -OH | -- | 1.42 (s) | 2.42 (s) |

**Table S7. Sequences for proteins identified in this manuscript**

| Abbreviation | Full name | Gene ID from *T. chinensis* | Sequences |
|---|---|---|---|
| FoTO1 | facilitator of taxane oxidation | ctg24251_gene.1 | **MAETMNEKVGADEKLIEDSRNTGQEENDYSKRLGPLSRNIIPHLINIYTCIATPRDLEIYHPDATFEDFFVRAFGIKEIKSIHYSFPKLVYDGKILEYSVEENETSPGCGELLFNIKQQYKVLYLGKEVNITTLMVLQIENGKIIKHEDRINQHPVRGRHDISVPLVGRAREGIRRLTMLMLHVRMGFGKDPTPP** |
| T9αH-750C | taxane 9-alpha hydroxylase | ctg10747_gene.1 | **MAFSRLIEAAAAEAPTIVTVLLLSFIFYLWRRNSSSSSTRLPPGPFQWPVIGNLHQFGRLPHLSIQHLANKYGPIMWLRLGYYPVVVVSTTEMAKEFLKIHDLAFSSRPKSGVGEHLVYNYKSMGFSPYGDYWRHIRRVWMTELMTPKRLSSFRSIREEEVCSAMRSIWEKSEQGRVAVNVSKAIDWISSSIVWRTVAGKKCSEDRDGKDLCDMVKRLMLTVKEVNGREIIPCIGWFDLQGVTRRMKETHRIFDGVAQNIIDQHINGRKREQSSDVKDIVDVLLEMADTGAINIQLDSIKAIIFDVLTGGIETATSSLEWTMTEMVRNPDIARKLQQEIESVVGKHRTVTESDLPNIEYLQCVVQESLRLHPPAPLIFPRESTEACTVGAEGYVIPPKTRLMINVWAIGRDPAVWEDSLTFKPERFMGKDMDIKGRSDFRMLPFGGGRRGCPGAQMAIGNMELILAQLMHCFDWRAEGDPSELDMSEALGTSLSRKHNLFAVPTLKLLNCI** |
| T7AT | taxane 7β-*O*-acetyltransferase | ctg3030_gene.2 | **MENPSSTDFLVKKFDPVVVAPSLPLPKTTLQLSPIDNQIGFRGFFNSLSVYNAPDDISADPVKIIREALSKVLVHYFPLAGRFRNKENGELEVDCTGEGALFVEAMVEDNISVLRDFDDLNPSFQQLVFWPPMGANIEDLHLLVVQVTRFTCGDITIGVTVCHSIFDGCGAAQFVTALADMARGEVKPLLEPIWNRELLKPEDPLHLQLYQFDSLCPPPILFEELGQASLIINSNTIKYMKQCIMEECKVFCSTFEVMAALVWVARTKAFQIPHTETVKLLFAMDMRRSFNPPFPNGYYGNAIGTAYAMDNVEDLLNGSLSRVVMIIKKSKVSLRDNYLRSNKVKDPYSLDVNKKDNNVLALSDWRRLGFHEANFGWGDPVNVTAPQLLGKGLPLLSYYLFLQPSKNQPDGIKILMSCMHPSAVKSIKMEMEAMINKFVNKS** |
| T1βH-686 | taxane 1-beta hydroxylase 1 | ctg5594_gene.1 | **MASSAQNGVELSYIDLSQFSFDSEGLKNLQNHPGVATVREACKEFGCFGVLNTGIPDDVVQKLESVSHELCAMPSEMKDRAITSNPYDSYNRIPYRESFWFPTTWDSDSVLAYFNKLWPEKDNLNLCETVMAYALGMAELQRKISFIIIASLGLDVETFYHSDFEKATSYMRVHHHYSEGKFAAGEEALFGHYDPNCFTMLYQDIGGGLQIESKEGKWVDAKPGSLVINVAESLKAWSNGRYSSAKHRVVYKDWMHRLSVGWVMQFPDKEICAPAELVDEQHPQLYRPFPYPPFLDASMKYRITIDKYAGISPIY** |

| | | | |
|---|---|---|---|
| T9dA (DeAc898) | taxane C9 deacetylase | ctg19840_gene.3 | **MEAATAEPRVMQDMHGFIKVYSDGSVVRAGEPYFPAAISEENNDKLGPYKDVVYNAELGLWARIYLPPPPHKKTRLPVLLFFHASGFCILSPATPVVHRLCLLWAAKTGVIIVSVKYRLAPEHRLPAAYDDSIAALQWLLAMKSTEPGAVAVDPWLHSHADLSNIFVAGESAGGNIAHYLGCWVAAQDGEIQAQVKGLILVCPFFGGEDRTPSEEGNLAVMSEADIIWKYALPVGSNRDHPFCNPVGEGRESAISSLALPPILFVIAGLDVLRDKELQYCELLKKCGKQLEVVMFDEENHGFTLFNAEDQKSLEVIRCISDFVWSKS** |
| T9ox | taxane C9 oxidase | ctg4166_gene.5 | **MEQNNMRNEVDLPFIDLSQFSFDSEGIKNLQNHPGVATVMESCQEWGFFRIMNTGIPNDVFQKVESVSHELFAMPQEMKDRAITSSPHDTYINHPYRESFWFPTPPHSDSVLAFCNKLWPEKDNLKLCETIGTYILSMEDLKRKISSIIIASLGLDKVETFYHSDFENGTSVFRIHHYYSDGKFAAGEEALFAHTDPHCFTILYQDNGGGLQIQSKEGNWVHVKNIPNSLIINVADSLKAWSNGRYRSVNHRVVYKDWTNRISLGWFMMFPDKEIRVPAEFIDDQHPQCYRPFTYLQFRDAFMKDRIDIDGYAGIIPTY** |
| T7dA (DeAc1023) | taxane C7 deacetylase | ctg1975_gene.2 | **MADNSEPRVVENLYDGVIKLYSDGSIVRGDQQSPPPPTDDYNCVPFKDIVFDHTLGLWARVYLPPQTAKTRVPVLVYYHGGGFCCEFPPSTAILDCMCHKWAATLGVIIVSAEYRLAPEHRLPAAYHDAISALHWIDSMKSGAVEVDPWFRSHADFCKVFVAGDSAGGNIANHVGIWAAGAHGDGDLQIQIKGIILGCPFFGGEERTPSGSHNSPVFNLEISDTMWRLSLPLGSNRDHPFCNPVGVGDLKEADLPPMLFVIAGQDILKDKQLQYCEFLKGCGKQVEVHVFEEEDHGFTALKMENRSAVEALRCISHFINLTN** |
| PAL | phenylalanine-CoA ligase | ctg867_gene.5 | **MDAEVVKSVRELGVDDVVQAGLPRHRAEIFYGQLQRAIADIGGSQTSLWHRVSKELLAPHHPHALHQLMYYSIYKNWDTSENGPPLYWFPTQESARLTNLGWMMETYGPQLLGSSYYNPITSFQSFQQFTVDHPEVYWSLVLKELSVVFHESPRCILDTSDKSRSGGVWLPGSVLNVAESCLSAKESINKTDNSIAIVWREEGRNEYPVNKMTLGELRAKVMRIANALDVVFTKGDAIAIDMPMTVNAVAIYLALILAGYVVVSIADSFVPKEIATRVRVTKAKGIFTQDFILRGGKRIPLYSRVVESGAPKAIVIPAEEELGTQLREIDVAWSKFLSFSDHLRSPEYYSAVRQPVDARTNILFSSGTSGEPKAIPWTHSPPIRCGSECWSHLDVKAGDIFCWPTNLGWVMGPVLVYSCFLSGATAAIYEGSPLDRGFGKFVQDARVTVLGTVPSMVKTWKSTGCMEGLDWSHIRTFASTGEASSIDDDLWLSSKGWYKPVIELCGGTELSACFVHGSLLQPQALGMFSTPTMTTGFVLFDDQQIPYPNDQPCIGEIGLFPRFFGSSYTLLNADHDAVYFKGMPMYKGMRLRRHGDMIERTVGGYYKAHGRSDDTMNLGGIKTSAIEIERVCNRAHEQVLETAAISISSSEGGPELLAILTVLKDGPTVSMDTLKLAFSKAIQSNL** |

| | | | |
|---|---|---|---|
| | | | **NPLFKVSFVKVISDFPRTASNKIMRRVLRDQIKQEFSLHKSRL** |
| T1βH-184 | taxane 1-beta hydroxylase 2 | ctg13625_gene.1 | **MASSLQNEDDLPIISLSQFSFESEGLKNLQNYPGLAKVREACKKWGFFRIVNTGIPNDVFRKMESVSHELCVMPQEMKDRAITSDPYDSYSQTPSRESFWFPTPSHSVSVQDFCNKLWPEKDNLKLCQTIGAYMFGMKELQRKISVIILASLGLDLETFYHSDFEKGTSIFRIHHHYSDGKFAVGEEALFGHTDPNCFTILYQDNGGGLQIQSKEGNWVDVKPVPNSLVINIADSLKAWSNGRYRSAKHRVVYKDWTNRISFVWMLMFPDKEIRAPTELIDEQHPQHYRPFTYHPFREATMKDHVNIDGYAGIFPTY** |

**Table S8. Experimental setup for compounds isolated in this study.**

| Compound | Genes | Plants | Yield | Column condition | NMR |
|---|---|---|---|---|---|
| taxusin (**6**) | tHMGR, GGPPS, tTDS1, tTDS2, FoTO, T5αH, TAT, T10βH, DBAT, T13αH, T9αH-750C, TAX19, CJM236* | 12 x 5-week old plants (10.94 g DW) | 4.1 mg (375 μg/g DW) | 1. 100 g silica column (EA:Hex = 1:4, 2 L) 2. Biotage C18 6 g column (45% 2 CV, 45-49% over 5 CV, 49% 2 CV, 49-52% over 6 CV, 52-55% over 10 CV) | **Table S3** |
| 13β-taxusin (**6'**) | | | <1 mg (<91 μg/g DW) | | **Table S3** |
| 1β-hydroxytaxusin (**6-O1**) | tHMGR, GGPPS, tTDS1, tTDS2, FoTO, T5αH, TAT, T10βH, DBAT, T13αH, T9αH-750C, TAX19, T1βH-184 | 8 x 4-week old plants + 6 x 5-week old plants (12.25 g DW) | 2.2 mg (180 μg/g DW) | 1. 100 g silica column (EA:Hex = 1:4, 1.5 L; EA:Hex = 1:3, 0.5 L; EA:Hex = 2:3, 0.5 L; EA:Hex = 3:2, 0.5 L) 2. Biotage C18 6 g column (40% 5CV, 40-45% over 15 CV) | **Table S4** |
| 15-hydroxy-11(15→1)*abeo*-taxusin (**6-O2**) | | | 0.4 mg (33 μg/g DW) | | **Table S5** |
| baccatin III (**16**) | HMGR, tGGPPS, TDS, FoTO, TAT, T10βH, DBAT, T13αH, T9αH-CYP750C, T2αH, TBT, T7βH, T7AT, TOT, T1βH-686, DeAc898, DeAc1023, and T9ox (Note: exclude T5αH) | 53 x 4-week old plants (30.70 g DW) | 1.33 mg (Partially pure; 43 μg/g DW) | 1. 100 g silica column (EA:Hex = 1:1, 1 L; EA:Hex = 3:2, 0.5 L; EA:Hex = 7:3, 0.5 L; EA:Hex = 4:1, 0.5 L; EA:Hex = 9:1, 0.3 L; EA, 0.5 L) 2. Biotage C18 6 g column (30% 2 CV, 30-32% over 2 CV, 32-37% over 10 CV) | **Fig. S22** |

Solvent system for biotage Sfär C18 D Duo 100 Å 30 μm column: A =water, B = acetonitrile. Percentage of solvent B is listed. DW: dry weight, EA: ethyl acetate, Hex: hexane, CV: column volume.

*CJM236 is a non-functional gene accidentally included in this expression. It does not affect the accumulation of taxusin (**6**) and 13β-taxusin (**6'**).

**Table S9. Oligonucleotide primers used in this study.**

| | | |
|---|---|---|
| **Agrobacterium vector (pEAQ) cloning primers** | | |
| | FoTO1 fwd | atattctgcccaaattcgcgaccggtATGGCAGAAACAATGAATGAGAAAGTTGGCG |
| | FoTO1 rev | taatgaaaccagagttaaaggcctcgagTCATGGAGGAGTTGGATCCTTTCCAAAGC |
| | T9αH-750C fwd | ttctgcccaaattcgcgaccggtATGGCATTTTCTAGACTGATTGAAGCAGC |
| | T9αH-750C rev | gagttaaaggcctcgagTTAGATGCAATTTAACAATTTCAGGGTAGGAACTG |
| | T7AT fwd | atattctgcccaaattcgcgaccggtATGGAGAATCCAAGCTCAACAGACTTCCT |
| | T7AT rev | agagttaaaggcctcgagTCATGATTTATTCACAAATTTGTTTATCATGGCTTCC |
| | T1βH-184 fwd | aagcttctgtatattctgcccaaattcgcgaccggtATGGCTTCCTCGCTGCAGAATGAA |
| | T1βH-184 rev | taatgaaaccagagttaaaggcctcgagTCAATAAGTAGGAAATATGCCTGCGTAACCA T |
| | T1βH-686 fwd | tctgtatattctgcccaaattcgcgaccggtATGGCTTCCTCGGCGCAGAAT |
| | T1βH-686 rev | aaaccagagttaaaggcctcgagTCAATAAATAGGGGATATGCCGGCGTATT |
| | T9ox fwd | tattctgcccaaattcgcgaccggtATGGAACAGAATAACATGCGAAATGAAGTTGATCT |
| | T9ox rev | atgaaaccagagttaaaggcctcgagTCAATAAGTAGGAATTATGCCTGCATAACCGTC A |
| | T9dA fwd | agcttctgtatattctgcccaaattcgcgaccggtATGGAAGCAGCTACGGCGGAGC |
| | T9dA rev | aaaccagagttaaaggcctcgagCTAACTTTTACTCCAGACGAAATCAGAGATGCAG |
| | T7dA fwd | agcttctgtatattctgcccaaattcgcgaccggtATGGCGGATAACAGCGAGCCTC |
| | T7dA rev | ccagagttaaaggcctcgagTTAATTAGTTAAGTTAATGAAGTGAGAGATGCAACGG |
| **FoTO1 mutants** | | |
| | C49T fwd | CCTTATCAATATATATTGTTGCATAGCAACACCAC |
| | C49T rev | GTGGTGTTGCTATGCAACAATATATATTGATAAGG |
| | F83H fwd | GAGATCAAGTCAATCTTTTATTCCTTTCCAAAG |
| | F83H rev | CTTTGGAAAGGAATAAAAGATTGACTTGATCTC |
| | A61H-N63D fwd | GACCTTGAAATATATGCTCCGAATGCGACATTTGA |
| | A61H-N63D rev | TCAAATGTCGCATTCGGAGCATATATTTCAAGGTC |
| | F57L fwd | GCAACACCACGTGACTTTGAAATATATCATCC |
| | F57L rev | GGATGATATATTTCAAAGTCACGTGGTGTTGC |
| | I114F fwd | GCGGAGAGTTACTGATTAACATCAAGCAACAG |
| | I114F rev | CTGTTGCTTGATGTTAATCAGTAACTCTCCGC |
| | Truncate N-term | ctgtatattctgcccaaattcgcgaccggtATGAGATTAGGACCGCTTTCACGCAATATAAT TCCTC |

| | | |
|---|---|---|
| | Truncate C-term | aatgaaaccagagttaaaggcctcgagCTAGTTAATCCGGTCTTCGTGCTTGATGATCTTCCC |
| **Tagged constructs** | | |
| | V5 Tag - C terminus fwd | TAGAATCAAGTCCAAGAAGTGGATTTGGAATTGGCTTTCCTCCAGAGCCTGGAGGAGTTG |
| | V5 Tag - C terminus rev | atgaaaccagagttaaaggcctcgagCTAAGTAGAATCAAGTCCAAGAAGTGGATTTGG |
| | V5 Tag - N Terminus fwd | GCCAATTCCAAATCCACTTCTTGGACTTGATTCTACTggctctggaGCAGAAACAATGAA |
| | V5 Tag - N Terminus rev | atattctgcccaaattcgcgaccggtatgGGAAAGCCAATTCCAAATCCACTTC |
| | HA-T5aH fwd | GAAACCAGAGTTAAAGGCCTCGAGTTAAGCATAATCTGGAACATCATATGGATAtccagagcC |
| | HA-T5aH rev | TTAAGCATAATCTGGAACATCATATGGATAtccagagcC TGGTCTCGGAAACAGTTTAAT |
| **E. coli expression vector (pET28a) cloning primers** | | |
| | pET fwd | TAGctcgagcaccaccaccaccaccactgagatccgg |
| | pET rev | atggctgccgcgcggcaccaggccgctgctgtgatgatgatgatgatggctgctgcccat |
| | 3xFLAG-TEV fwd | ggcctggtgccgcgcggcagccatgactataaagatgatgacgacaaaGACTATAAGGAC |
| | 3xFLAG-TEV rev | AGACTGAAAGTATAAATTCTCtttatcgtcgtcatccttgtaatcCTTATCATCGTCGT |
| | T5αH fwd | gacgataaagagaatttatactttcagtctTCCTCCCTTAAACTTCCTCCTGGGAAATTAG |
| | T5αH rev | tctcagtggtggtggtggtggtgctcgagCTATGGTCTCGGAAACAGTTTAATGGAAAAT |
| | TDS2 fwd | tgacgacgataaagagaatttatactttcagtctagcggtagcccgaccaagttggcta |
| | TDS2 rev | tggtggtggtggtggtgctcgagCTAtacttgaatcggttcaatgtaaacttttcttata |
| | mTurq-FoTO1 fwd | acgataaaGAGAATTTATACTTTCAGTCTGTGAGTAAGGGTGAAGAATTATTTACTGGCG |
| | mTurq-FoTO1 rev | accgctcatgcttaatttctcctctttaattCTATGGAGGAGTTGGATCCTTTCCAAAGC |
| | mTurq-FoTO1Δ Cterm rev | tcagtggtggtggtggtggtgctcgagTCAGTGTTGGTTAATCCGGTCTTCGTGC |
| **Others** | | |
| FoTO1-mTurq2 | FoTO1 rev | AAATAATTCTTCACCCTTACTCACtccagagccTGGAGGAGTTGGATCCTTTCCAAAGCC |
| | C-term mTurq2 fwd | ggctctggaGTGAGTAAGGGTGAAGAA |

| | | |
|---|---|---|
| | C-term mTurq2 rev | ttaatgaaaccagagttaaaggcctcgagCTATTTATACAACTCATCCATTCCGAGCGTGATG |
| mTurq2-FoTO1 | N-term mTurq2 fwd | ctgtatattctgcccaaattcgcgaccggtATGGTGAGTAAGGGTGAAGAATTATTTACTGGC |
| | N-term mTurq2 rev | tccagagccTTTATACAACTCATCCATTCCGAGCGTGATGC |
| | FoTO1 fwd | GGAATGGATGAGTTGTATAAAggctctggaGCAGAAACAATGAATGAGAAAGTTGGCGCT |
| T5αH-mCherry | T5αH-mCherry rev | TCGCCCTTGCTCACCATGCTTCCCGAACCTGGTCTCGGAAACAGTTTAATGGAAAATCCC |
| | T5αH-mCherry fwd | CCATTAAACTGTTTCCGAGACCAGGTTCGGGAAGCATGGTGAGCAAGGGCGAGGAGG |

#### Supplemental Note 1

We isolated a fungal species from our *Taxus* needles and included it as a component of our elicitation. We amplified the ITS2 sequence and found it matched *Aspergillus tubingensis* NRRL 4875. *A. tubingensis* and other *Aspergillus ssp.* have been previously identified from *Taxus* tissues [19].

ITS2 sequence:
TGAATCATCGAGTCTTTGAACGCACATTGCGCCCCCTGGTATTCCGGGGGGCATGCCTGTC
CGAGCGTCATTGCTGCCCTCAAGCCCGGCTTGTGTGTTGGGTCGCCGTCCCCCTCTCCGG
GGGGACGGGCCCGAAAGGCAGCGGCGGCACCGCGTCCGATCCTCGAGCGTATGGGGCTT
TGTCACATGCTCTGTAGGATTGGCCGGCGCCTGCCGACGTTTTCCAACCATTTTTTCCAGG
TTGACCTCGGATCAGGTAGGGATACCCGCTGAACTTAAGCATATCAANANNCGGAGGAAGA
CAAGATCGGTCTGGCCGCCACCGCCTCCACCACCTATGGCGAGCCCTATCACACCGCCCG
CGCCTTCGCGTCCATCGACCATCTGAGCCACGGCCGCGCGGCGTGGAACATCGTCACCAC
CTCCTATGCCCGCACGGCGGCGAATTTTTCCAAGAGCCATCCCGAGCATGACGAGCGTTAT
GCCGTGGCGGAAGAATATGTAAACGTCGTGCGCGGCCTGTGGGACAGCTGGGACGACGAT
GCCTTTGTCAAGGACAAGCAGGCTGGGCGTTATGTCGATCCGGAAAAGGTCCATATCCTCG
ACCACGAAGGCAAATATTTCNCGGTCAAAGGCCCACTCAACATTCCGCGCTCGCCGCAGGT
TCACCTACGGAA

ITS primers:

| | |
|---|---|
| fwd ITS1 | TCCGTAGGTGAACCTGCGG |
| fwd ITS2 | GCTGCGTTCTTCCATCGATGC |
| fwd ITS5 | GGAAGTAAAAGTCGTAACAAG |
| rev ITS3 | GCATCGATGAAGAACGCAGC |
| rev ITS4 | TCCTCCGCTTATTGATATGC |

### Supplementary Figures

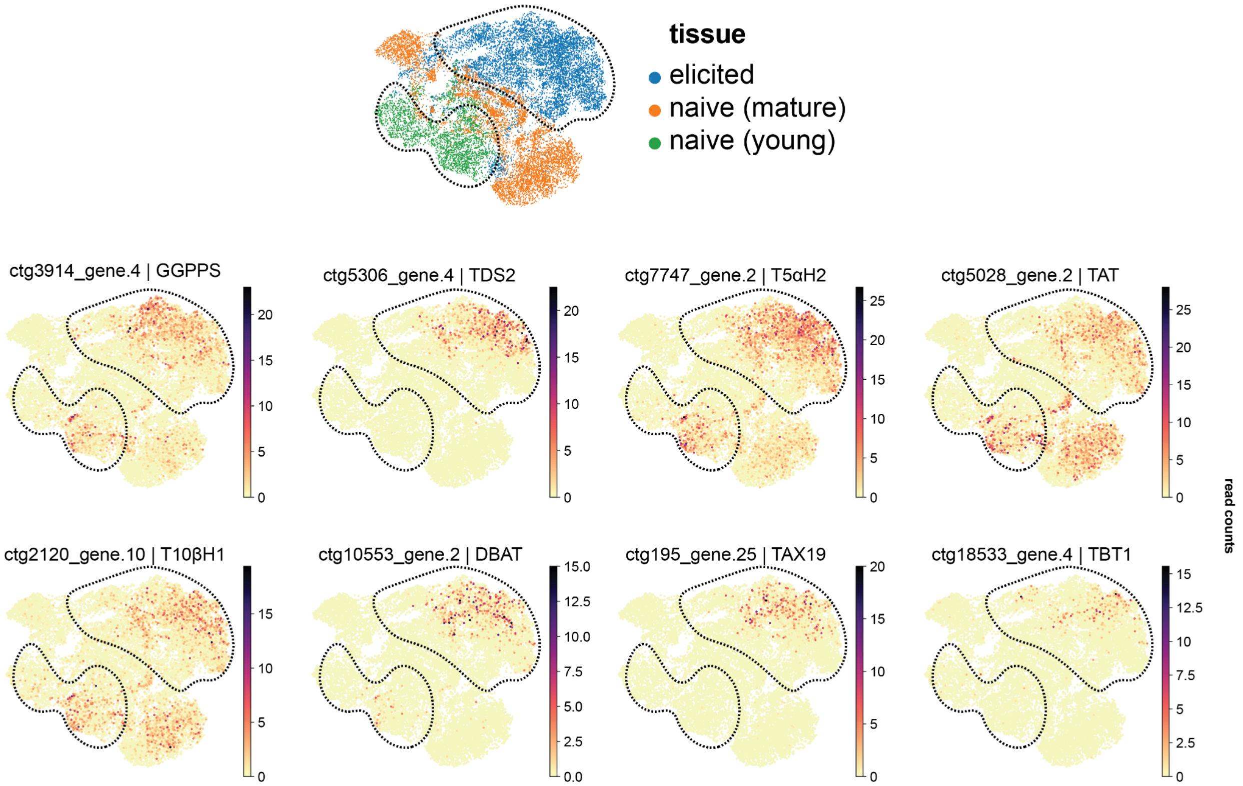


**Fig. S1.** UMAP[20] of Taxol biosynthetic enzyme expression across single cells from naive *Taxus media* tissues and elicited *T. media* tissues. Three separate single-cell transcriptomic experiments were integrated (cellbender[21], scvi[22], including mature naive aerial tissues (orange), new needle and bud growth (green), and needles from the multiplexed perturbation (blue). Dotted black line encircles the elicited (top right) and young tissue (bottom left) cells in each plot.

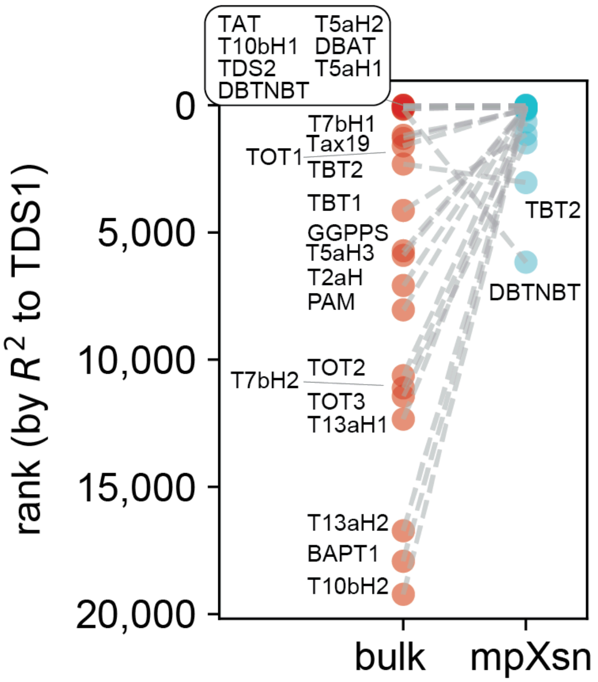


**Fig. S2. Annotated correlation rank plot of known Taxol biosynthetic genes (with bulk RNA-seq vs mpXsn)**. Genes in the *T. chinensis* genome are ranked by their Pearson correlation to TDS1 using either 79 bulk RNA-seq samples[23–28] or mpXsn data. The ranks of known Taxol biosynthetic enzymes and GGPPS are indicated in this plot.

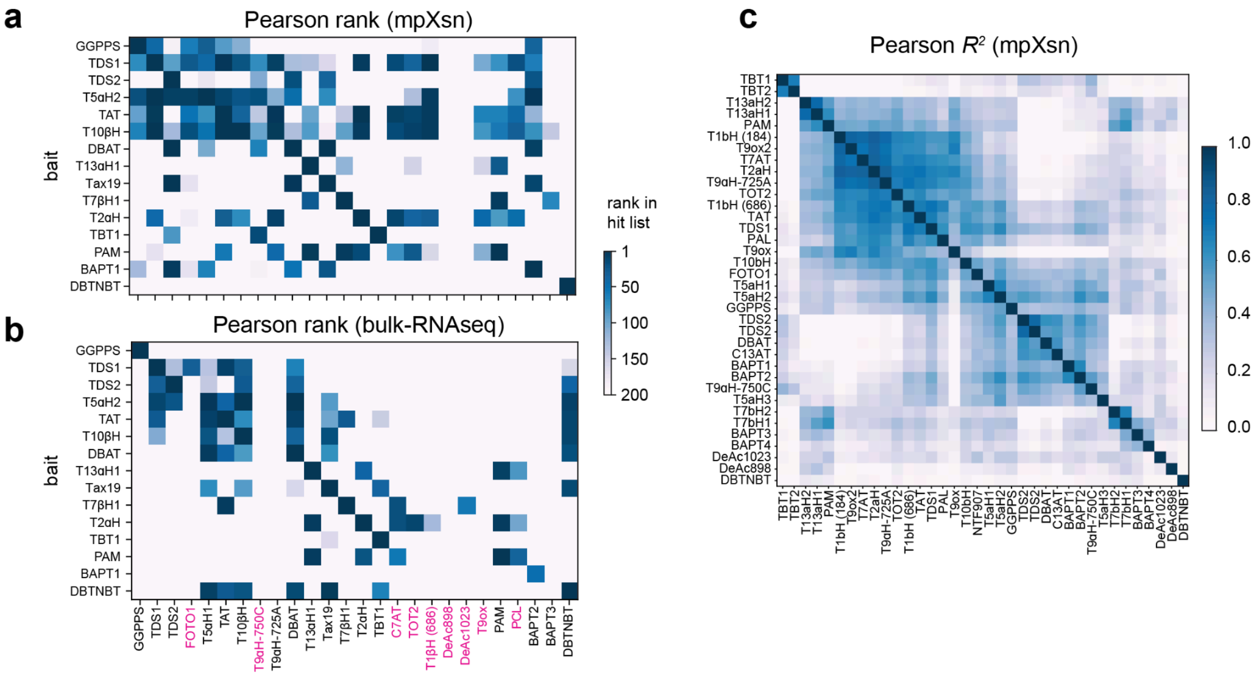


**Figure S3. Pearson correlation rank and scores between Taxol enzymes**. Using either mpXsn (**a**) or bulk RNA-seq (**b**) data, Pearson correlation ranks were calculated to all genes in this study (x axis) using the previously known Taxol biosynthetic enzymes as bait (y axis). (**c**) Pearson correlations between all Taxol genes in this study, using the mpXsn data. Genes are hierarchically ranked on both axes.

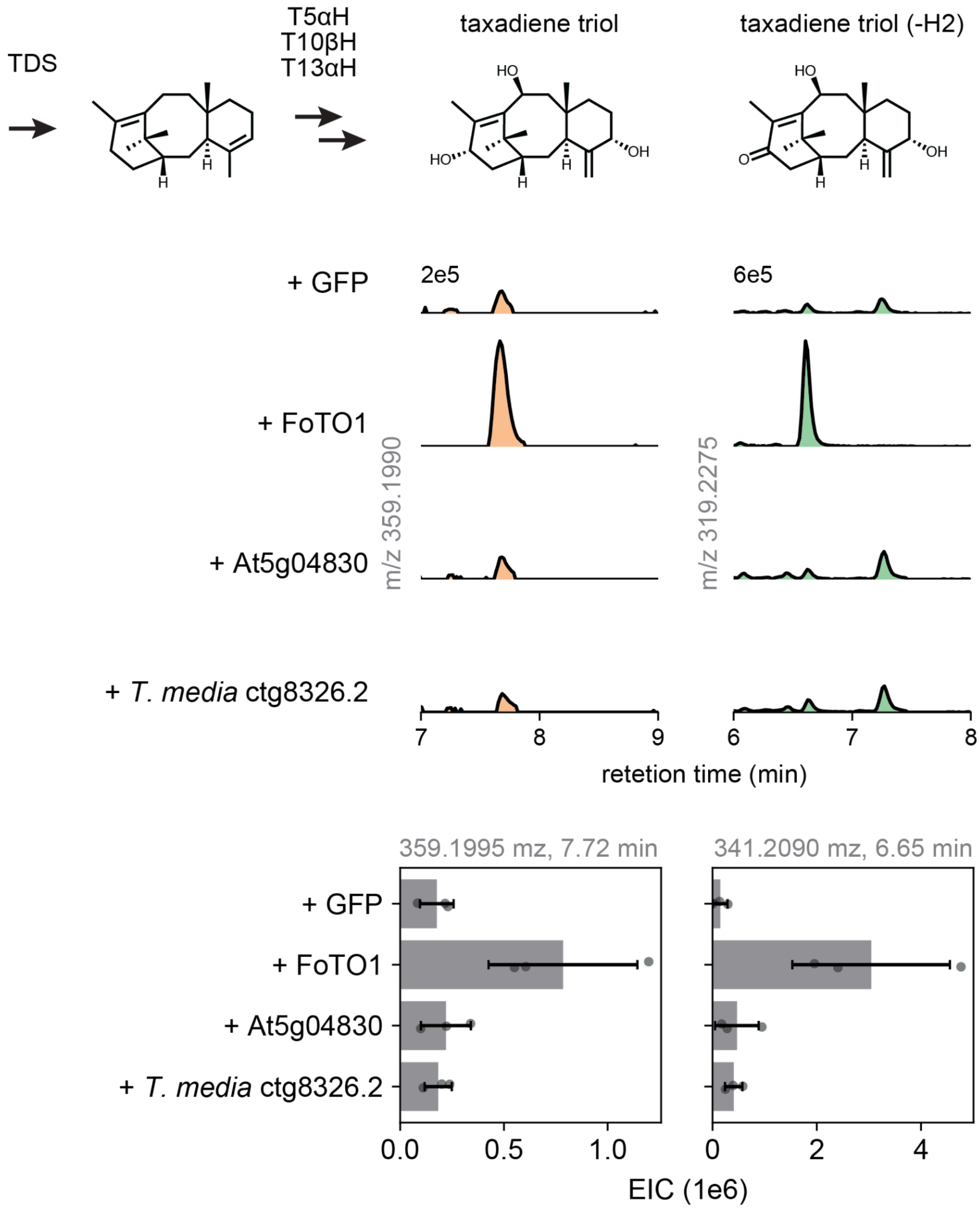


**Figure S4. FoTO1 homologs from *T. media* and *A. thaliana* have no effect on taxane yield increase**. TDS2 and the first three pathway oxidases were coexpressed in *N. benthamiana* with either GFP, FoTO1, At5g04830 (*Arabidopsis thaliana* homolog), or ctg8326.3 (*Taxus media* homolog). Masses corresponding to the expected products, 5α,10β,13α-triol and 5α,10β-diol 13-one, are displayed as extracted ion chromatograms (EICs) and bar graph quantified across three independent replicate leaves. Unlike FoTO1, neither FoTO1 homolog alters the product profile of this early subpathway.

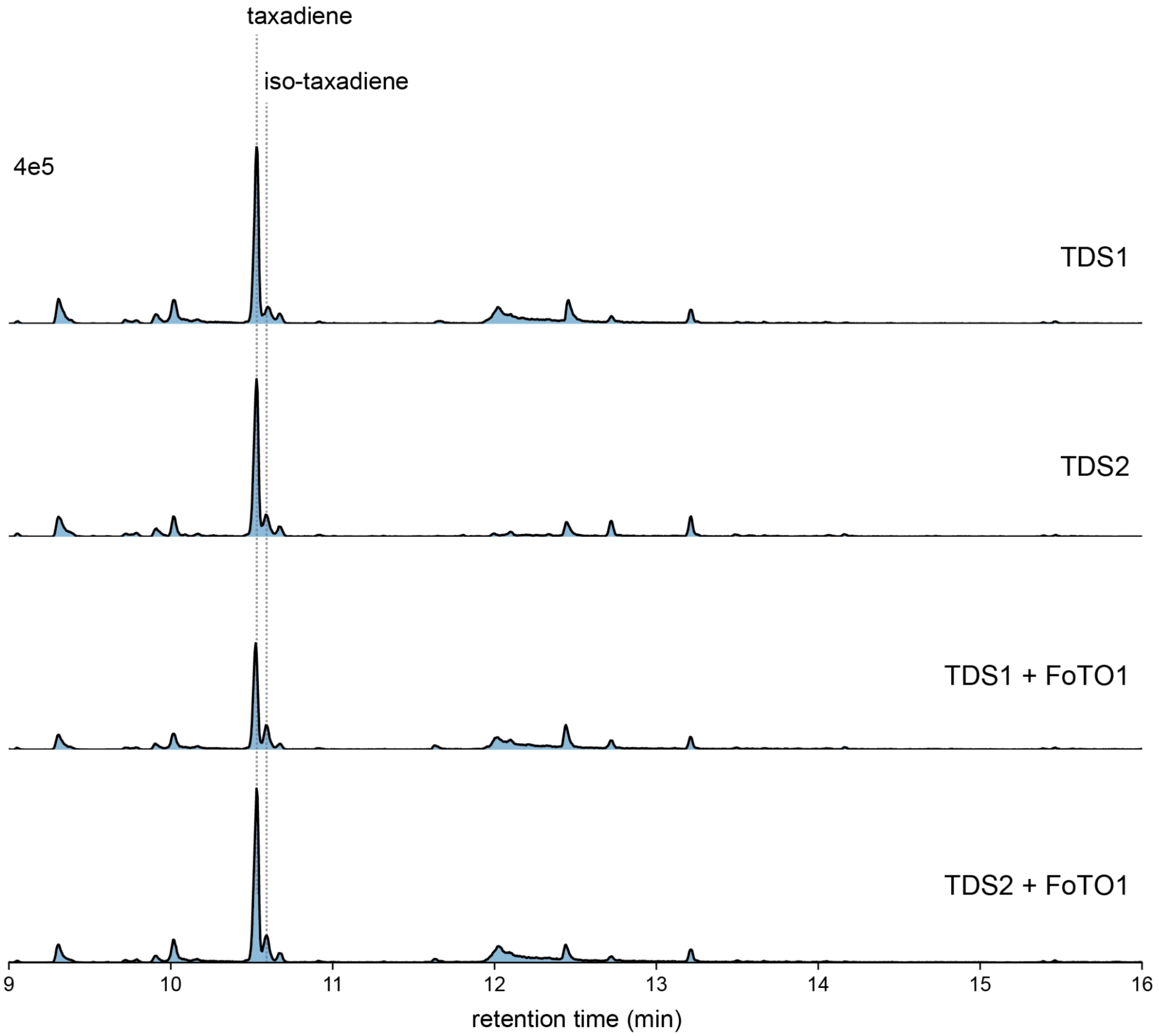


**Figure S5. FoTO1 does not affect the production of taxadiene (1) by TDS**. GCMS total ion content (TIC) traces of *N. benthamiana* expressing boost (tHMGR, GGPPS) and taxadiene synthase (TDS1 or TDS2) with and without FoTO1. When FoTO1 is coexpressed, the formations of taxadiene [**1**, taxa-4(5),11(12)-diene] or iso-taxadiene [taxa-4(20),11(12)-diene] remain the same and there are no observed new peaks. Representative traces of three biological replicates for each condition are shown.

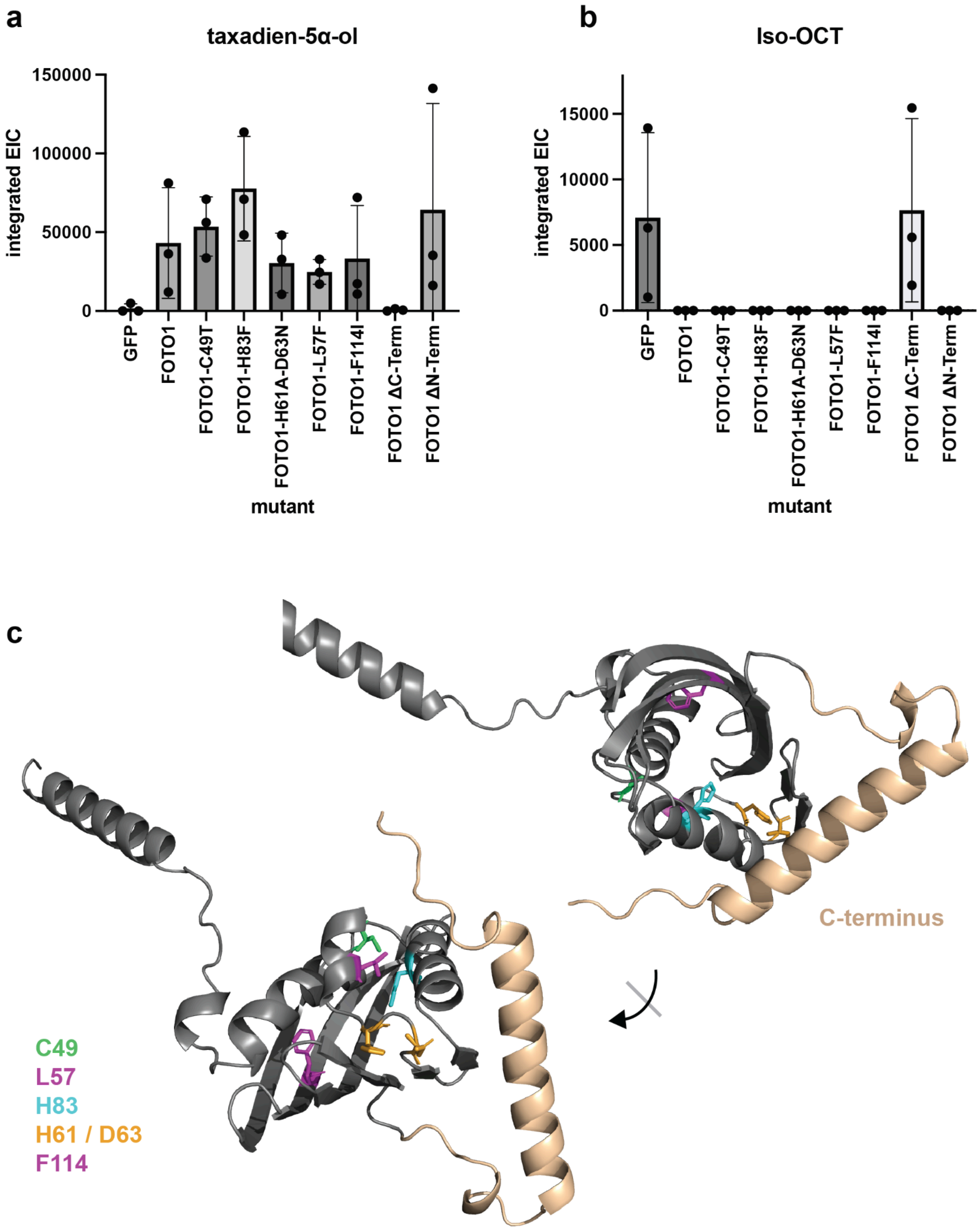


**Figure S6. Substitution and truncation mutants of FoTO1**. (**a-b**) Relative yields of taxadien-5α-ol (**2**, desired product) and iso-OCT (**2'b**, an undesired product) measured by GCMS when TDS2 and T5αH were coexpressed with various FoTO1 mutants in *N. benthamiana*. Candidate residues for catalytic or substrate interaction in the cavity of the NTF2-like fold and regions of truncation were selected based on the AlphaFold3 structure. (**c**) FoTO1 structure predicted by AlphaFold3[29].

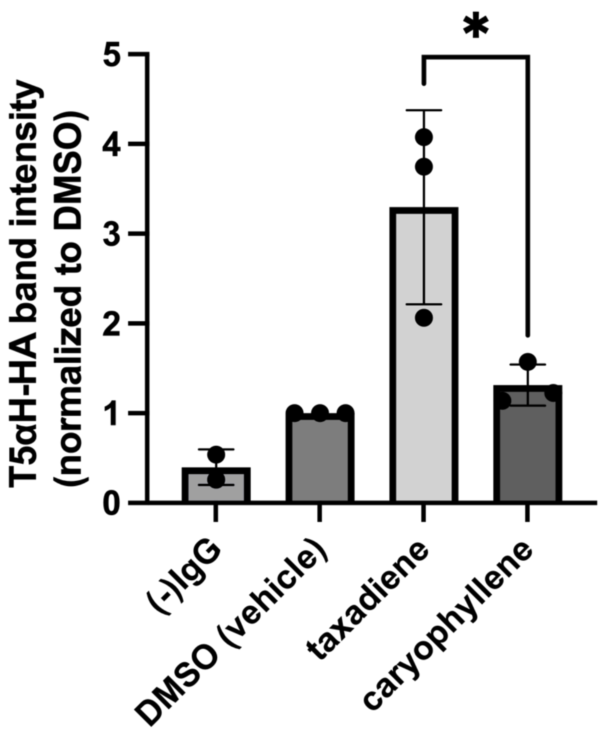


**Fig. S7. Quantification of co-immunoprecipitation T5αH by FoTO1**. V5-FoTO1 and T5αH-HA were coexpressed in *N. benthamiana* leaves for five days. Lysates were treated for 15 minutes with 400 µM taxadiene (**1**), caryophyllene or a DMSO vehicle prior to immunoprecipitation and quantification of T5αH-HA by western blot. Caryophyllene is a sesquiterpene with hydrophobicity similar to taxadiene. (-)IgG indicates immunoprecipitation without antibody, as a control for nonspecific binding to beads. Band intensities were normalized to the DMSO vehicle condition and p-values were calculated by two-tailed t-test. Three dots indicate independent replicates conducted on different days.

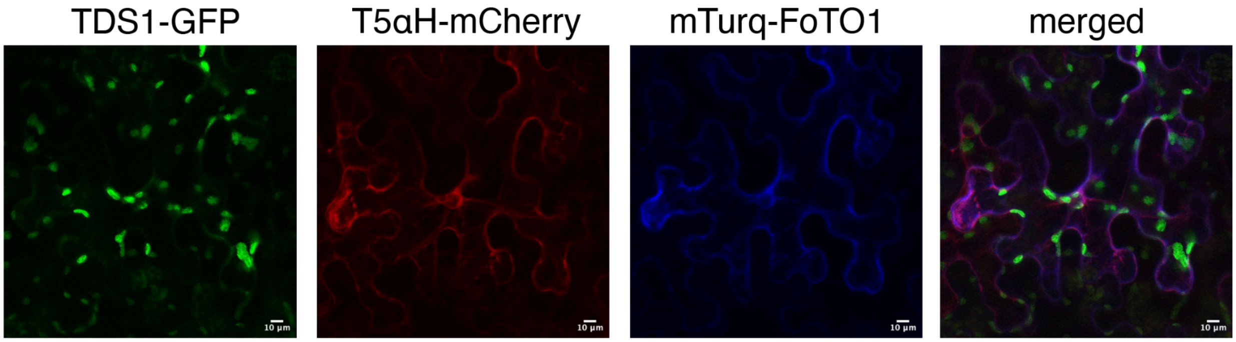


**Fig. S8. TDS1, T5αH, FoTO1 show different subcellular localizations in *Nicotiana benthamiana***. TDS1, T5αH, and FoTO1 were fused with fluorescent protein GFP, mCherry, and mTurq, respectively, and the fusion proteins were expressed in *N. benthamiana* using *Agrobacterium*-mediated expression. Leaves were harvested at 5 day post infiltration and imaged by Leica SP8 confocal microscopy. Fluorescence from GFP (green), mCherry (red), and mTurq (blue) are shown. Scale bars indicate 10 µm.

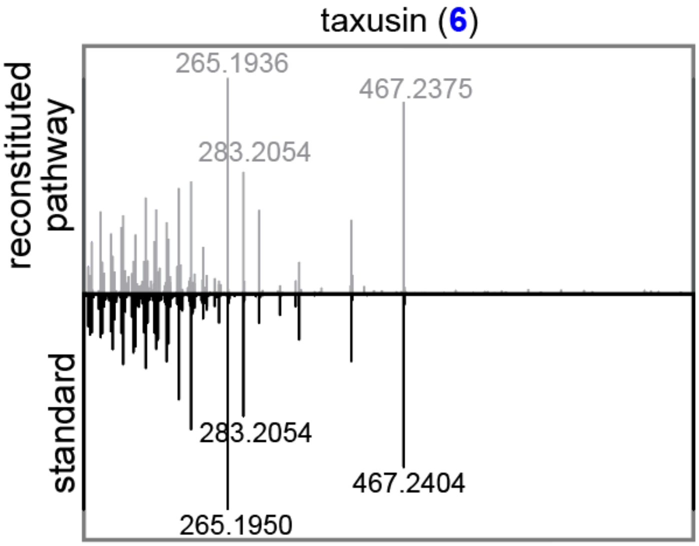


**Fig. S9. MSMS fragmentation patterns of heterologously produced taxusin (6) in *N. benthamiana* compared to that of taxsuin (6) standard**. MSMS fragmentations were generated using $[M+Na]^+$ (m/z = 527.2621) as the precursor ion and fragmented with a collision energy of 30 eV. Regions between m/z 100 to 800 are shown. The following gene set was heterologously expressed in *N. benthamiana* via *Agrobacterium*-mediated transient expression: HMGR, tGGPPS, TDS, FoTO, T5αH, TAT, T10βH, DBAT, T13αH, T9αH-CYP750C, and TAX19.

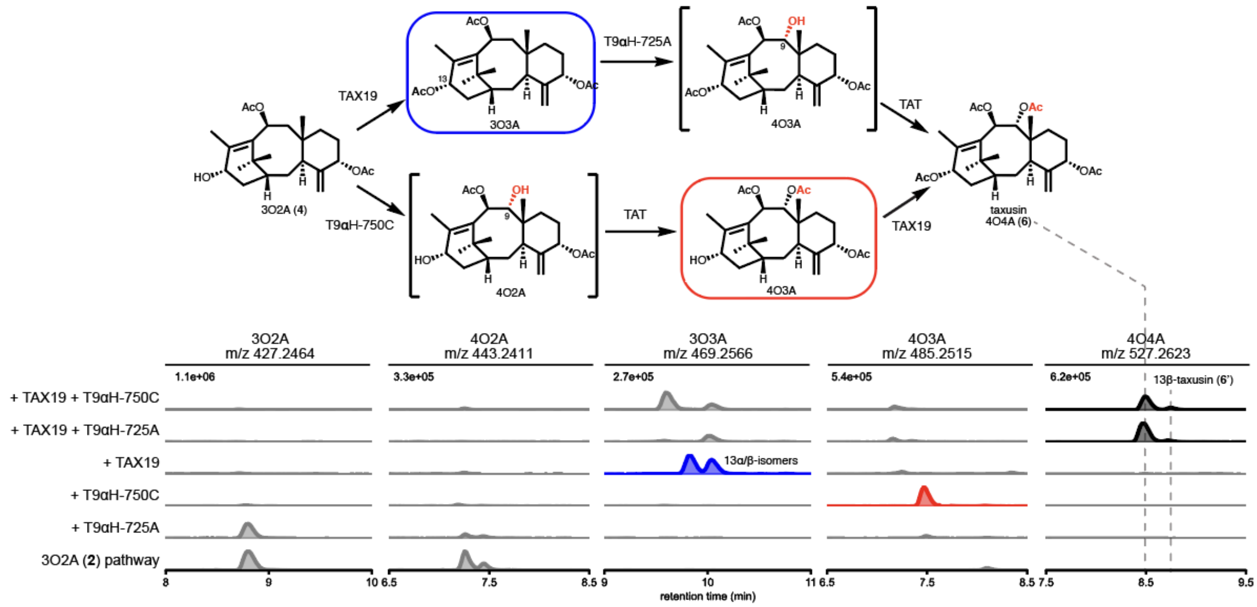


**Fig. S10. T9αH-725A and T9αH-750C utilize different biosynthetic routes to yield taxusin (6).** Combinations of genes, indicated on the left, are expressed in *N. benthamiana* via *Agrobacterium*-mediated infiltration, and the EICs of the corresponding intermediates are shown. Expression of T9αH-725A with the 3O2A (**2**) pathway did not yield any new product, while expression of T9αH-750C resulted in a 4O3A (red). 4O2A was proposed to be the intermediate; however, at the presence of TAT (within the early 3O2A pathway), it was quickly turned over to 4O3A and was not detectable. See **Fig. S11** for the bifunctional role of TAT. TAX19 produced two major products which we proposed to be the C-13α/β isomers (blue) that eventually resulted in taxusin (**6**) and 13β-taxusin (**6'**) in TAX19 + T9αH-725A and TAX19 + T9αH-750C conditions. As T9αH-725A did not affect 3O2A (**4**) accumulation but was able to produce taxusin (**6**) when TAX19 is added, it is only possible that T9αH-725A acts after TAX19, as shown on the upper route. The proposed intermediate 4O3A was quickly turned over by TAT and not detected in the TAX19 + T9αH-725A experiment.

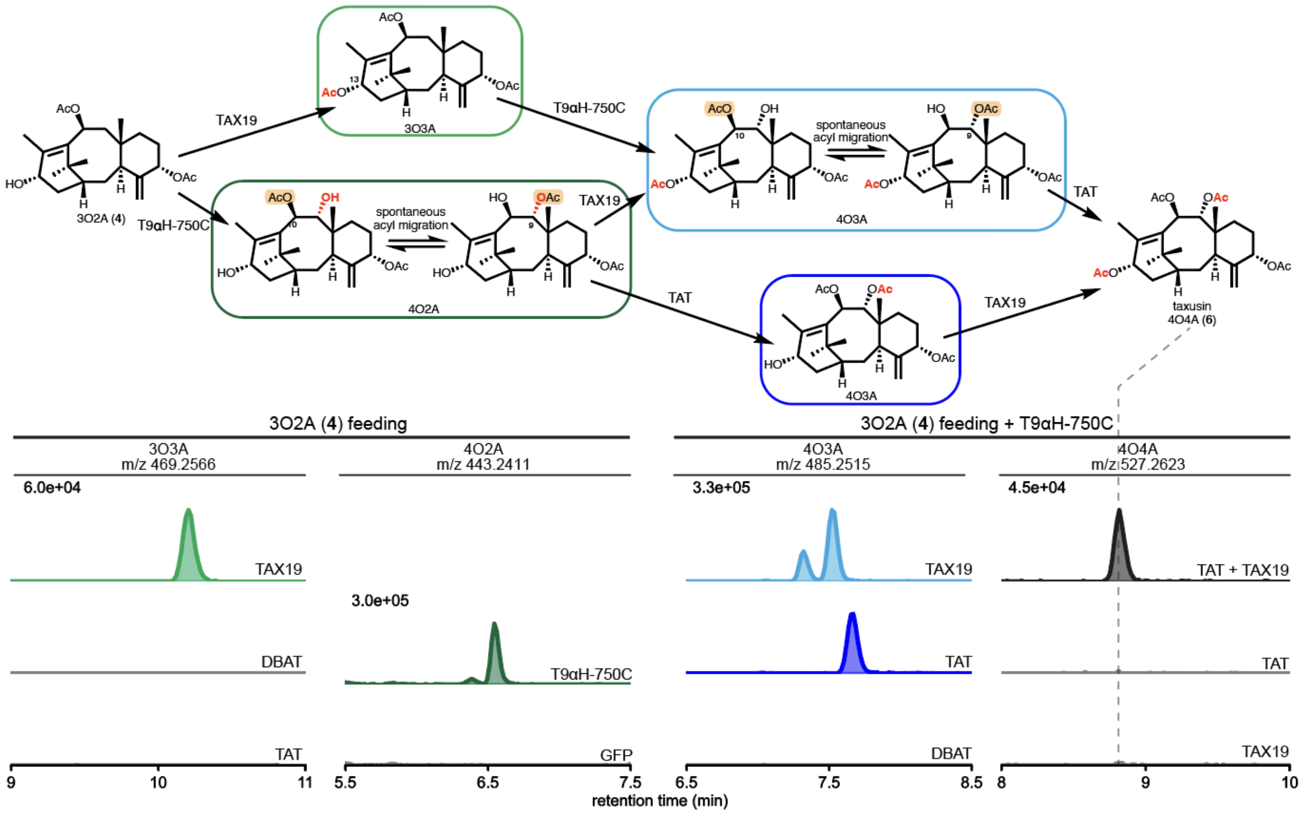


**Fig. S11. Metabolic network from 3O2A (4) to taxusin (6).** To investigate the origin of the acetyl group on 9α-hydroxy in taxusin (**6**) and study the biosynthetic routes, we fed purified 3O2A (**4**) to *N. benthamiana* leaves expressing different combinations of T9αH-750C, TAT, DBAT, and TAX19. We found that only TAX19 (light green), the previously reported C-13α-O-acetyltransferase, yielded an acetylated 3O3A product and neither DBAT nor TAT did. Both T9αH-750C (dark green) and T9αH-750C+TAX19 (light blue) condition resulted in the formation of two isomers of 4O2A and 4O3A, respectively, which we proposed is due to the spontaneous acyl migration between 9α- and 10β-hydroxyl groups. As taxane scaffolds are highly strained and congested, acyl migration of acetyl groups are poised to happen, for example, the migration from 10β- to 7β-hydroxy[30], and from 2α- to 5α-hydroxy[31] have been reported. T9αH-750C+TAT (dark blue) generated a single 4O3A acetylated product different from the two isomers generated by T9αH-750C+TAX19 (light blue). This suggests a 9α-*O*-acetylated product and the involvement of TAT on 9α-*O*-acetylation. Lastly, T9αH-750C+TAT+TAX19 condition successfully resulted in the production of taxusin (**6**). Overall, these data suggest a metabolic network model from 3O2A (**4**) to 4O4A involving TAT, TAX19, and T9αH-750C, and reveal the bi-functional role of TAT on both 5α- and 9α-*O*-acetylation.

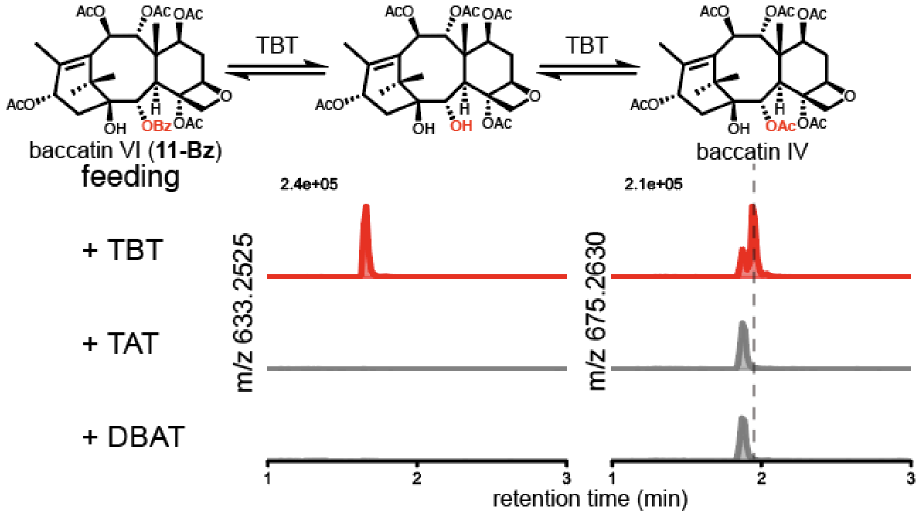


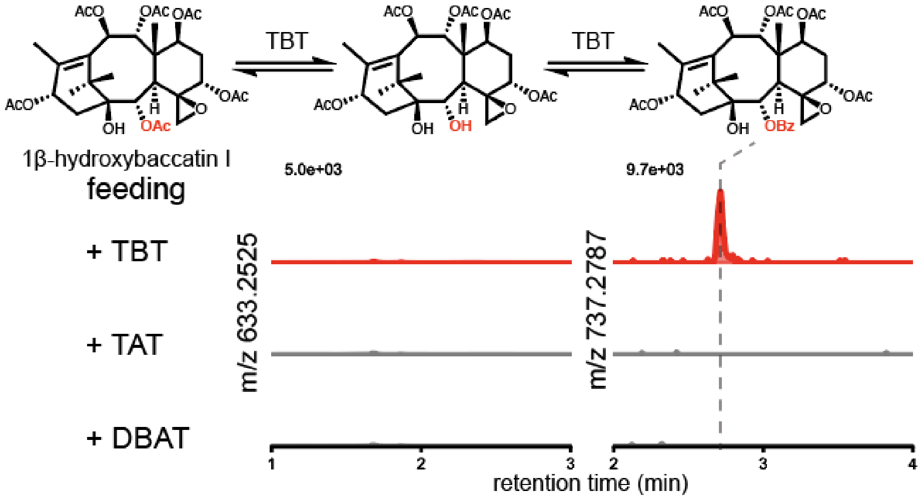


**Fig. S12**. **TBT mediates the interconversion between C-2α-benzoyl and acetyl group.** Baccatin VI (**11-Bz**) and 1β-hydroxybaccatin I were fed into *N. benthamiana* leaves expressing either TBT, TAT, or DBAT. Only TBT showed 2-*O*-debenzoylation activity and converted baccatin VI (**11-Bz**) into C-2α-debenzoylated and C-2α-debenzoylated acetylated products. TBT also converted 1β-hydroxybaccatin I into the corresponding C-2α-deacetylated benzoylated products; the deacetylated intermediate was not detected. The 2-*O*-debenzoylation activity of TBT has been characterized.[32]

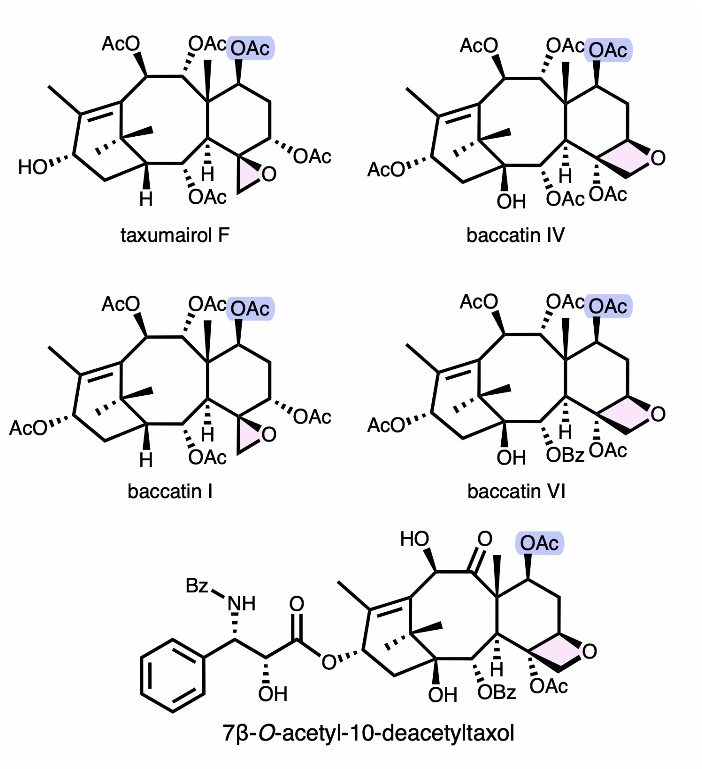


**Fig. S13. Structures of C-7β-acetylated taxane examples with oxetane or epoxide modification.** Many highly oxygenated taxanes with oxetane or epoxide group (highlighted in pink) derived from TOT activity are also C-7β-acetylated (highlighted in blue), including taxumairol F, baccatin I, baccatin IV, baccatin VI, and 7β-*O*-acetyl-10-deacetyltaxol.[33] This leads us to hypothesize that C-7β acetylation is a prerequisite for TOT function.

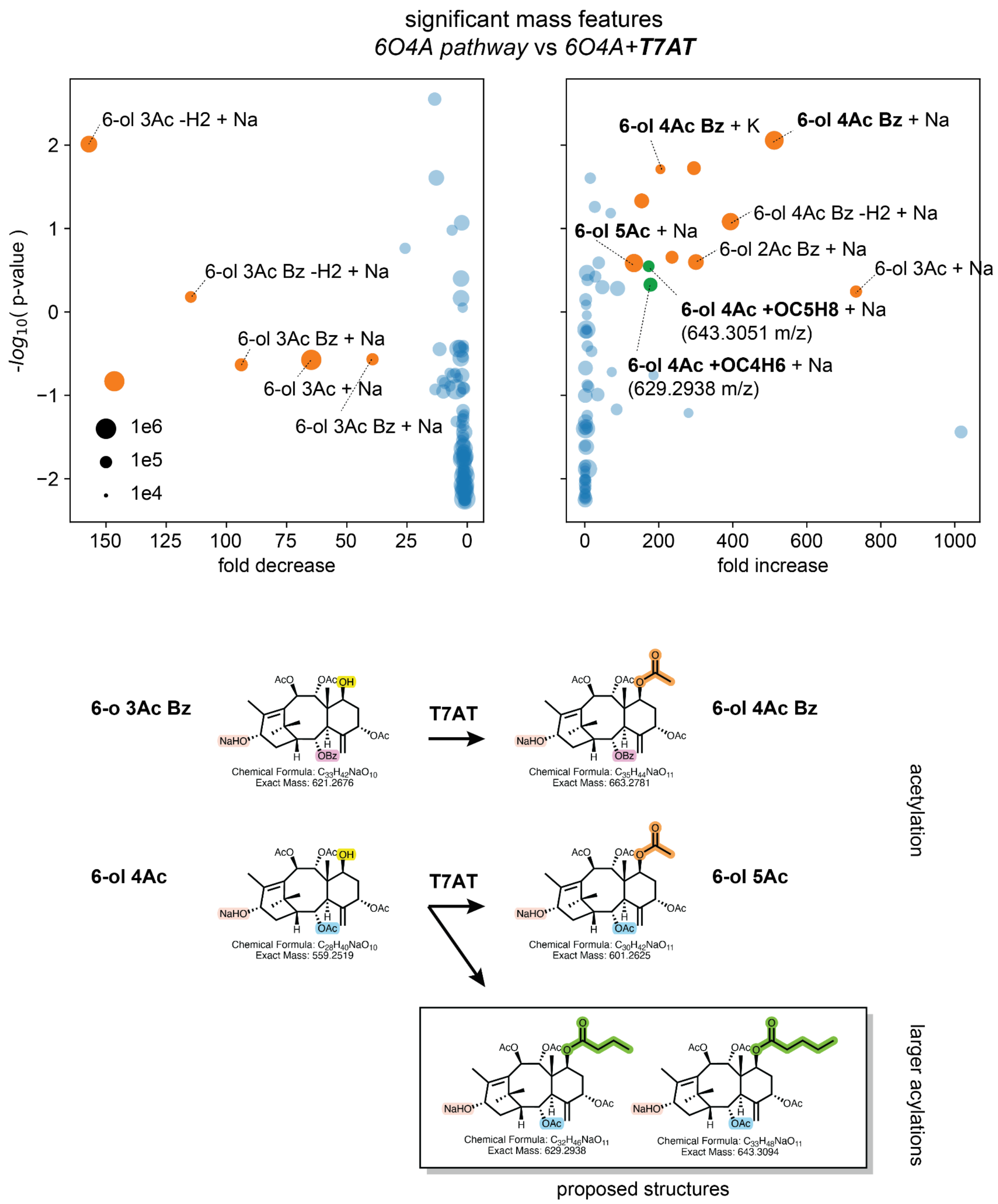


**Fig. S14. T7AT is an acyltransferase with multiple acylation activities.** Untransformed volcano plot comparing *N. benthamiana* leaves expressing the pathway to 6O4A with and without T7AT. Each mass feature is shown as a dot whose size indicates EIC integrated area. While the dominant product is the on-pathway 6-ol 4Ac Bz (m/z 663.2781 for M+Na adduct), additional minor products (green dots) correspond to the masses of C4 and C5 acylation products, suggesting that T7AT is capable of multiple acylation activities. Possible acylation structures are illustrated. Mass features corresponding to loss of two protons (-H2), which are likely C-13-ketone derivative,[34] also show significant changes. P-values calculated by t-test with a Bonferroni correction for multiple hypothesis testing.

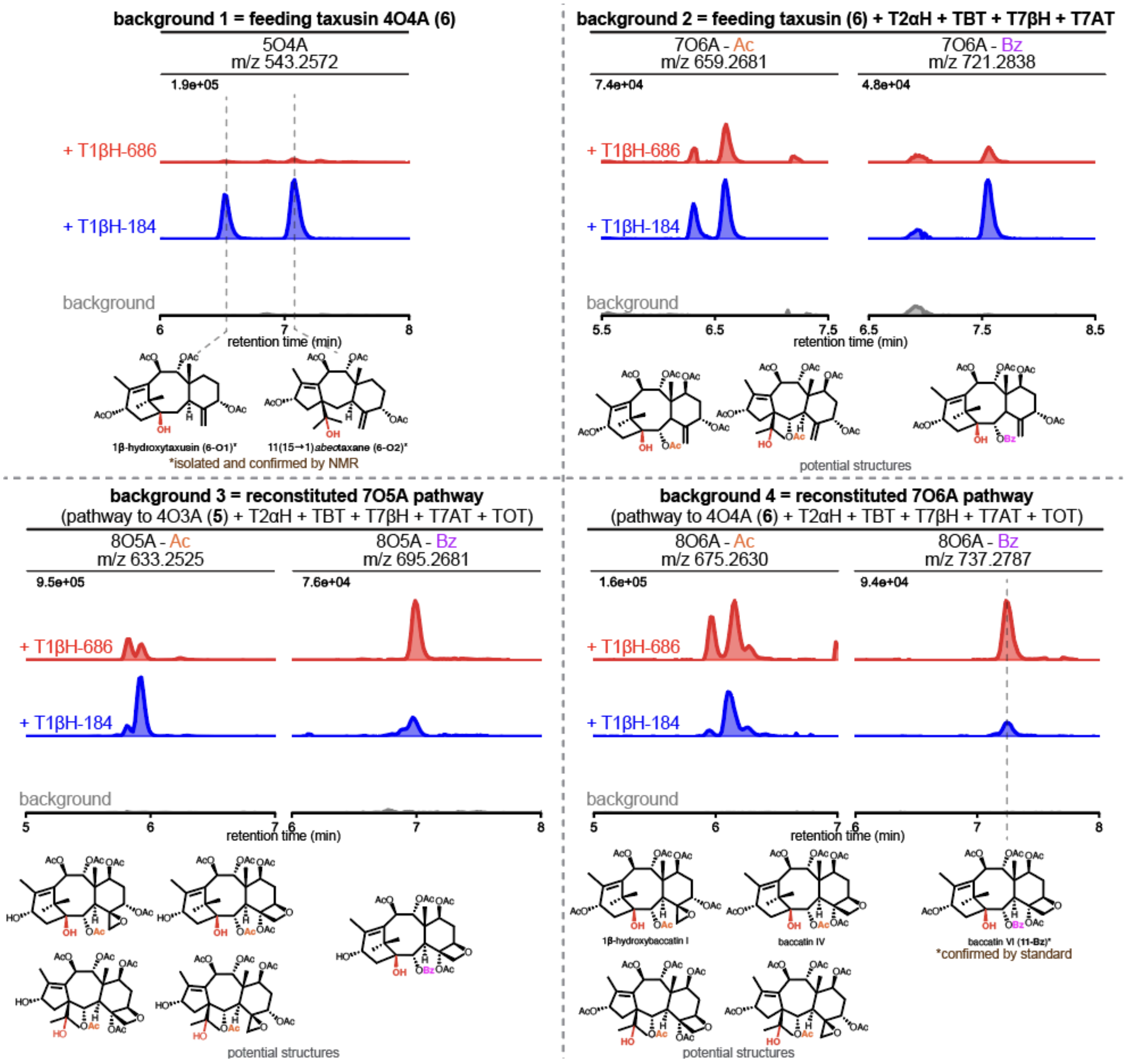


**Fig. S15. Product profiles of T1βH-184 and T1βH-686 with different upstream pathways.** Two 2-ODDs, T1βH-184 and T1bH-686, was expressed in *N. benthamiana* with four different background conditions: (1) feeding taxusin, (2) feeding taxusin and co-express T2αH, TBT, T7βH, T7AT, (3) co-express full biosynthetic pathway to 7O5A, i.e. HMGR, tGGPPS, TDS, FOTO, T5αH, TAT, T10βH, DBAT, T13αH, T2αH, TBT, T7βH, T7AT, TOT, and (4) co-express full biosynthetic pathway to 7O6A, i.e. HMGR, tGGPPS, TDS, FOTO, T5αH, TAT, T10βH, DBAT, T13αH, T2αH, TBT, T7βH, T7AT, TOT, TAX19. We proposed the potential structures of the products for each condition. Among all products, 1β-hydroxytaxusin (**6-O1**) and 15-hydroxy-11(15→1)*abeo*-taxusin (**6-O2**) are confirmed by NMR (**Table S4-6**) while baccatin VI (**11-Bz**) is confirmed by comparing to chemical standard (**Fig. 4c,e**). The observed product diversity (multiple peaks) likely arises from the dual-function of T1βH and TOT: T1βH performs both 1β-hydroxylation and rearrangement to 11(15→1)*abeo*taxane, and TOT generates both epoxide and oxetane products. In all conditions, C-2α-*O*-benzoylated products mostly show as single dominant peak, for example, baccatin VI (**11-Bz**). We proposed that the 1β-hydroxylation activity of T1βH is selective toward C-2α-*O*-benzoylated/oxetane intermediates.

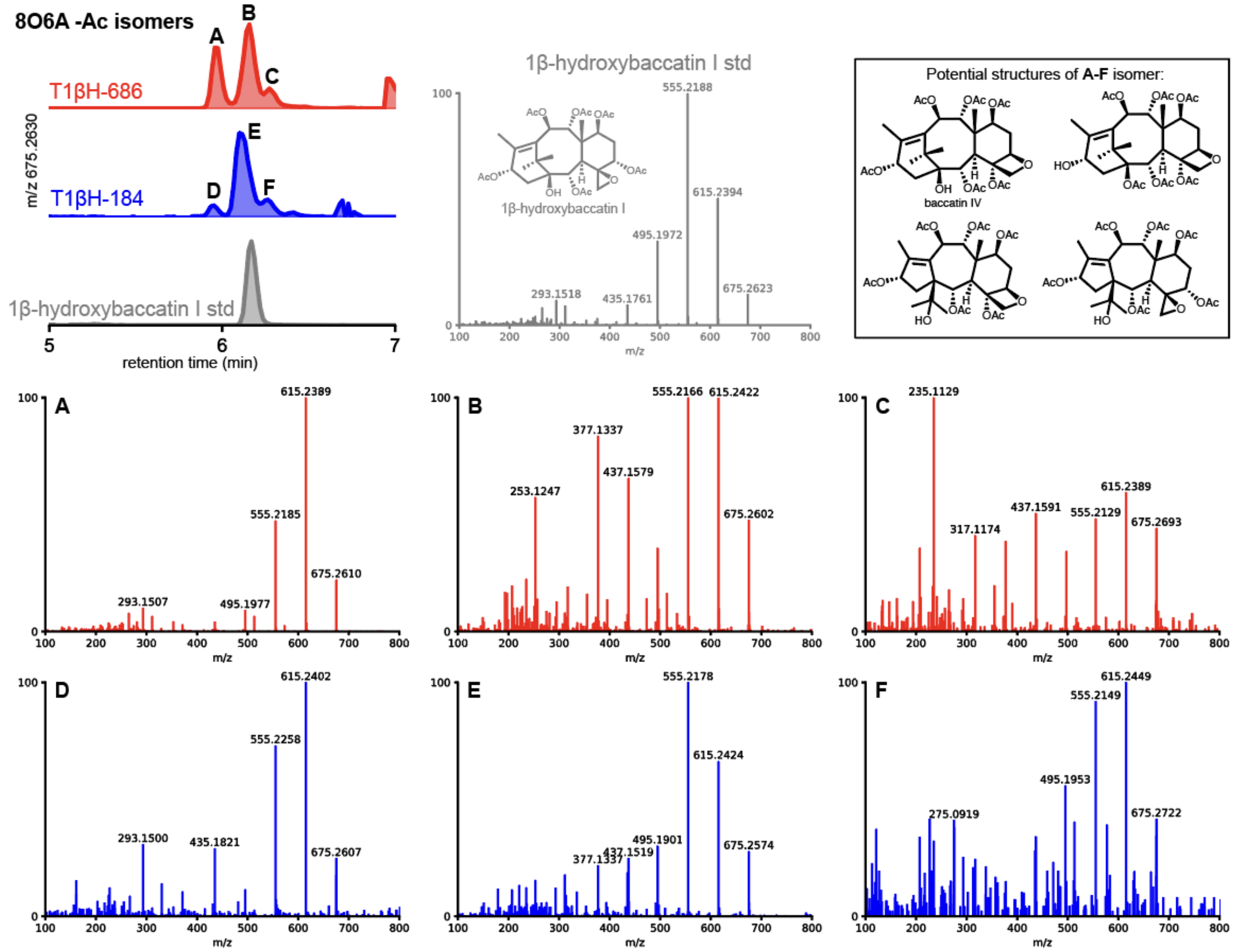


**Fig. S16**. **Production of hexaoxygenated hexacetylated (8O6A-Ac) isomers by the two T1βHs**. Both T1βH-184 and T1βH-686 produce 3 major 8O6A-Ac products (peak A-F). MSMS fragmentation patterns of all products and 1β-hydroxybaccatin I standard are shown. MSMS fragmentations were generated using $[M+Na]^+$ (m/z = 675.2630) as the precursor ion and fragmented with a collision energy of 30 eV. While peak B shares a similar retention time as 1β-hydroxybaccatin I, they show different MSMS patterns, thus are deemed as different compounds. The following gene sets were heterologously expressed in *N. benthamiana* via *Agrobacterium*-mediated transient expression: HMGR, tGGPPS, TDS, FOTO, T5αH, TAT, T10βH, DBAT, T13αH, TAX19, T2αH, TBT, T7βH, T7AT, TOT, T1βH-184 or T1βH-686.

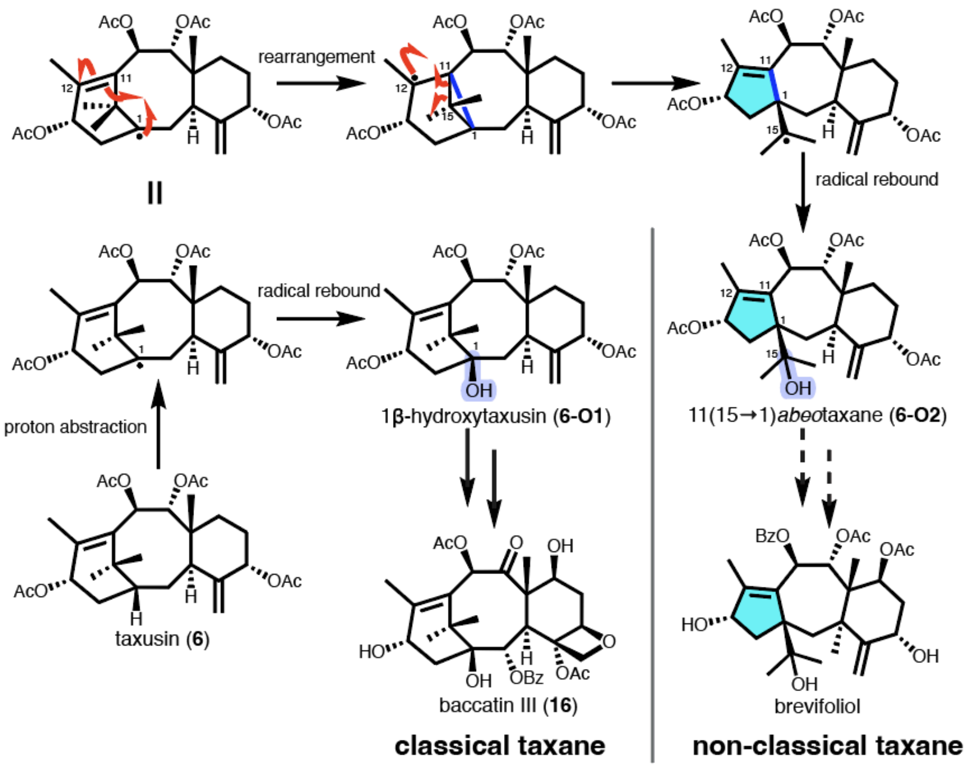


**Fig. S17**. **Proposed T1βH mechanism.** Formation of the two purified products 1β-hydroxytaxusin (**6-O1**) and 15-hydroxy-11(15→1)*abeo*-taxusin (**6-O2**) from taxusin (**6**) by T1βH-184 can be explained by the different fates of the C-1 radical after the first proton abstraction: direct hydroxyl radical rebound from the enzyme would yield 1β-hydroxytaxusin (**6-O1**) while radical rearrangement forming C1-11 bond followed by ring opening and hydroxyl radical rebound would give 15-hydroxy-11(15→1)*abeo*-taxusin (**6-O2**). We proposed that the rearrangement route leads to many non-classical *abeo*taxanes, like brevifoliol, while the 1β-hydroxylation leads to classical taxanes like baccatin III (**16**) and Taxol. Note that taxusin might not be the true biosynthetic intermediate for the highly modified taxanes.

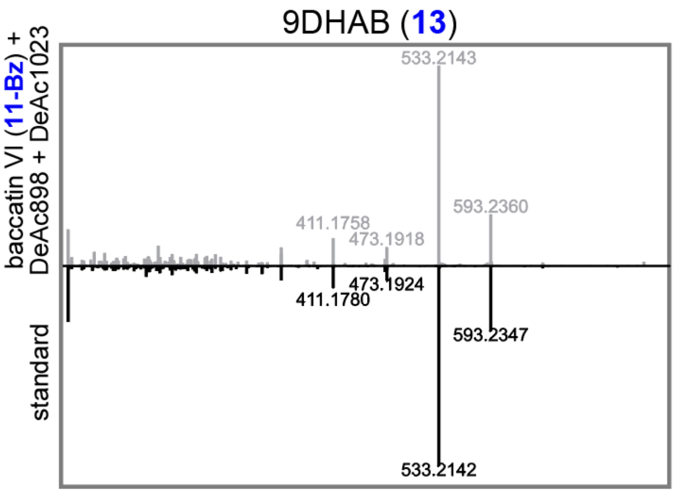


**Fig. S18. MSMS fragmentation patterns of 9-dihydro-13α-acetylbaccatin III (9DHAB, 13) when fed to *N. benthamiana* leaves expression DeAc898 and DeAc1023 compared to that of the standard**. MSMS fragmentations were generated using $[M+Na]^+$ (m/z = 653.2576) as the precursor ion and fragmented with a collision energy of 30 eV. Regions between m/z 100 to 800 are shown. DeAc898 and DeAc1023 were heterologously expressed in *N. benthamiana* via *Agrobacterium*-mediated transient expression and baccatin VI (**11-Bz**) 20 uM was fed to the leaves three days after infiltration.

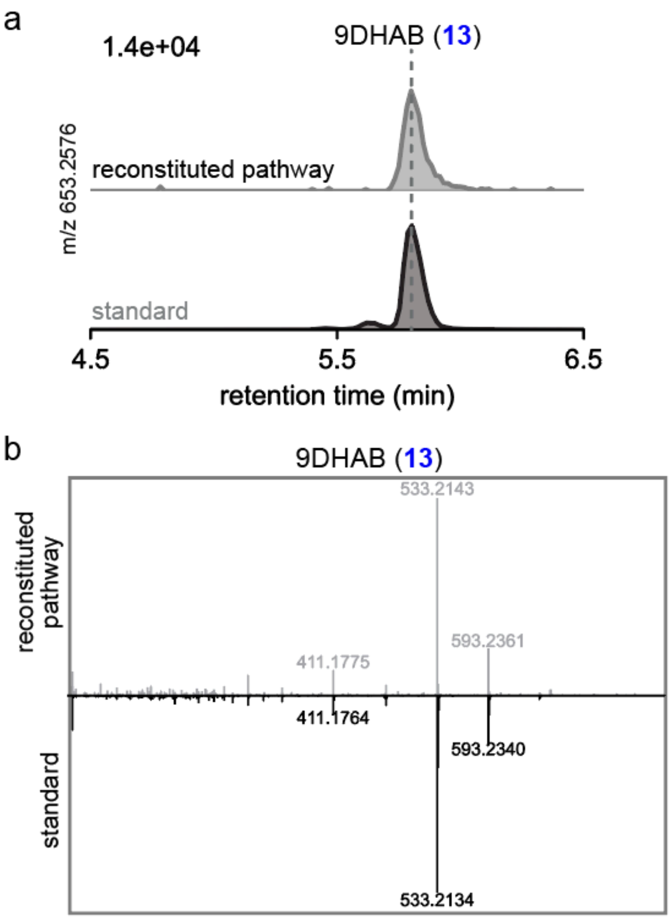


**Fig. S19. Reconstituted biosynthesis of 9-dihydro-13α-acetylbaccatin III (9DHAB, 13) in *N. benthamiana*.** (**a**) EICs of leaves expressing the 9DHAB biosynthetic pathway (below) compared to the 9DHAB standard. (**b**) MSMS fragmentation patterns of heterologously produced 9DHAB (**13**) compared to that of the standard. MSMS fragmentations were generated using $[M+Na]^+$ (m/z = 653.2576) as the precursor ion and fragmented with a collision energy of 30 eV. Regions between m/z 100 to 800 are shown. The following gene set was heterologously expressed in *N. benthamiana* via *Agrobacterium*-mediated transient expression: HMGR, tGGPPS, TDS, FoTO, T5αH, TAT, T10βH, DBAT, T13αH, T9αH-CYP750C, TAX19, T2αH, TBT, T7βH, T7AT, TOT, T1βH-686, DeAc898, DeAc1023, and T9ox.

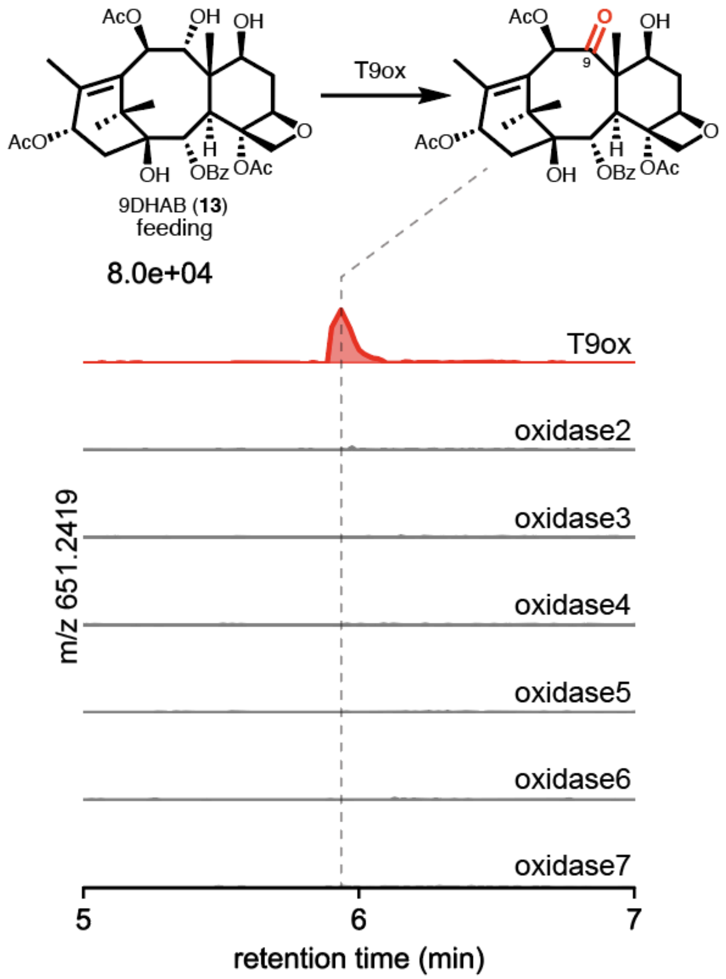


**Fig. S20**. **Screening of oxidases identified the taxane C-9 oxidase (T9ox)**. *N. benthamiana* leaves expressing individual candidate oxidase via *Agrobacterium*-mediated infiltration and fed with 9DHAB (**13**) at 3 day post infiltration (dpi). Over 30 P450s and 2-ODDs were screened in batches and the only active batch with 8 candidates was deconvoluted as shown here. EIC of C-9 ketone product (m/z 651.2419) was shown.

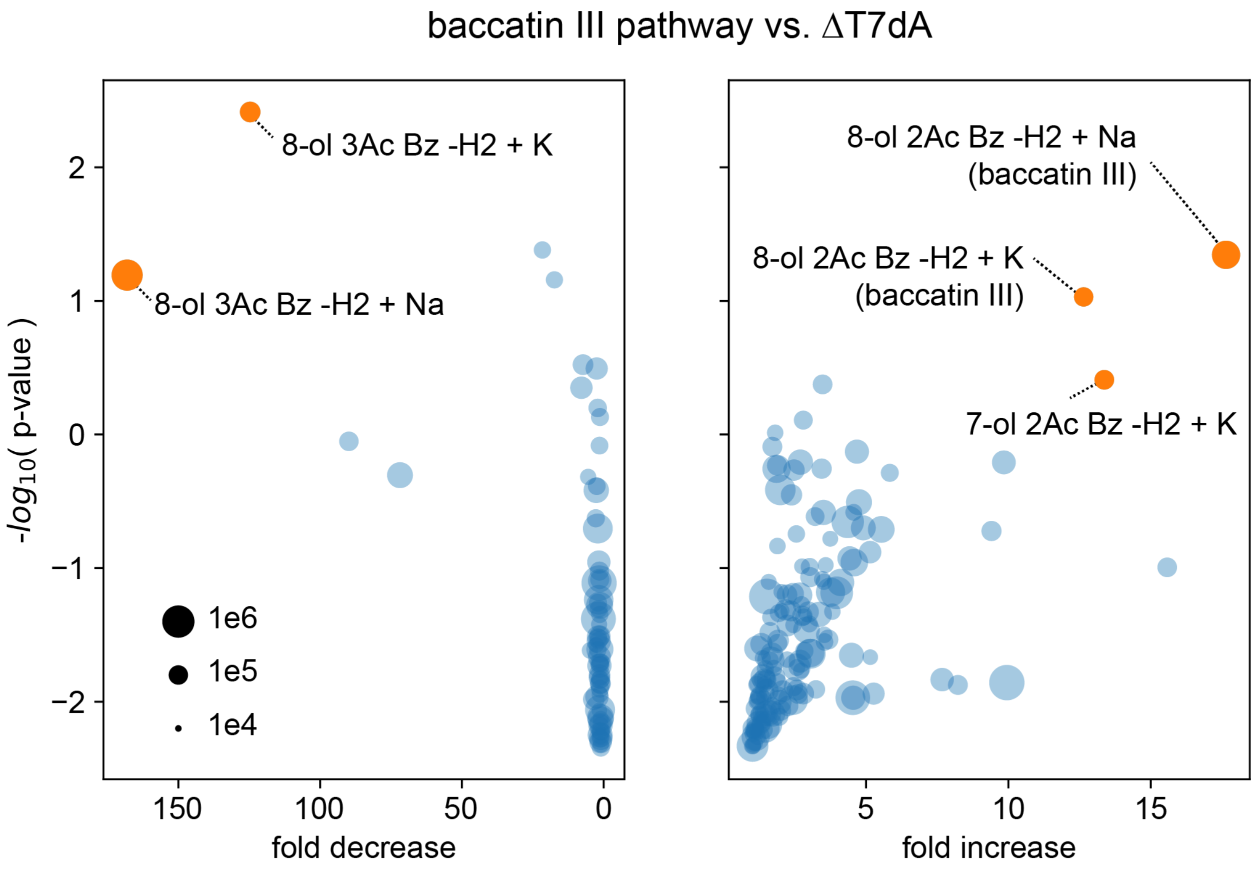


**Figure S21**. **Untargeted analysis shows baccatin III (16) as a major product in the final step of pathway.** Linear volcano plot comparing *N. benthamiana* leaves expressing the full baccatin III (**16**) pathway and full pathway without the penultimate enzyme T7dA. Metabolomic features with putative taxane masses are shown are dots whose sizes indicate EIC integrated area. P-values calculated by t-test with a Bonferroni correction for multiple hypothesis testing. The full baccatin III (**16**) pathway include HMGR, tGGPPS, TDS, FoTO, T5αH, TAT, T10βH, DBAT, T13αH, T9αH-CYP750C, T2αH, TBT, T7βH, T7AT, TOT, T1βH-686, DeAc898 (T9dA), DeAc1023 (T7dA), and T9ox.

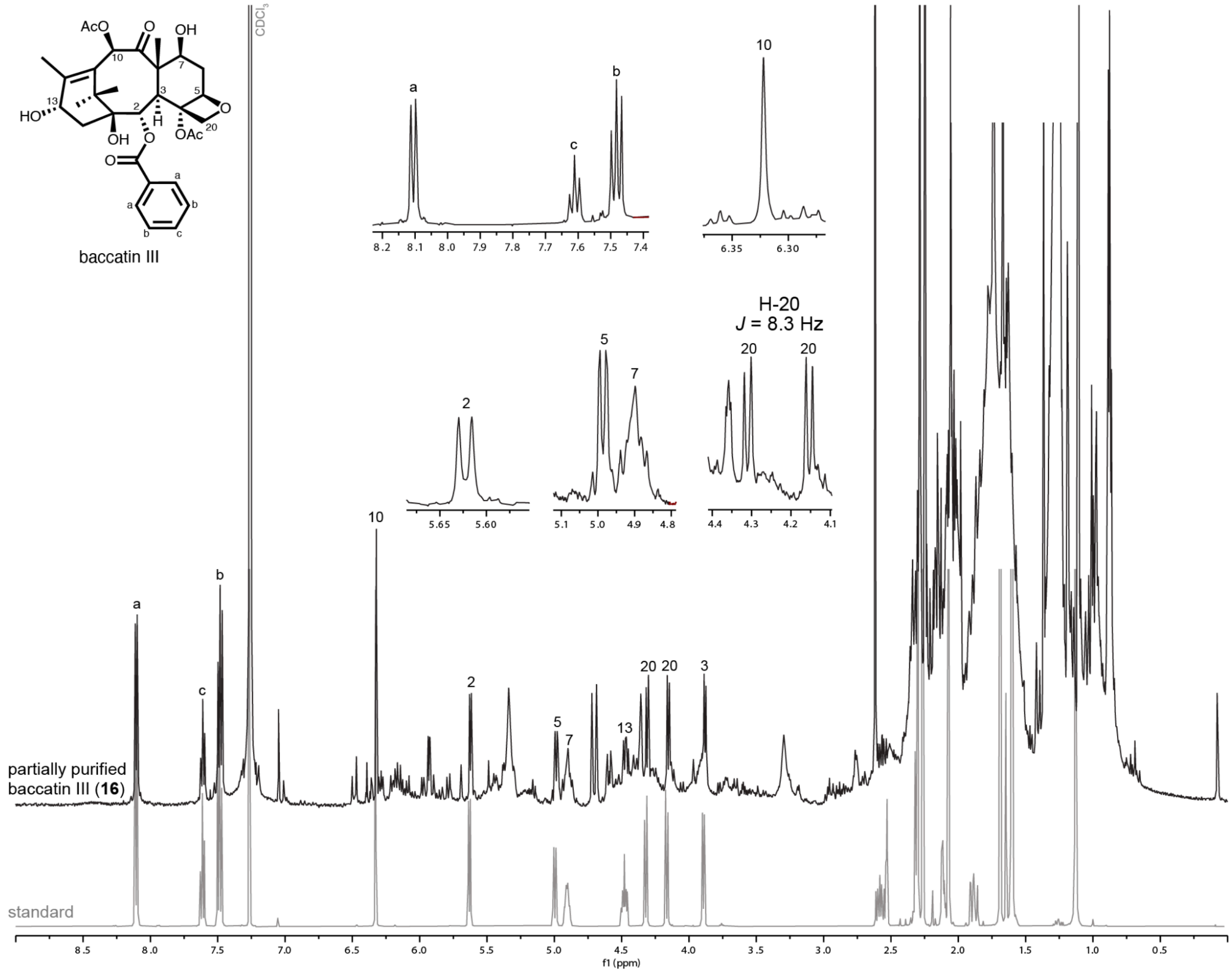


**Figure S22**. **$^1$H-NMR spectra of partially purified baccatin III (16) and baccatin III (16) standard in $CDCl_3$.** The spectra of our partially purified baccatin III (**16**) align with the standard, exhibiting all characteristic peaks (labeled) as well as the H-20 coupling constant ($J$ = 8.3 Hz) of the oxetane. The following gene set was heterologously expressed in 53 *N. benthamiana* plants via *Agrobacterium*-mediated transient expression: HMGR, tGGPPS, TDS, FoTO, TAT, T10βH, DBAT, T13αH, T9αH-CYP750C, T2αH, TBT, T7βH, T7AT, TOT, T1βH-686, DeAc898, DeAc1023, and T9ox. Note that T5αH was excluded from this gene set, as we show that T5αH is not required for the complete biosynthesis of baccatin III (**16**) (**Fig. 6c**).

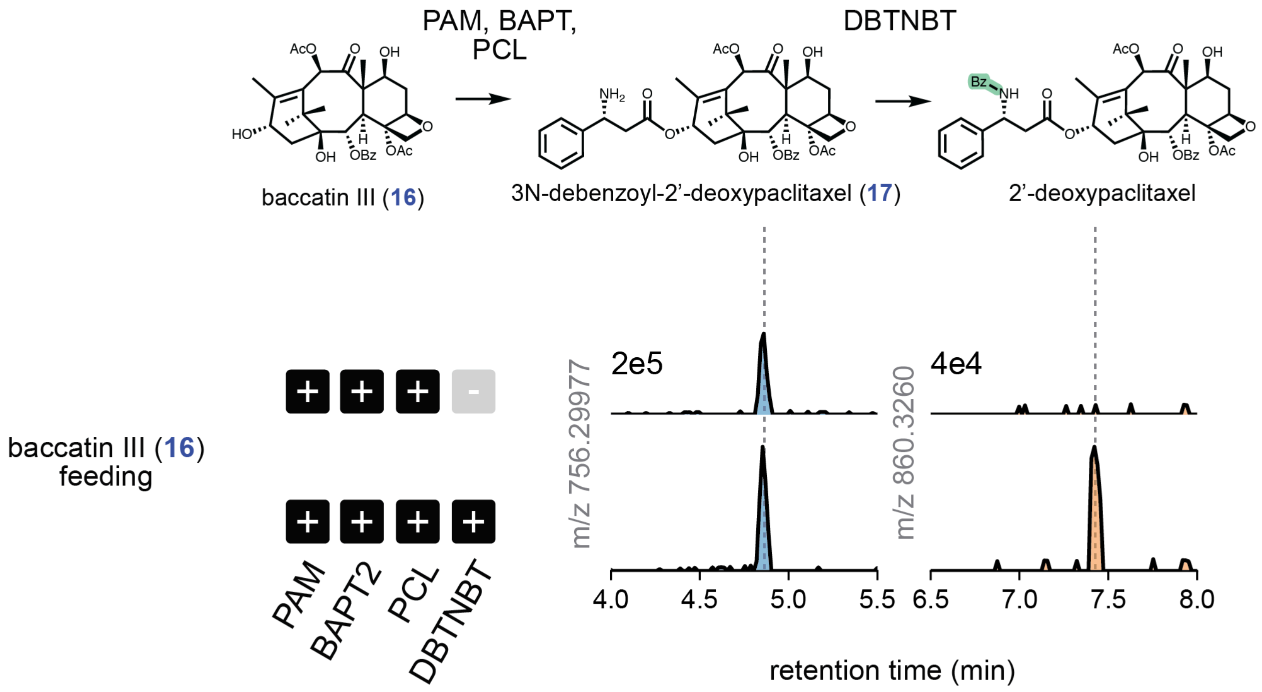


**Figure S23**. **Production of putative 2'-deoxypaclitaxel at low level**. EICs of *N. benthamiana* leaves expressing side chain enzymes (PAM, BAPT, PCL) with or without DBTNBT via *Agrobacterium*-mediated infiltration and fed with 25 µM baccatin III (**16**) at 3 day post infiltration (dpi). DBTNBT yields a new product whose mass matches the expected product, 2'-deoxypaclitaxel The low conversion from 3'-*N*-debenzoyl-2'-deoxypaclitaxel (**17**) to 2'-deoxypaclitaxel suggests that 2'-α-hydroxylation is a prerequisite for 3'-*N*-benzoylation, which is consistent with previous hypothesis[35].

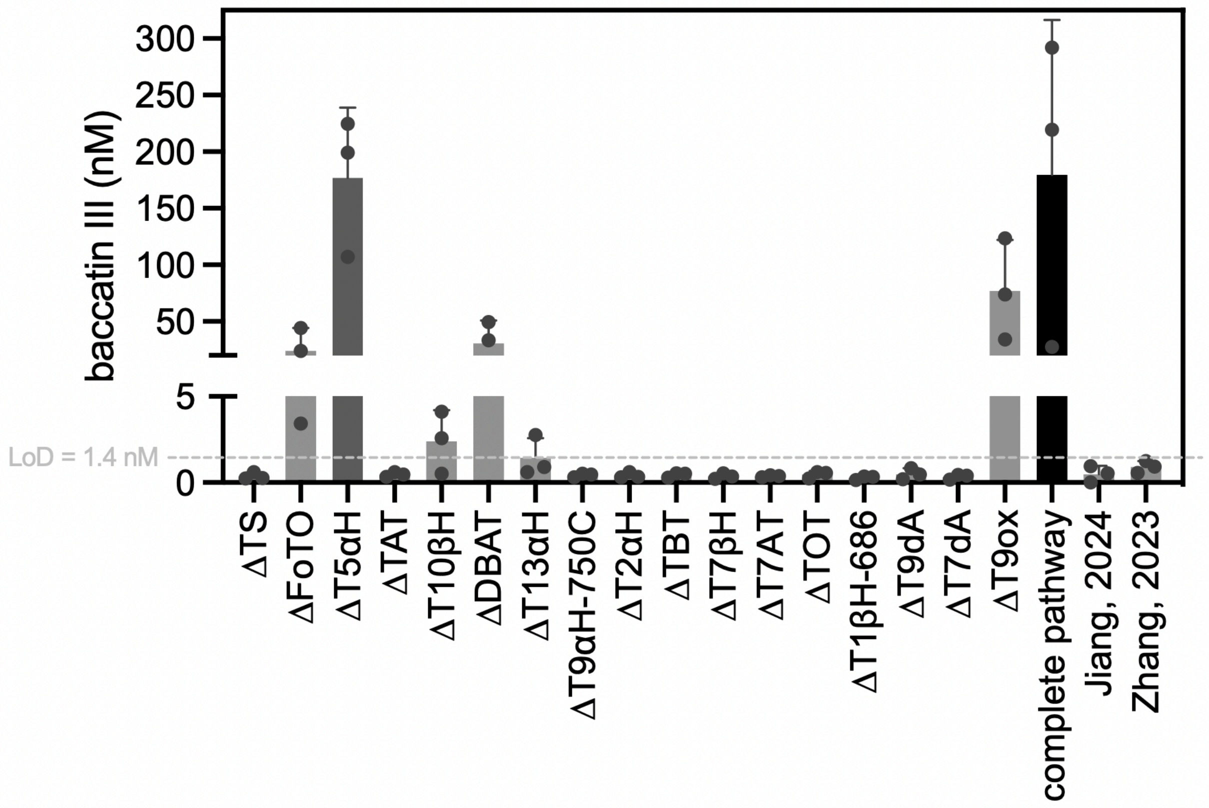


**Fig. S24. Bar graph of baccatin III (16) concentrations of the samples in Fig. 6d.** Baccatin III (**16**) concentration of *N. benthamiana* leaves expressing the indicated strain combinations via *Agrobacterium*-mediated infiltration. Leaf discs (6*1 cm diameter per sample; average dried weight = 10 mg) were harvested and extracted with 700 uL extraction buffer (75% ACN/water) and subjected to Agilent 6470 QQQ LCMS analysis. Limit of detection (LoD) is calculated using equation LoD = 3.3*SD/slope of calibration, where SD is the standard deviation (SD) of low concentration samples (10 pM baccatin III standard, n=10). Data are shown as the mean ± standard deviation, n = 3. See **Methods** and **Source Data** for detailed experimental setup and raw values.

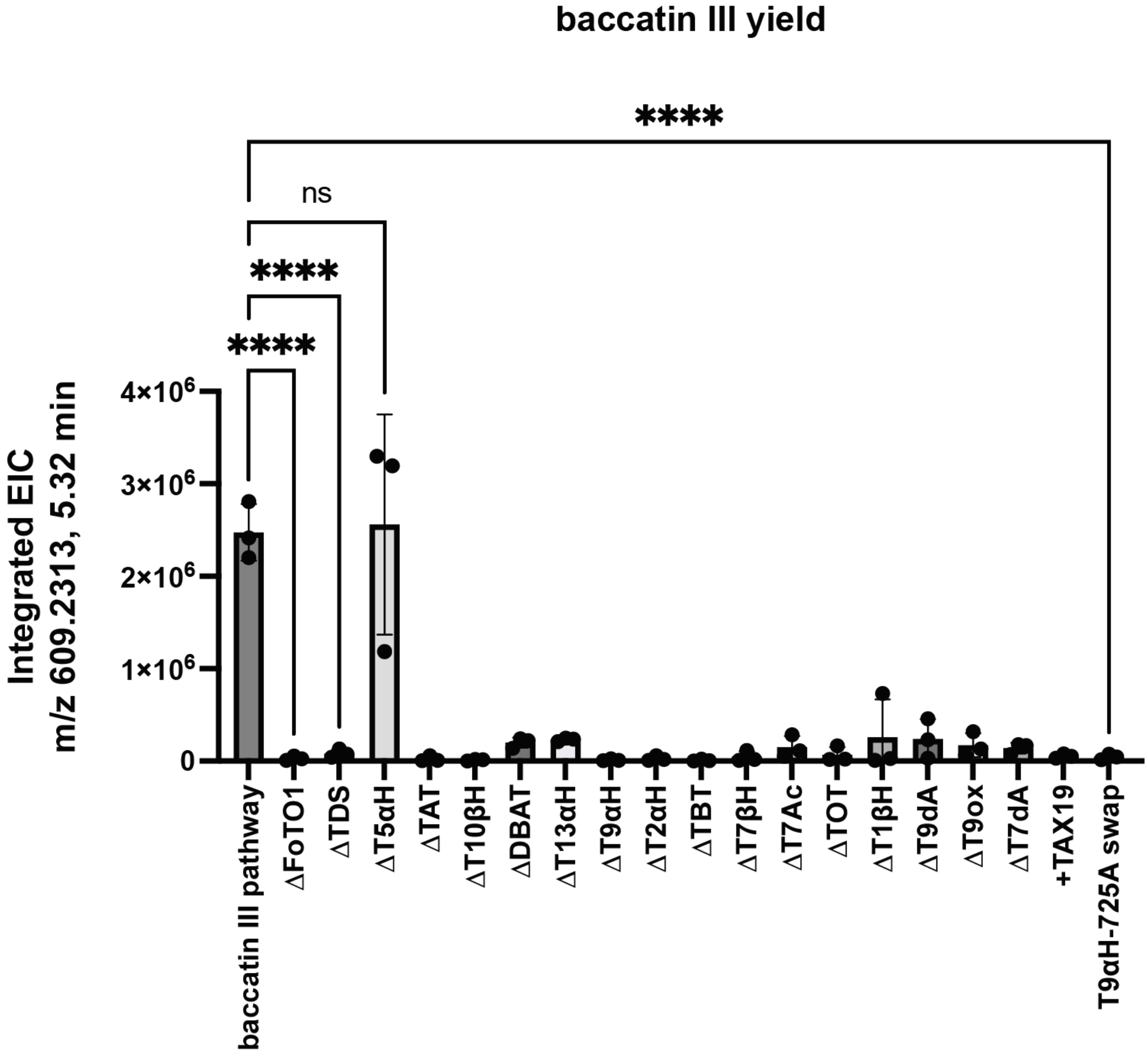


**Fig. S25. Independent replicate data for full-pathway gene dropout experiment.** EIC peak area of baccatin III (**16**) from *N. benthamiana* leaves expressing the full 17-gene baccatin III pathway, as well as single gene dropouts, addition of Tax19, or exchanging our T9αH-750C for the alternative T9αH-725A. Samples were subjected to Agilent 6520 Q-TOF LCMS analysis. Significance indicates results of an ordinary one-way ANOVA comparison to the full pathway (**** indicates $p < 0.0001$).

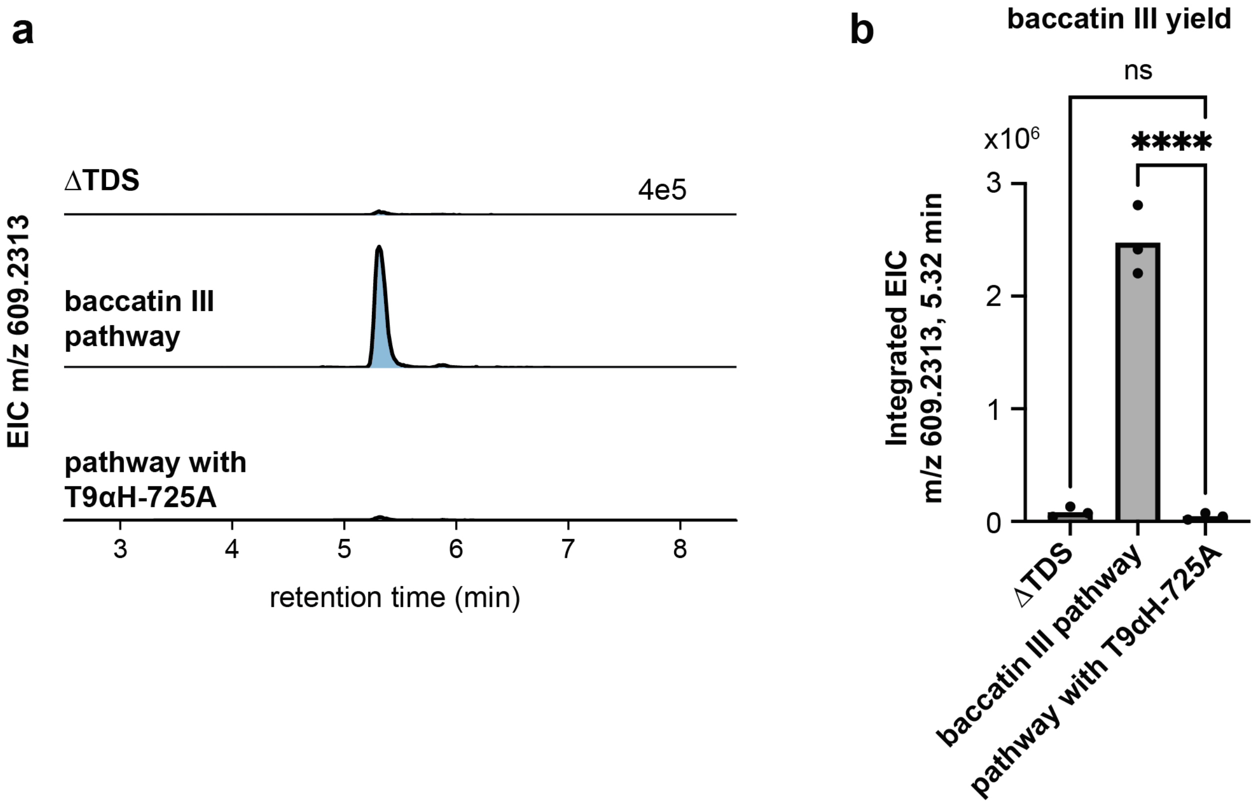


**Fig. S26. Baccatin III (16) production with T9αH-750C exchanged for T9αH-725A.** EIC of *N. benthamiana* leaves expressing our full baccatin III pathway gene set, compared to full gene set without TDS (ΔTDS) or an exchange of our T9αH-750C for the recently reported T9αH-725A.[7–9] (**a**) EIC traces baccatin III (m/z 609.2313) and (**b**) quantification of integrated EICs for this peak across three replicates. T9αH-725A pathway yields negligible baccatin III, statistically indistinguishable to a ΔTDS negative control. This would be expected from our finding that T9αH-725A appears to require a 13α-*O*-acetylation (**Fig. 3b**) that is absent from our baccatin pathway and from Taxol. The full baccatin III (**16**) pathway genes include HMGR, tGGPPS, TDS, FoTO, T5αH, TAT, T10βH, DBAT, T13αH, T9αH-CYP750C, T2αH, TBT, T7βH, T7AT, TOT, T1βH-686, DeAc898 (T9dA), DeAc1023 (T7dA), and T9ox. Significance indicates results of an ordinary one-way ANOVA comparison to the full pathway (**** indicates $p < 0.0001$).

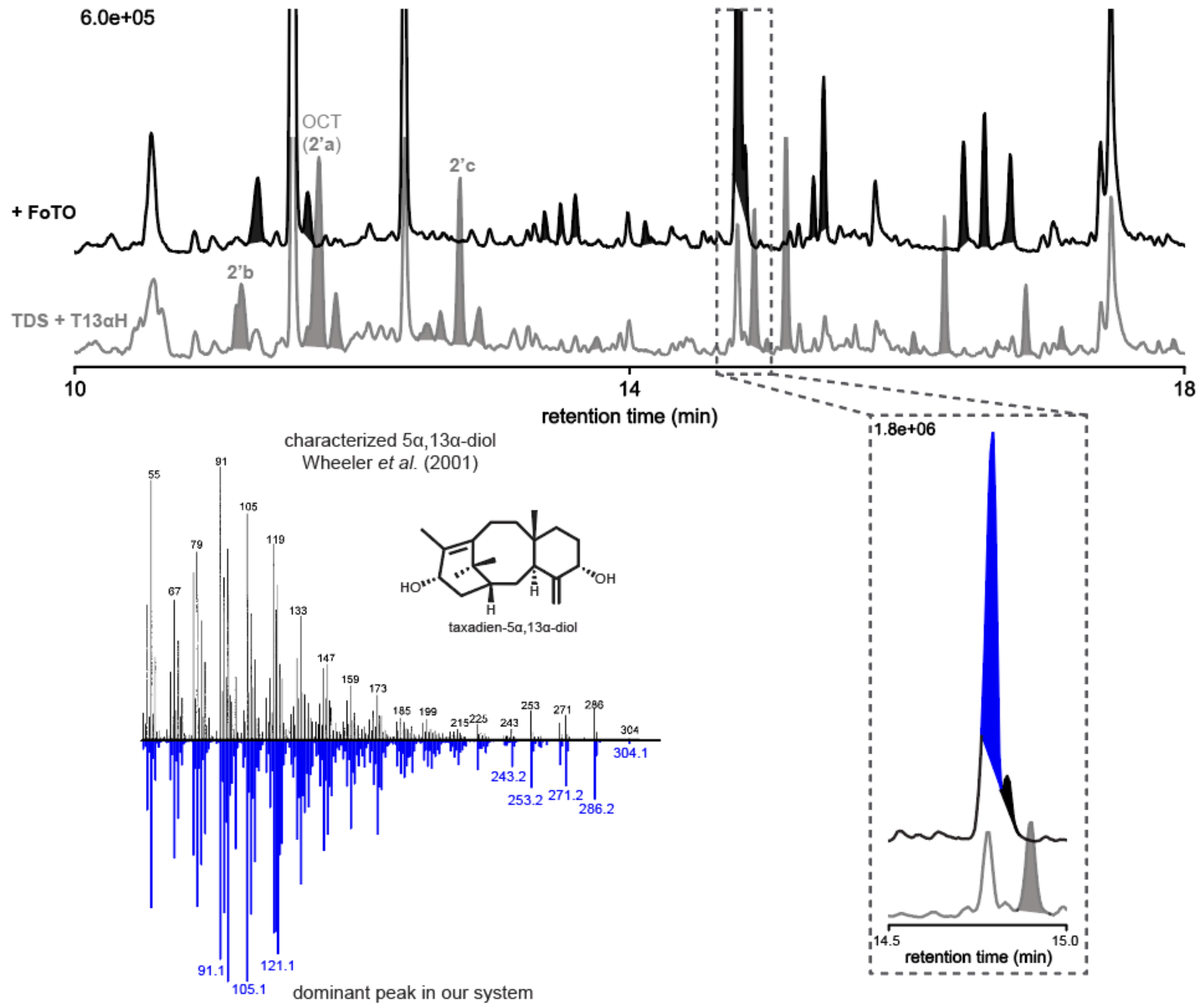


**Fig. S27. FoTO1 affects T13αH oxidation on taxadiene (1).** GCMS TIC of *N. benthamiana* leaves expressing TDS + T13αH and TDS + T13αH + FoTO1 (both with HMGR and tGGPPS) and the MS fragmentation patterns of the dominant product in the +FoTO1 condition are shown. Without FoTO, T13αH oxidizes taxadiene (**1**) to multiple products, including OCT (**2'a**), iso-OCT (**2'b**), rearranged products (**2'c**) that are also made by T5αH, and others. Similar to the effect of FoTO1 on T5αH, FoTO1 significantly changes the product profile of T13αH. The dominant peak (colored in blue) was proposed to be taxadien-5α,13α-diol (m/z 304) based on its MS comparison to previously published MS spectra[36] and its reactivity toward TAT and TAX19 (**Fig. S28**). Regions between m/z 50 to 320 are shown for the MS spectrum.

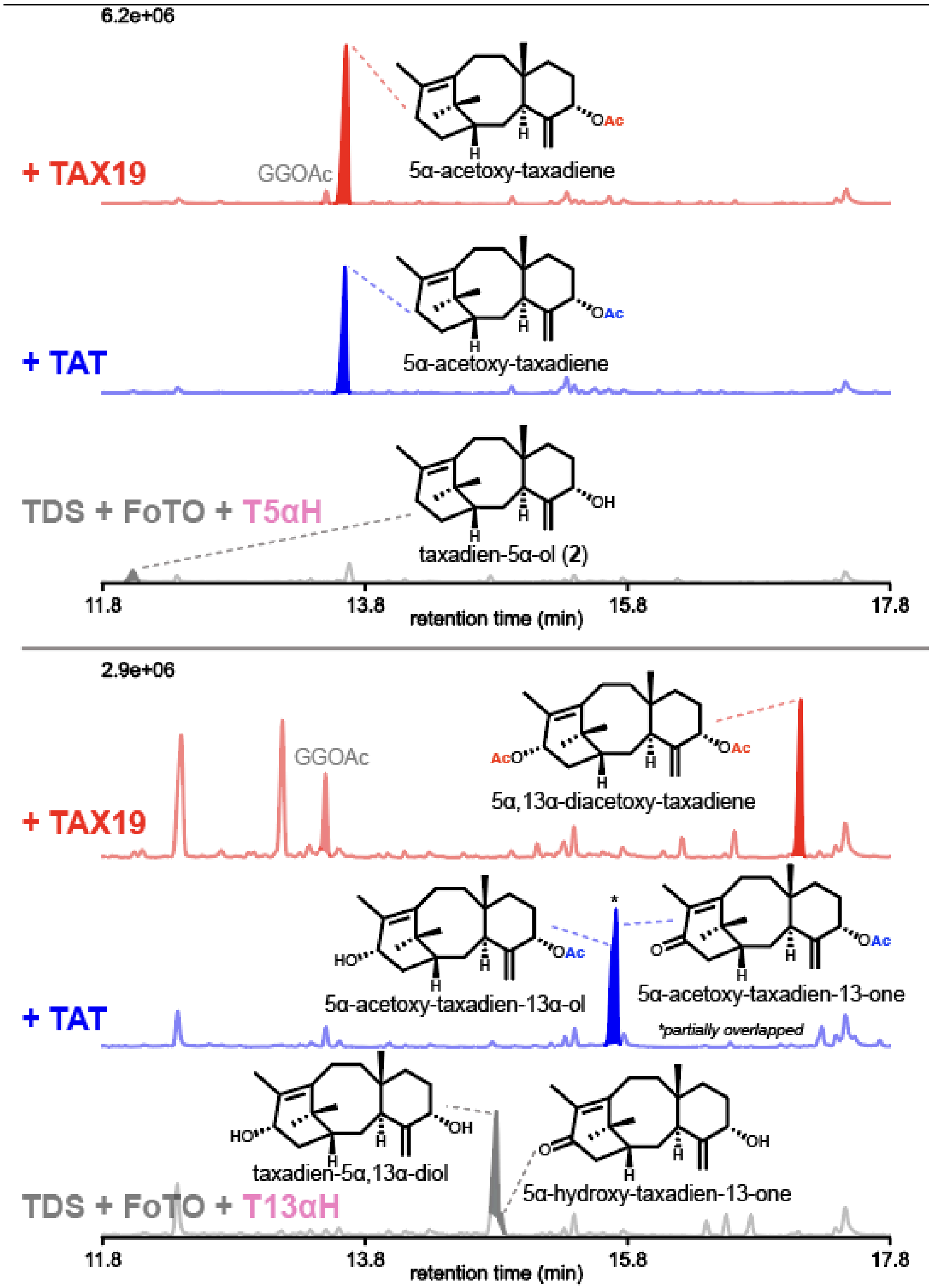


**Fig. S28. Products of T5αH and T13αH can be acetylated by TAT or TAX19.** GCMS TIC of *N. benthamiana* leaves expressing TDS + FoTO1 + T5αH + TAT or TAX19 (top) and TDS + FoTO1 + T13αH + TAT or TAX19 (bottom), all with HMGR + tGGPPS. Both TAX19 and TAT can acetylate taxadien-5α-ol (**2**) to 5α-acetoxy-taxadiene. However, TAX19 and TAT acetylate T13αH products (**Fig. S27**) differently: TAT results in two major products, presumably 5α-acetoxy-taxadien-13α-ol and 5α-acetoxy-taxadien-13-one, while TAX19 yields one major diacetoxy-taxadiene product (among other uncharacterized products). These data are consistent with previous characterization of TAX19's regioselective 5α- and 13α-*O*-acetylation activity.[37] All structures, except taxadien-5α-ol (**2**) are proposed based on the characterized enzyme functions and MS fragmentation patterns.

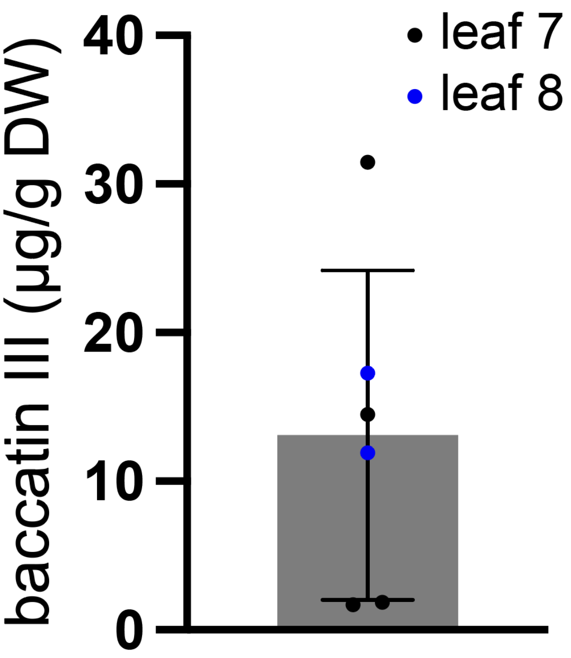


**Fig. S29. Variations in baccatin III (16) production.** Bar graph showing baccatin III (**16**) yields from *N. benthamiana* leaves expressing the full 17-gene pathway from different plants. Each sample (dot) was harvested from the 7th or 8th leaf counted from the bottom from separate plants under the same experimental conditions. The yields range from 1.7~31.5 µg/g dry weight (DW), comparable to the ranges reported from the twig and leaf samples from *T. chinensis, T. cupsidata,* and *T. media.*[38] Data are shown as the mean ± standard deviation, n = 4. The full baccatin III (**16**) pathway genes include HMGR, tGGPPS, TDS, FoTO, T5αH, TAT, T10βH, DBAT, T13αH, T9αH-CYP750C, T2αH, TBT, T7βH, T7AT, TOT, T1βH-686, DeAc898 (T9dA), DeAc1023 (T7dA), and T9ox.

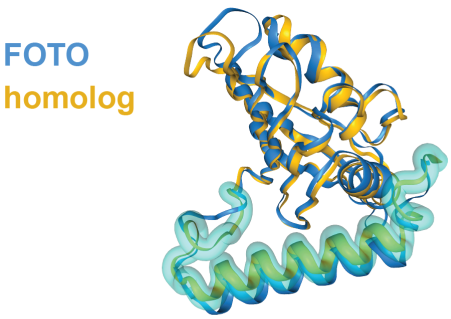


Q5JJV6 (*Oryza sativa Japonica*)
TM-Score: 0.87009

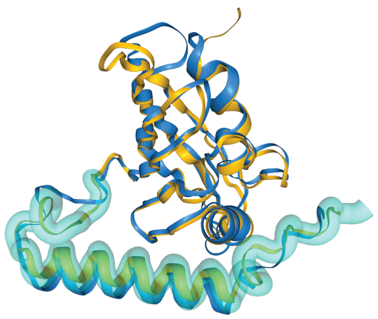

K7V1E4 (*Zea mays*)
TM-Score: 0.86595

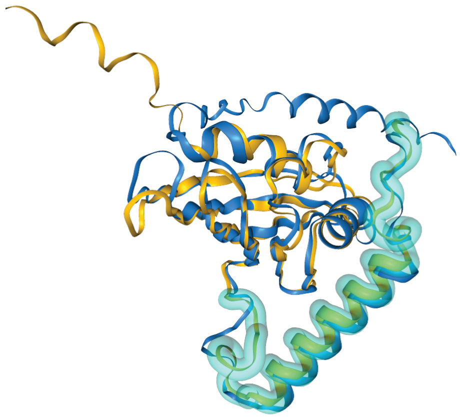

Q9FMC7 (*Arabidopsis thaliana*)
TM-Score 0.79643

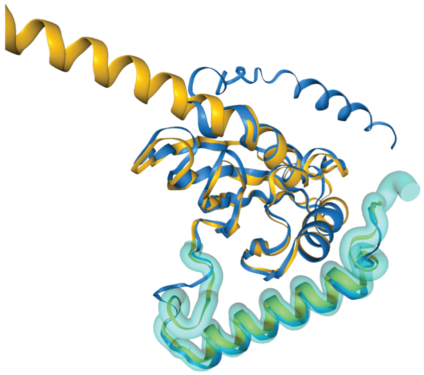

I1LCK9 (*Glycine max*)
TM-Score: 0.79063

**Fig. S30. FoTO1 structural homologs identified in other land plants**. Using FoldSeek v4[39] we searched all pre-folded protein databases in FoldSeek for full-length structural homologs of FoTO1. The top five hits were proteins from the genomes of model plants *Oryza sativa* Japonica (rice), *Zea mays* (corn), *Arabidopsis thaliana* and *Glycine max* (soy). After the NTF2 domain, each of these homologs contain the alpha-helical C-terminus (highlighted in transparent turquoise) that we found to be crucial for FOTO1's phenotype *in vivo* and for binding to TDS and T5αH. The identification these structural homologs indicates that FoTO1 is not just restricted to gymnosperms, but has structural analogs across both angiosperms.

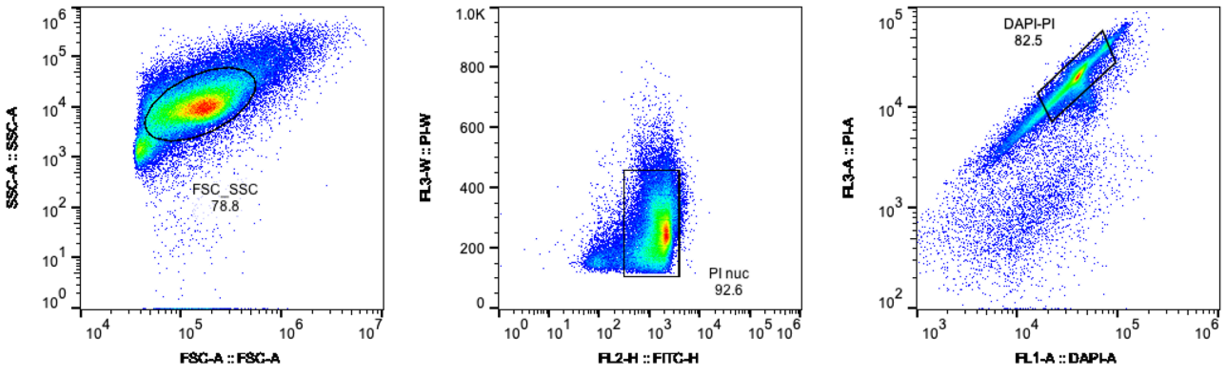


**Fig. S31. Gating strategy for nuclei sorting**. FACS plots generated with FlowJo v10 indicating the three step gating strategy to sort intact Taxus nuclei. Following the tissue disruption and lysate filtering outlined in **Methods**, nuclei are stained with 5ng/uL 4,6-diamidino-2-phenylindole (DAPI) and 5ng/uL propidium iodide (PI). Nuclei are gated on three subsequent gates: (i) size selection using forward scatter (FSC) vs side scatter (SSC), (ii) singlet selection using PI fluorescence height and width, and (iii) co-staining with DAPI and PI to identify clean nuclei.

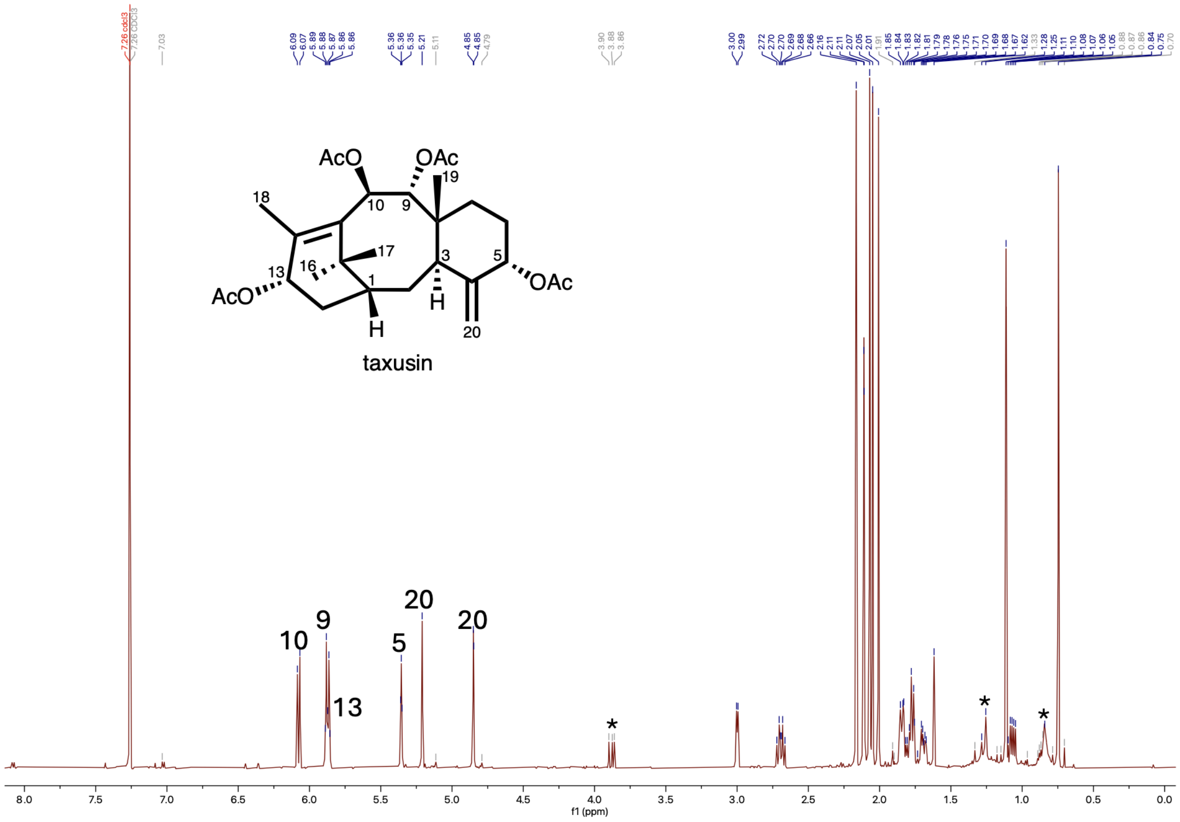


**Fig. S32. $^{1}$H-NMR spectrum of taxusin (6) in $CDCl_3$.** Asterisk indicates impurities.

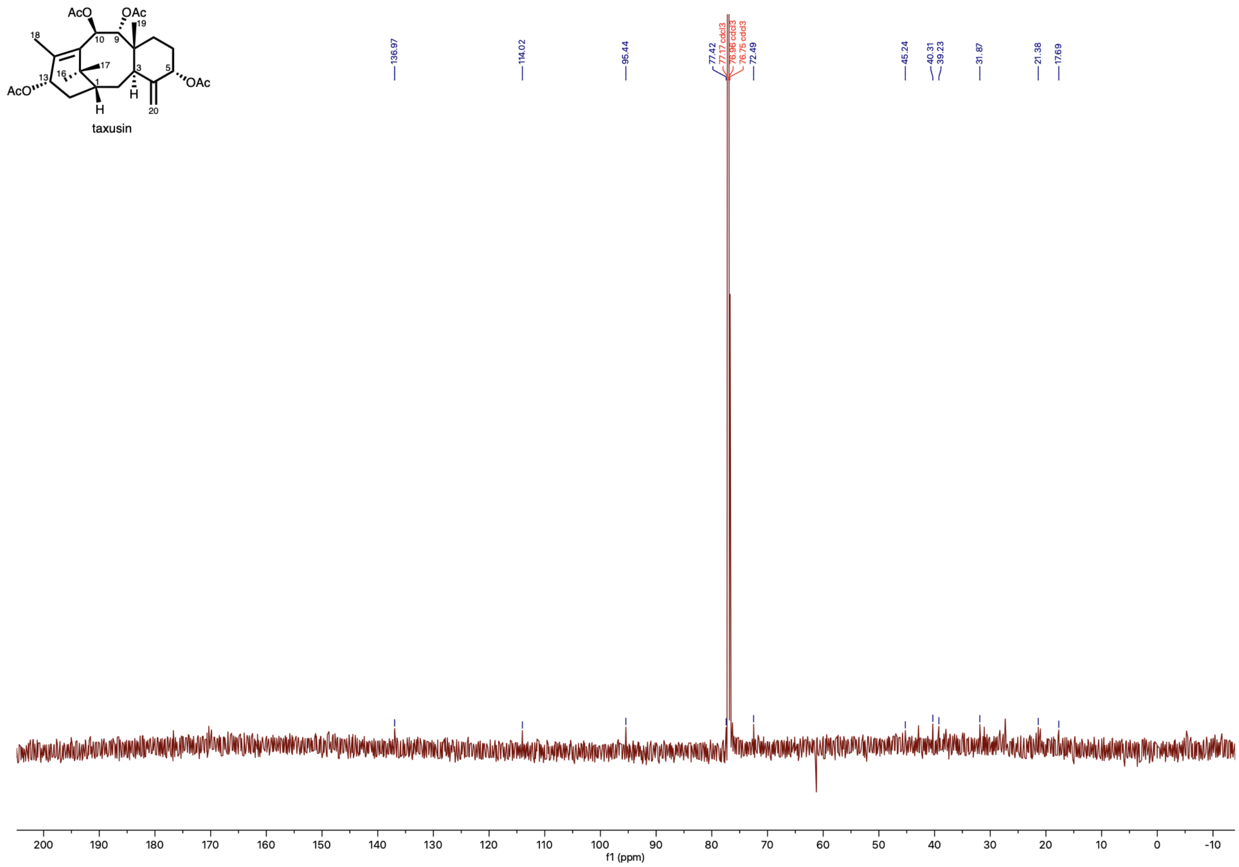


**Fig. S33. $^{13}$C-NMR spectrum of taxusin (6) in $CDCl_3$.**

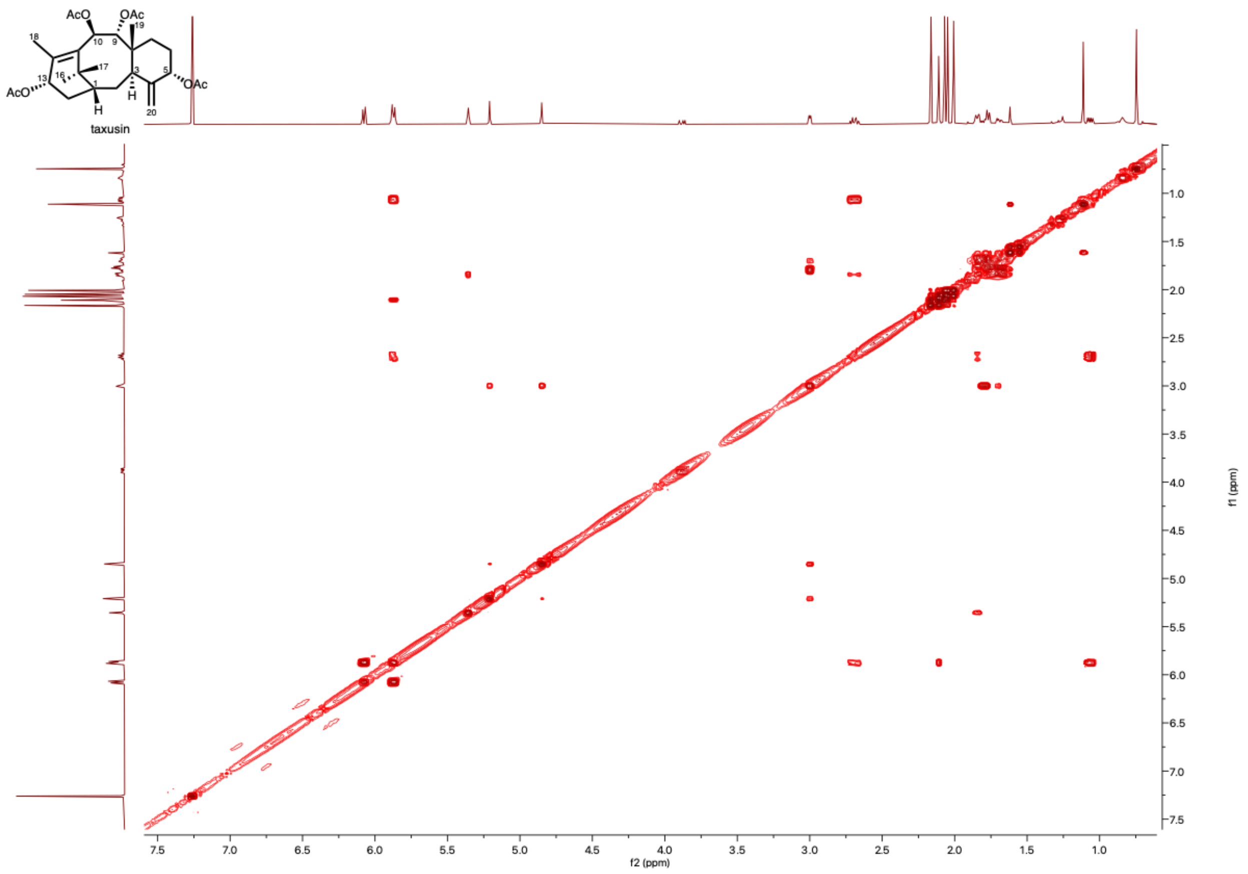


**Fig. S34. COSY spectrum of taxusin (6) in $CDCl_3$.**

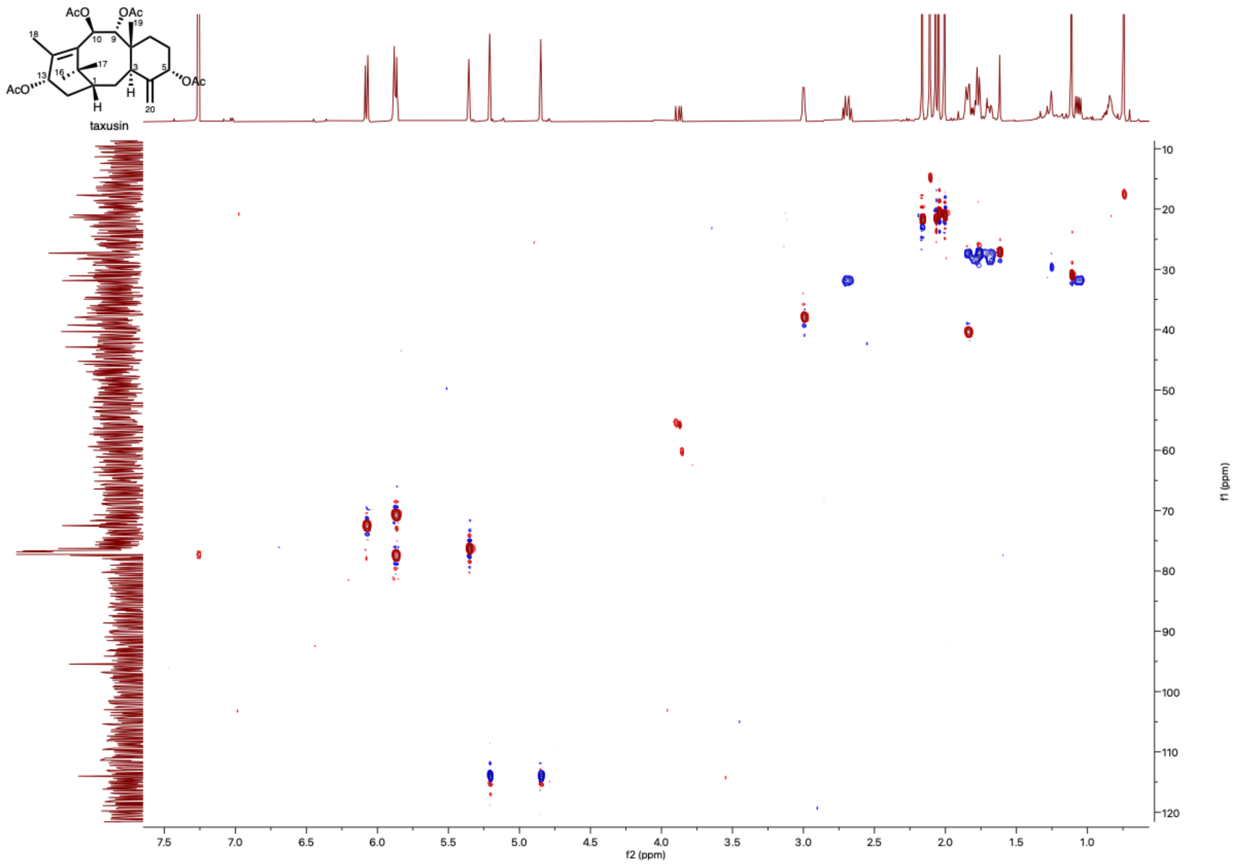


**Fig. S35. HSQC spectrum of taxusin (6) in $CDCl_3$.**

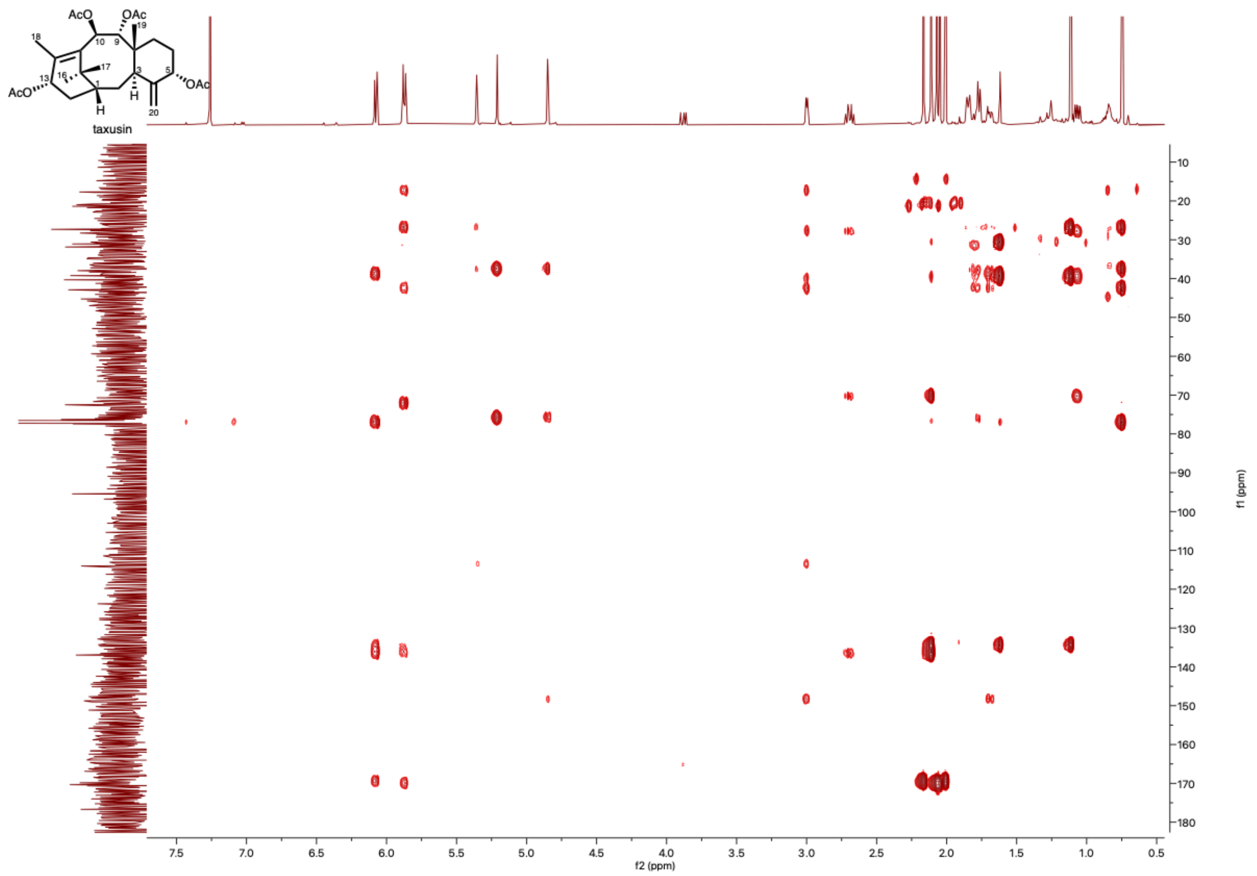


**Fig. S36. HMBC spectrum of taxusin (6) in $CDCl_3$.**

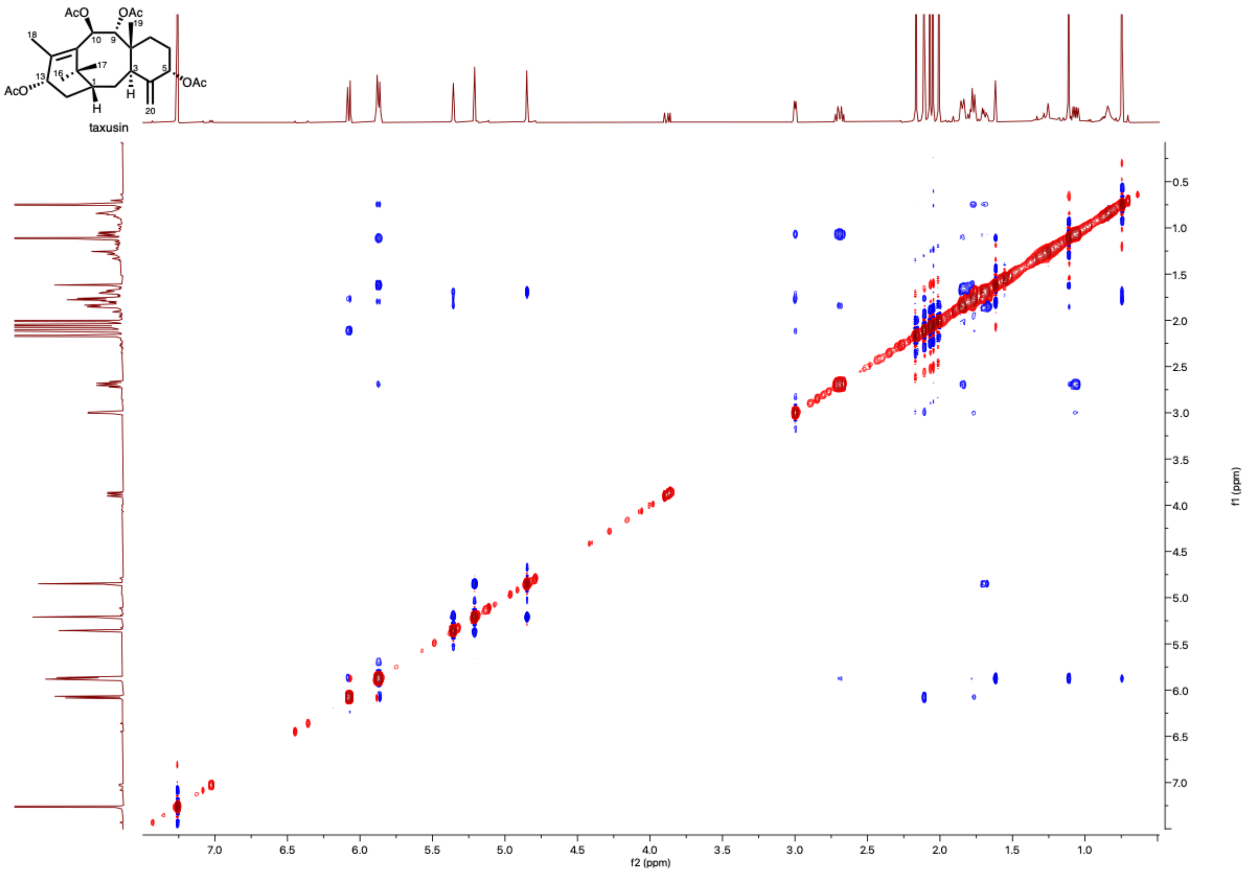


**Fig. S37. ROESY spectrum of taxusin (6) in $CDCl_3$.**

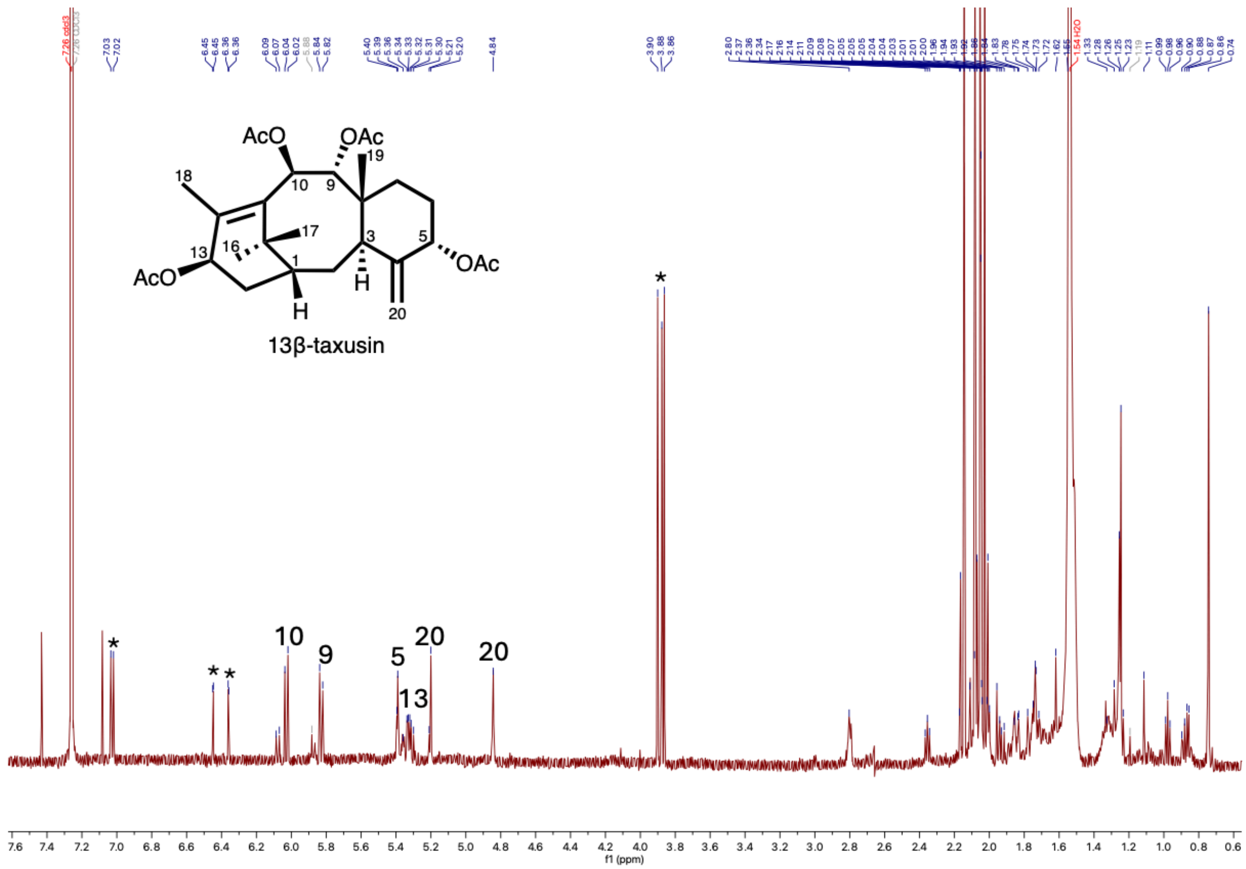


**Fig. S38. $^1$H-NMR spectrum of 13β-taxusin (6’) in $CDCl_3$.** Asterisk indicates impurities.

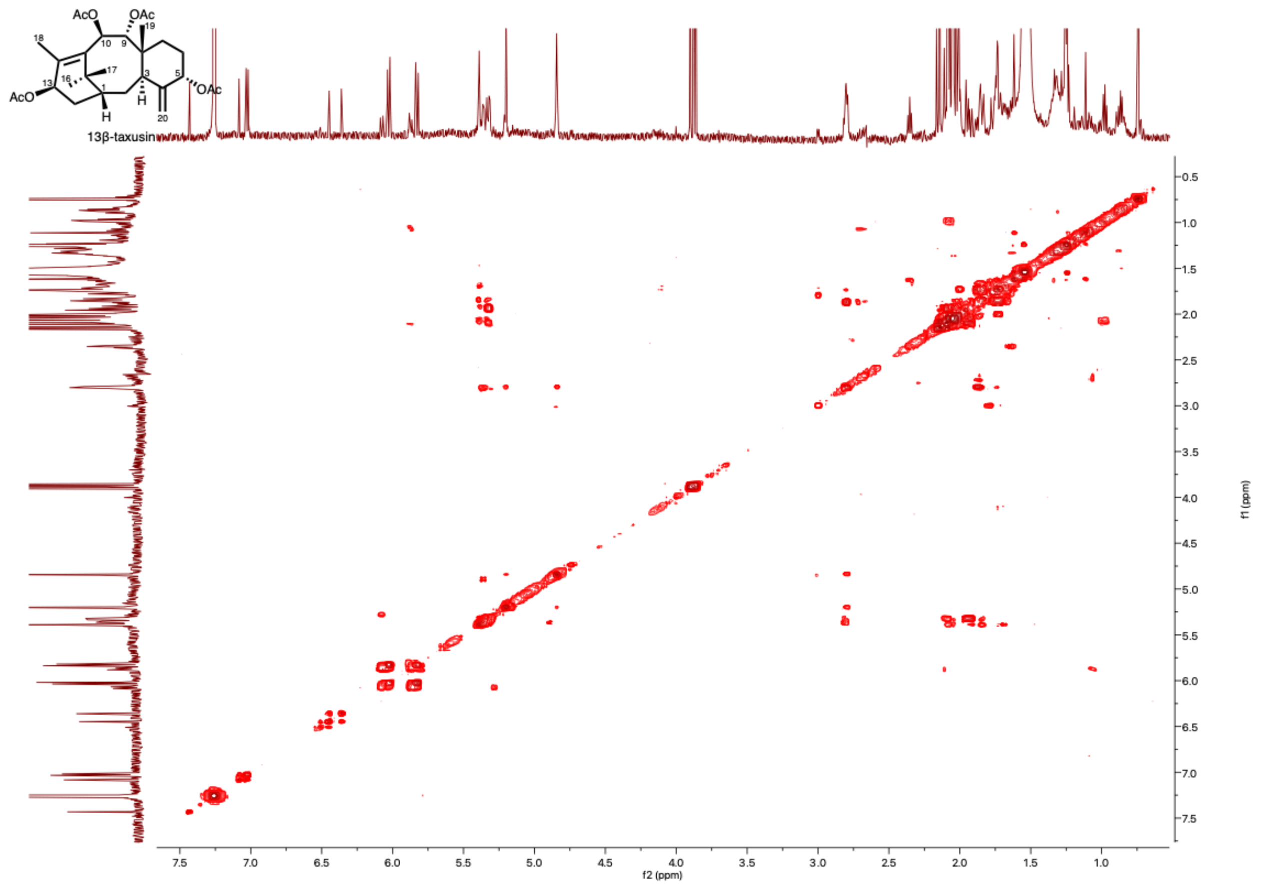


**Fig. S39. COSY spectrum of 13β-taxusin (6') in $CDCl_3$.**

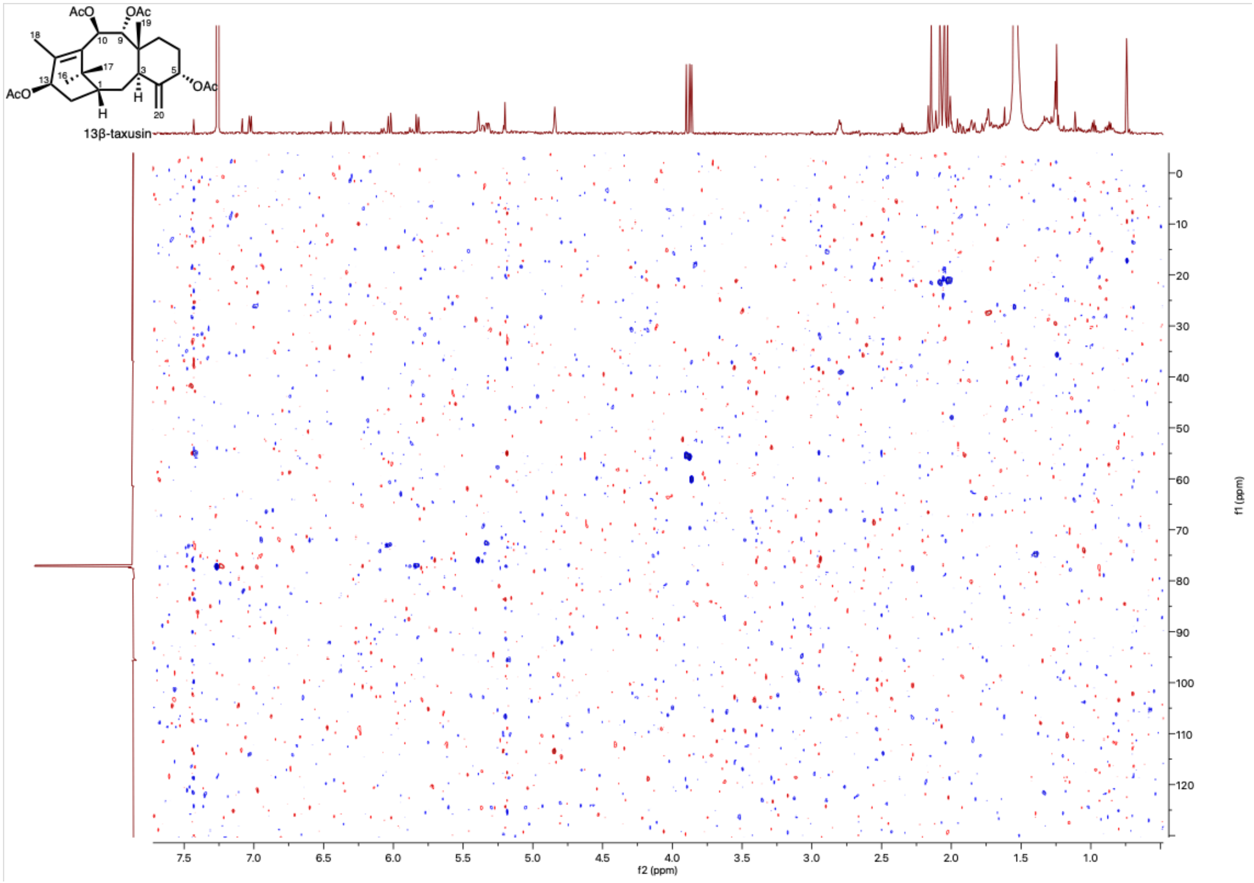


**Fig. S40. HSQC spectrum of 13β-taxusin (6') in $CDCl_3$.**

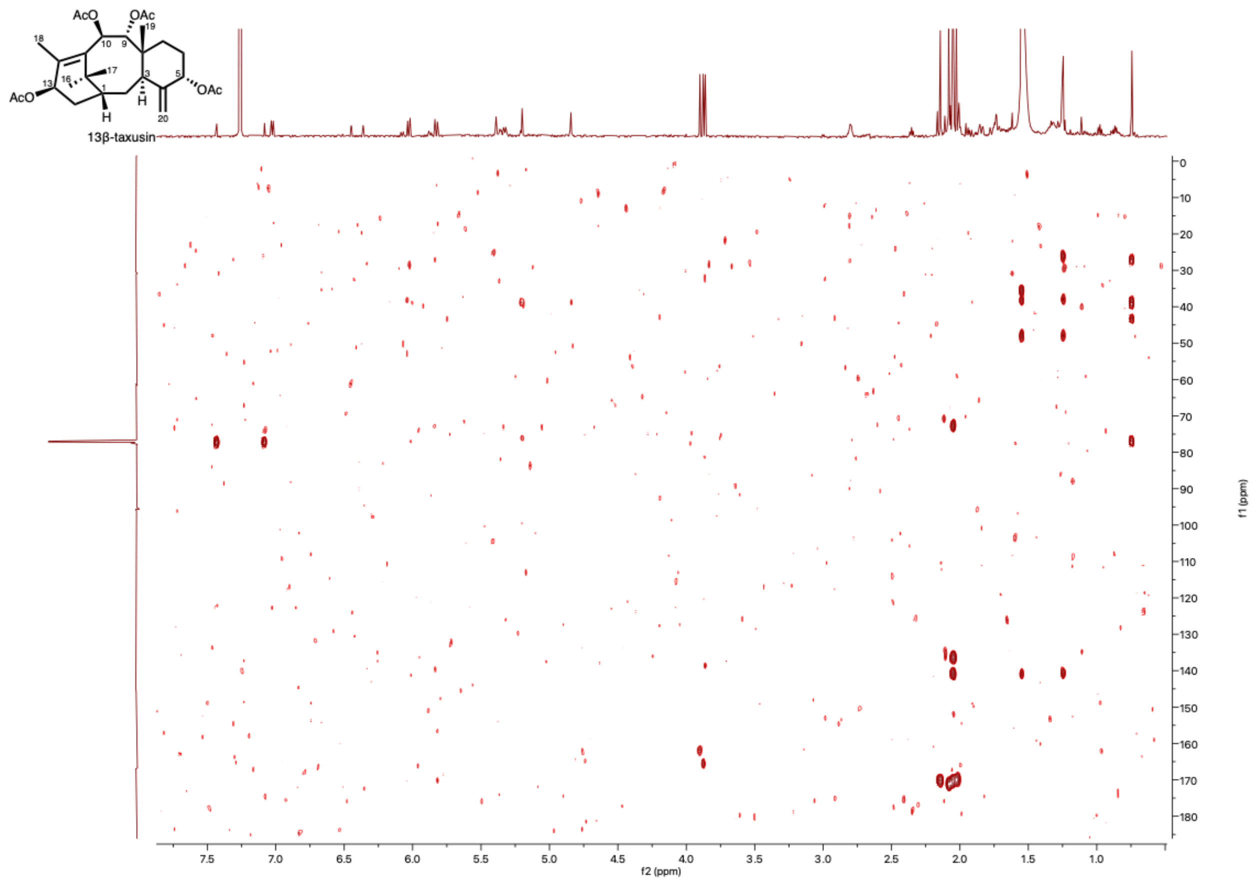


**Fig. S41. HMBC spectrum of 13β-taxusin (6') in $CDCl_3$.**

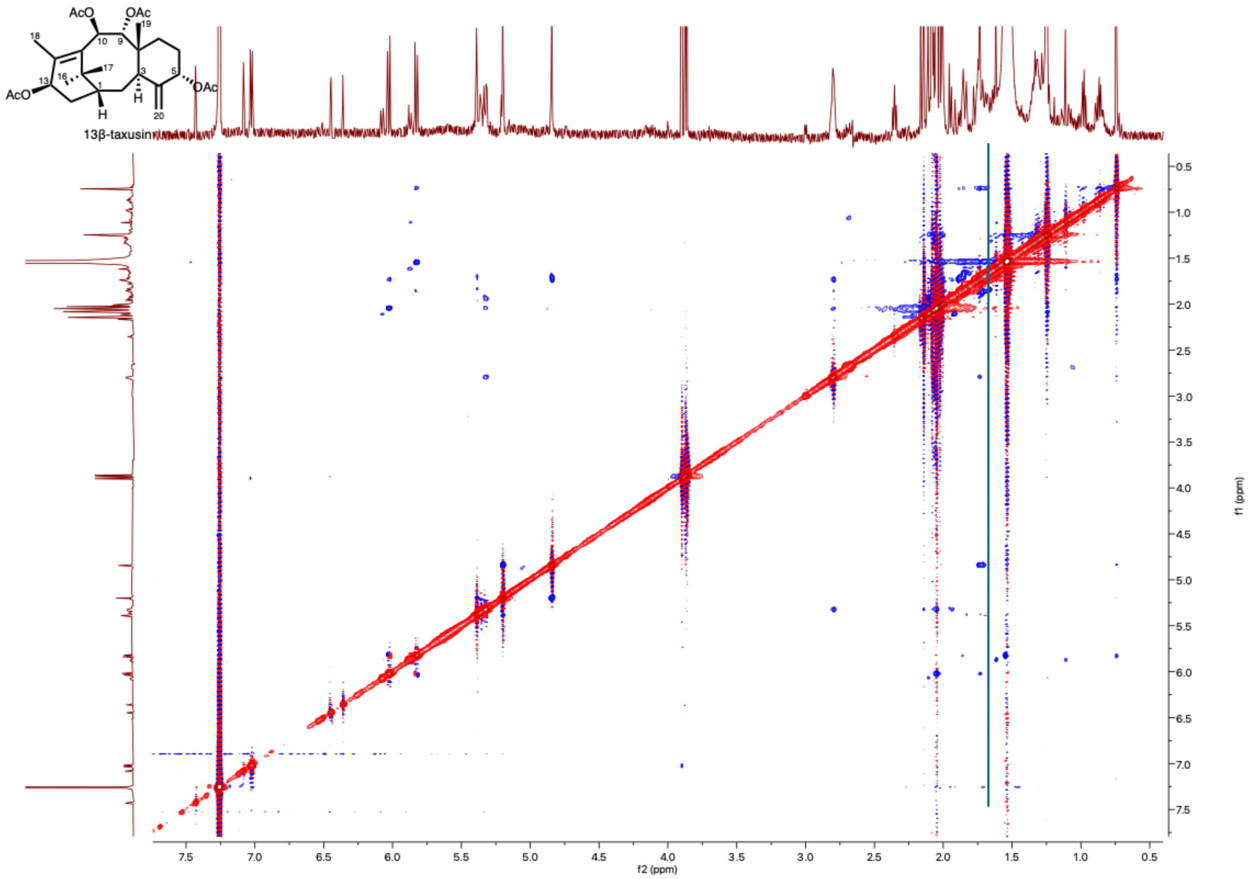


**Fig. S42. ROESY spectrum of 13β-taxusin (6’) in $CDCl_3$.**

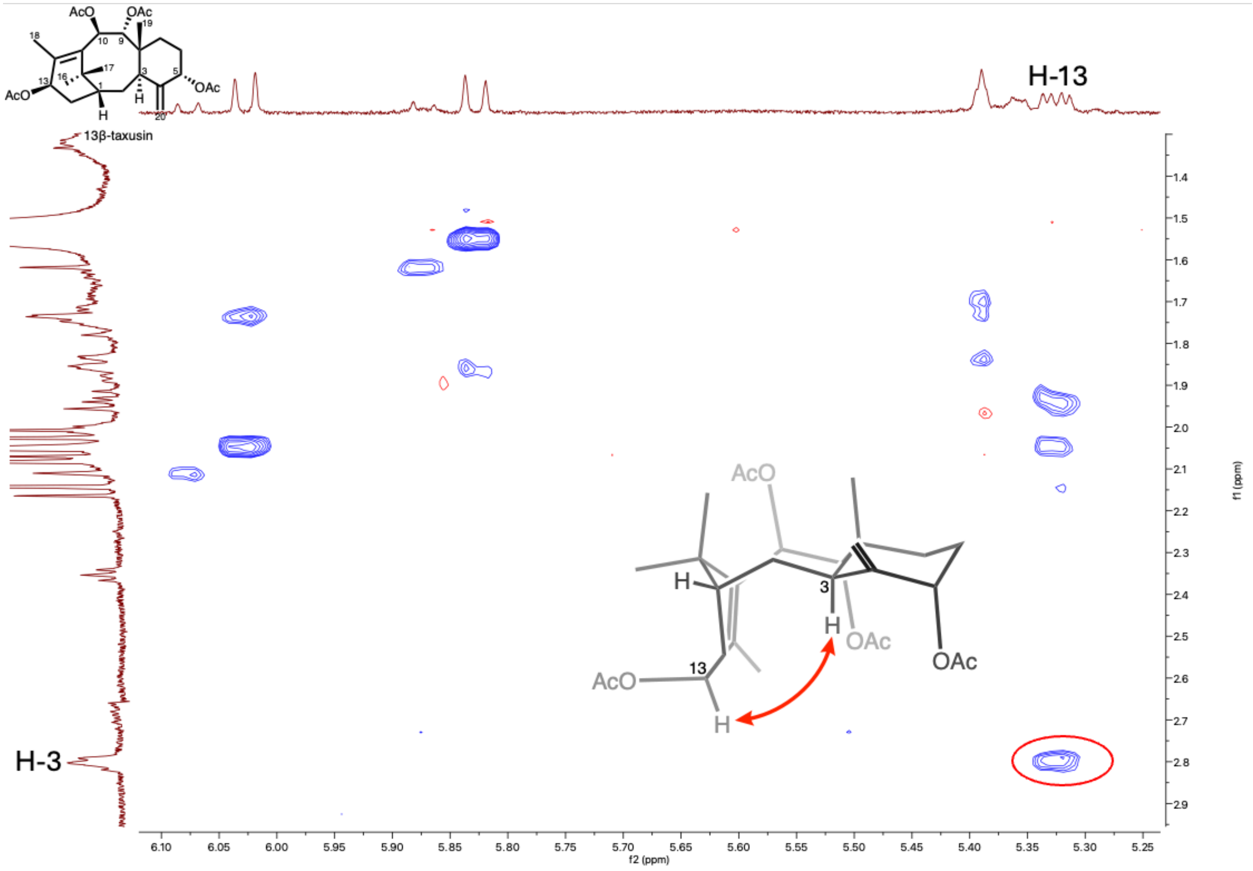


**Fig. S43. ROESY spectrum of 13β-taxusin (6') in $CDCl_3$ showing key ROESY correlations.**

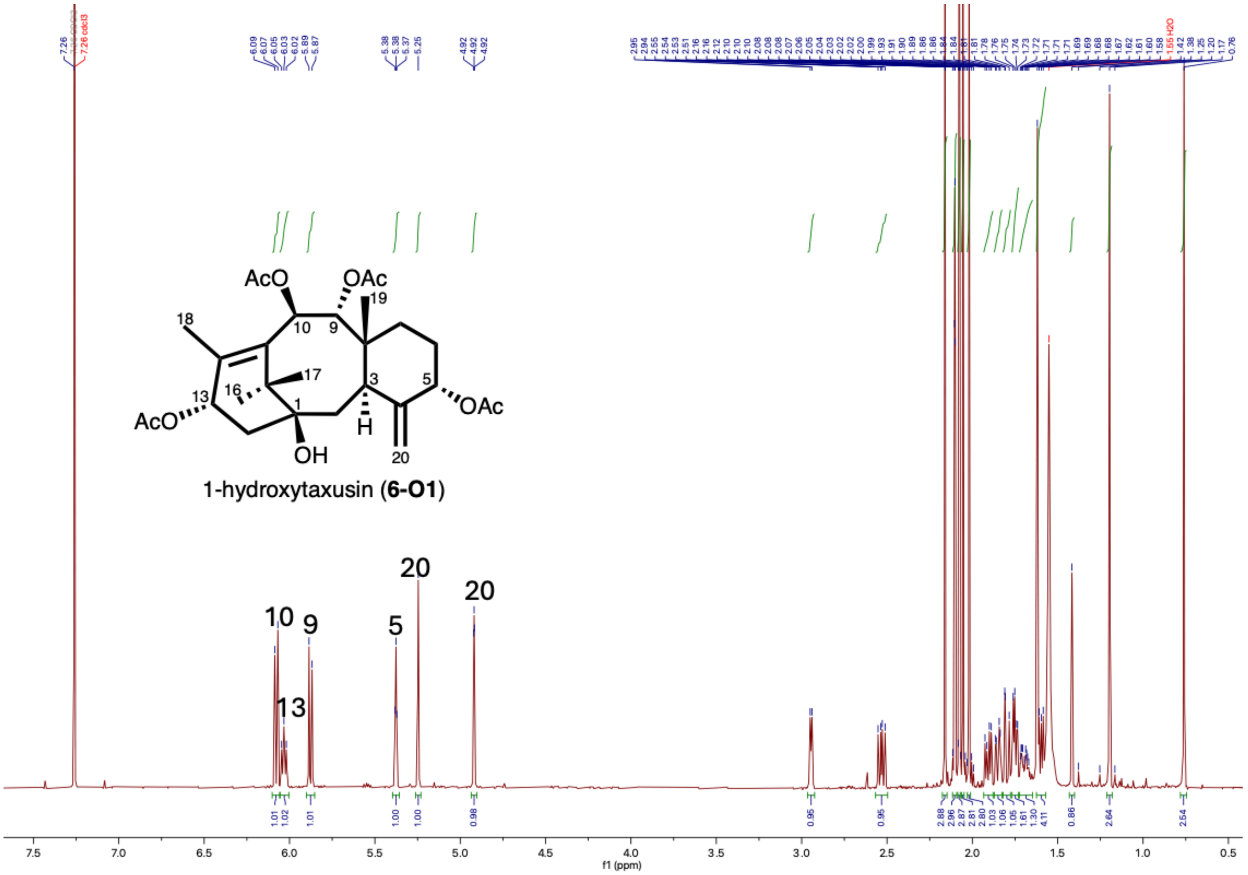


**Fig. S44.** $^{1}$**H-NMR spectrum of 1β-hydroxytaxusin (6-O1) in** $CDCl_3$**.**

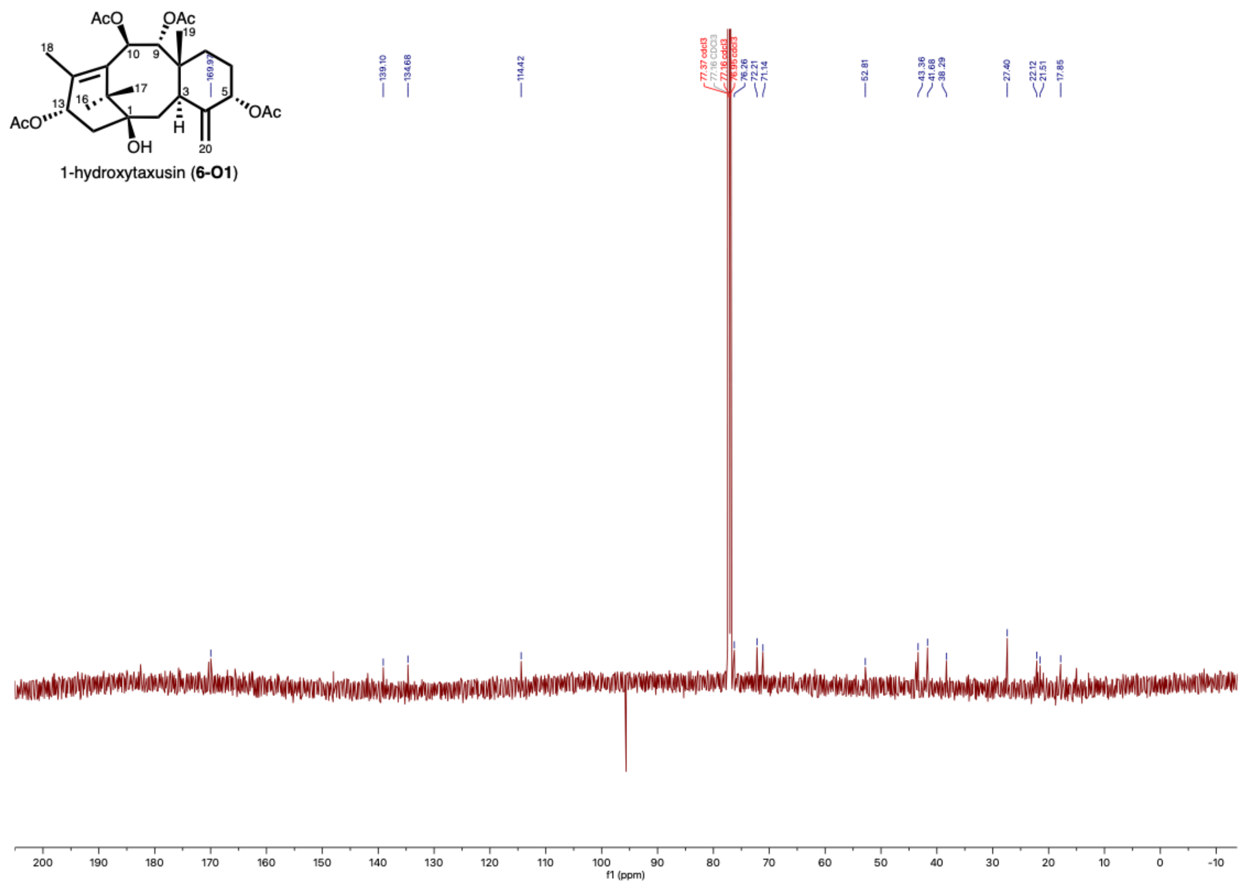


**Fig. S45. $^{13}$C-NMR spectrum of 1β-hydroxytaxusin (6-O1) in $CDCl_3$.**

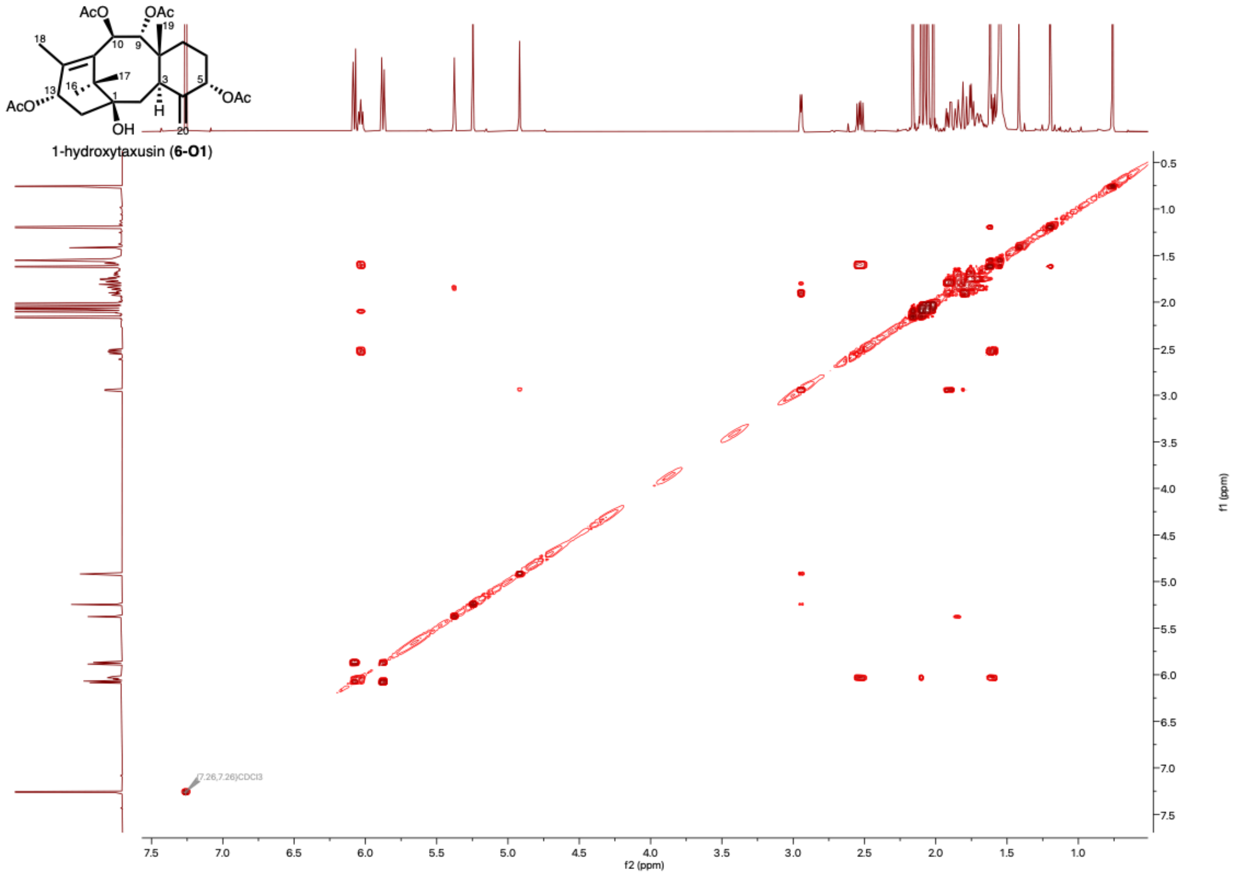


**Fig. S46. COSY spectrum of 1β-hydroxytaxusin (6-O1) in $CDCl_3$.**

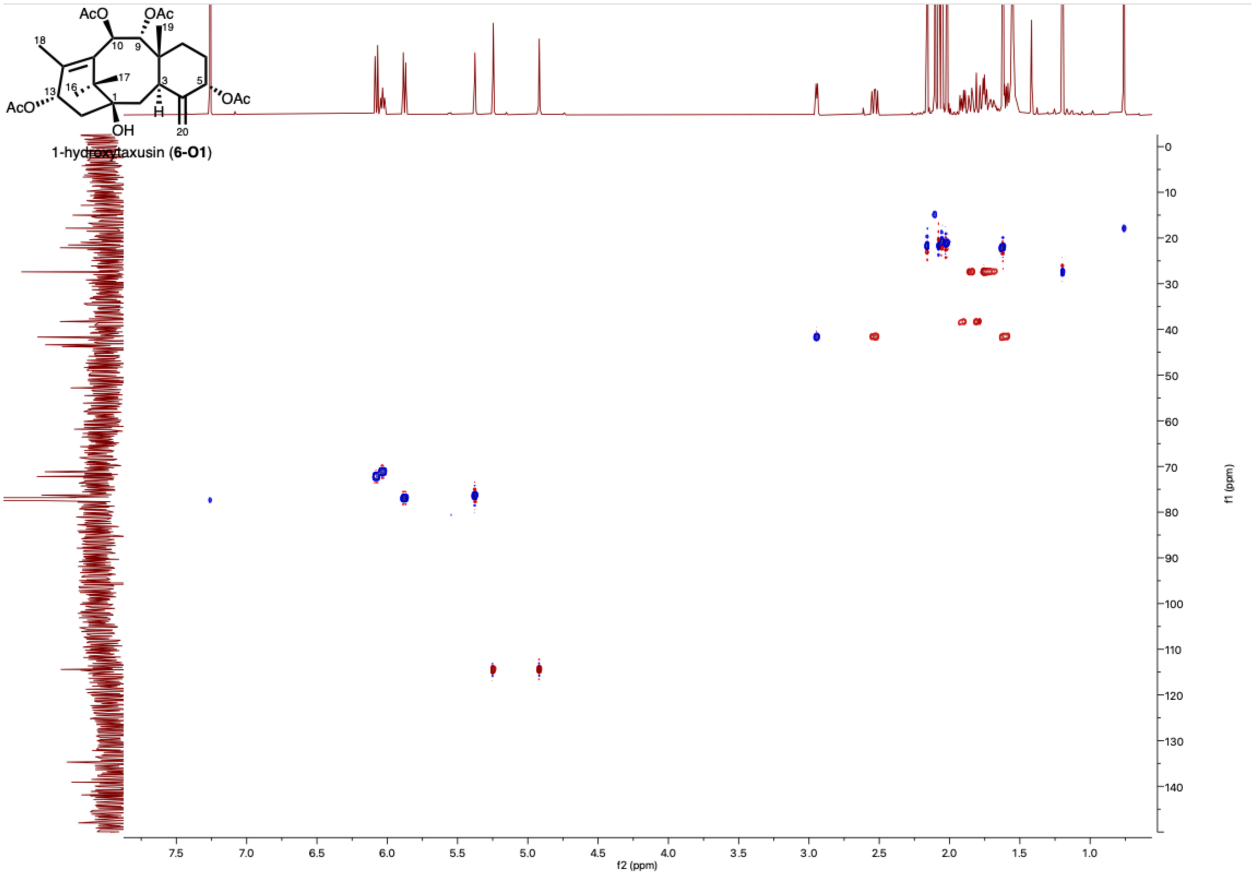


**Fig. S47. HSQC spectrum of 1β-hydroxytaxusin (6-O1) in $CDCl_3$.**

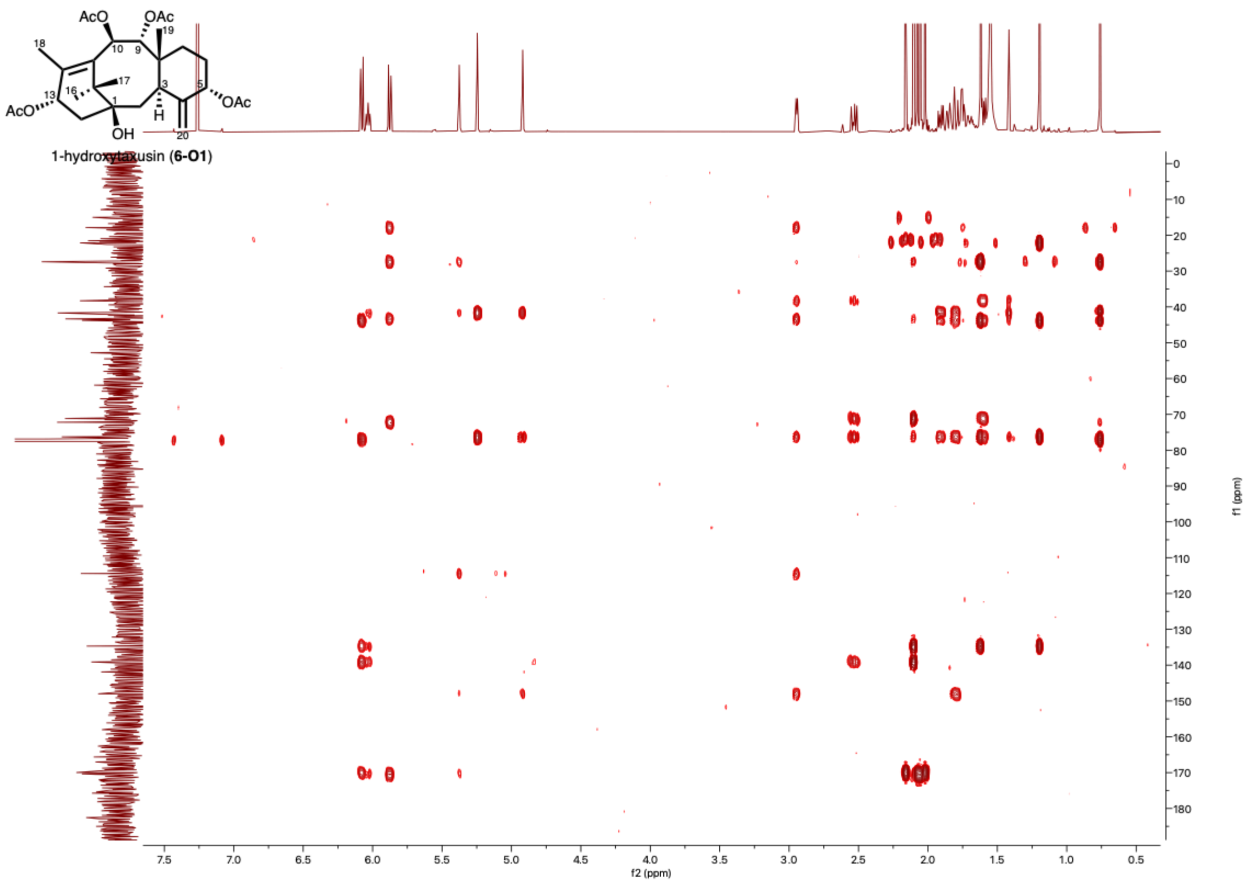


**Fig. S48. HMBC spectrum of 1β-hydroxytaxusin (6-O1) in $CDCl_3$.**

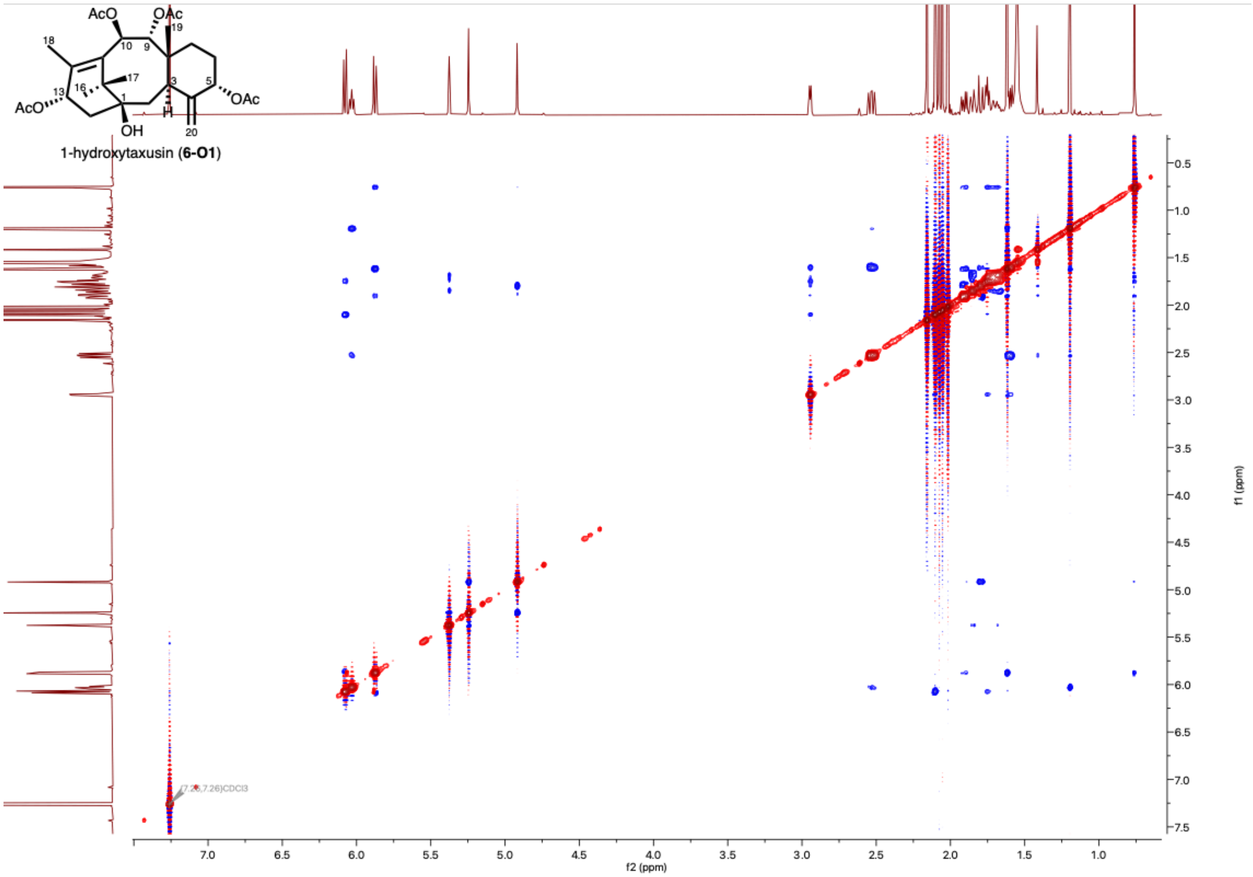


**Fig. S49. ROESY spectrum of 1β-hydroxytaxusin (6-O1) in $CDCl_3$.**

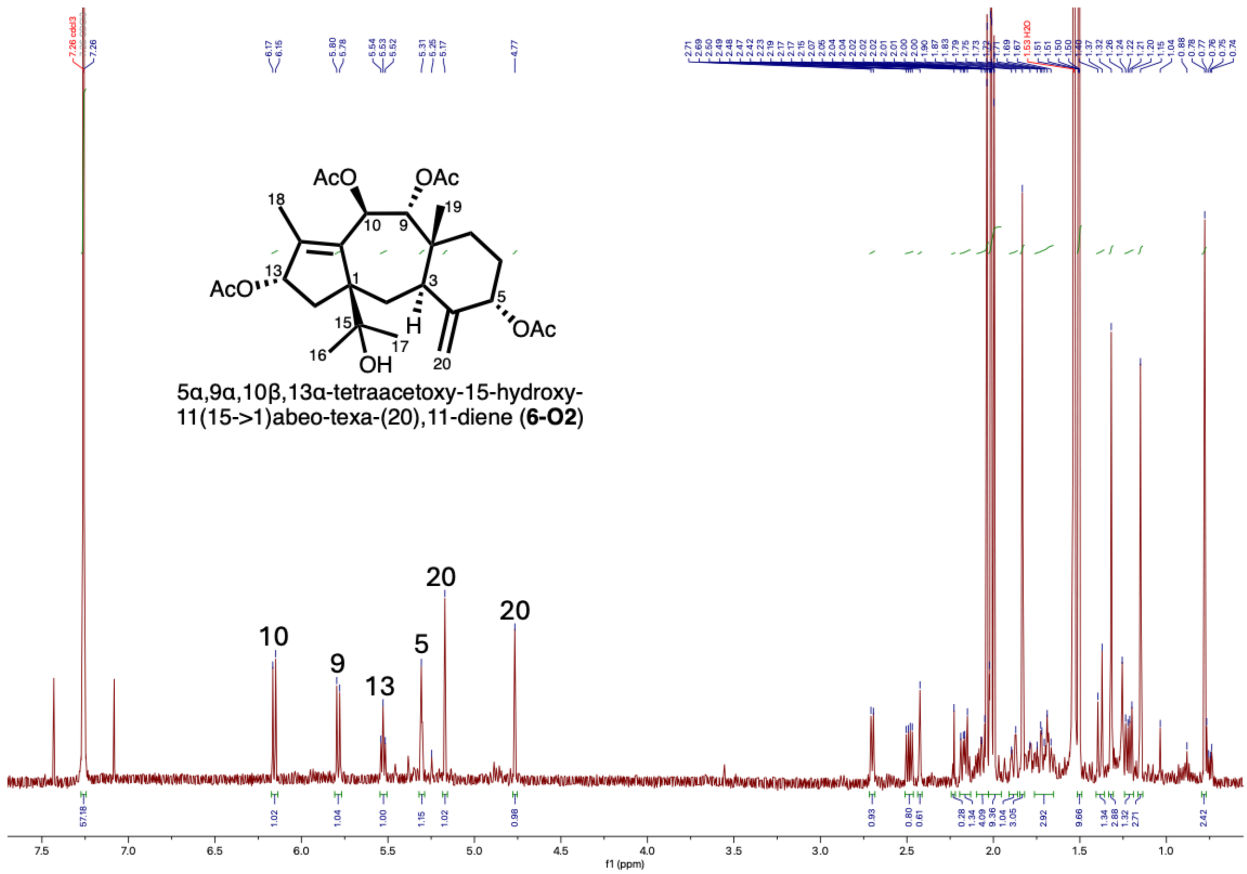


**Fig. S50.** ¹**H-NMR spectrum of 15-hydroxy-11(15→1)*abeo*-taxusin (6-O2) in $CDCl_3$.**

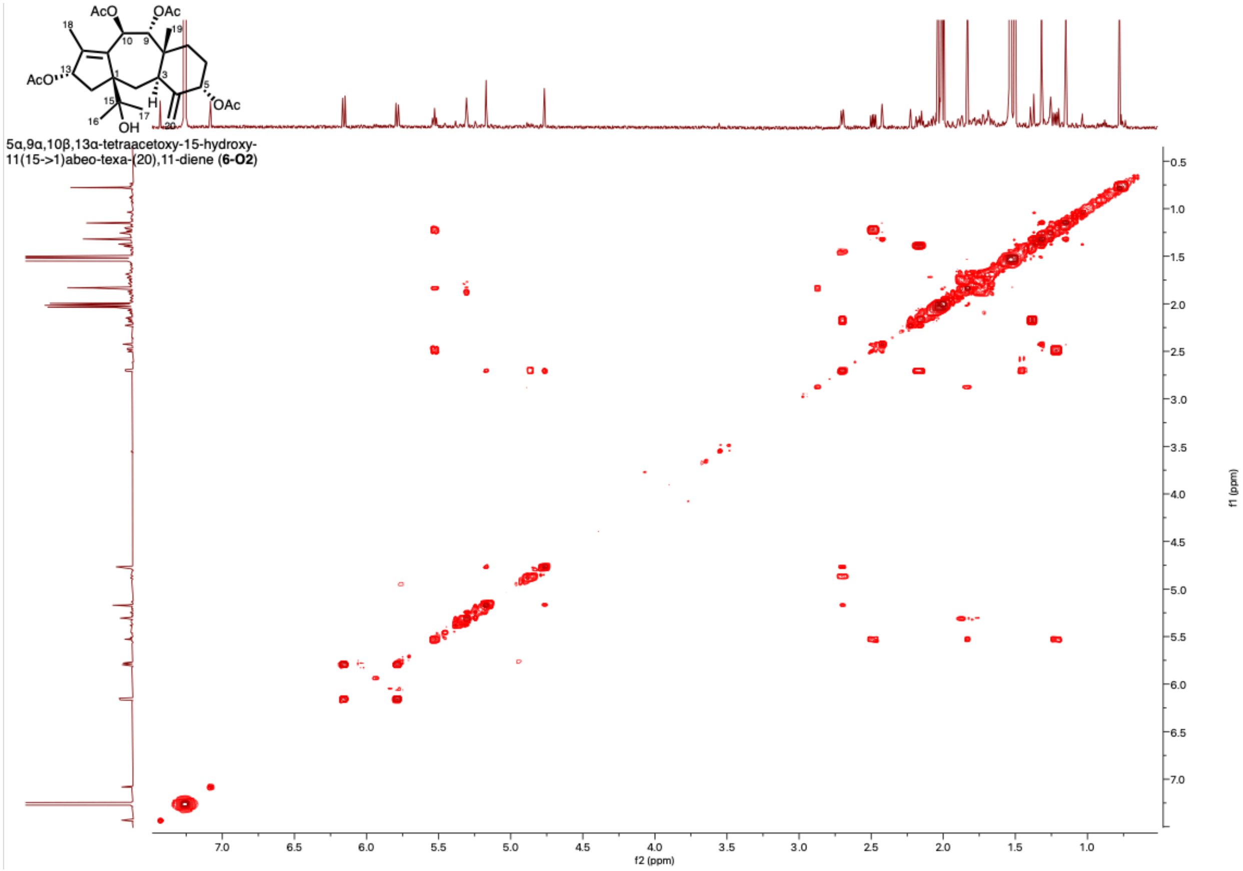


**Fig. S51. COSY spectrum of 15-hydroxy-11(15→1)*abeo*-taxusin (6-O2) in $CDCl_3$.**

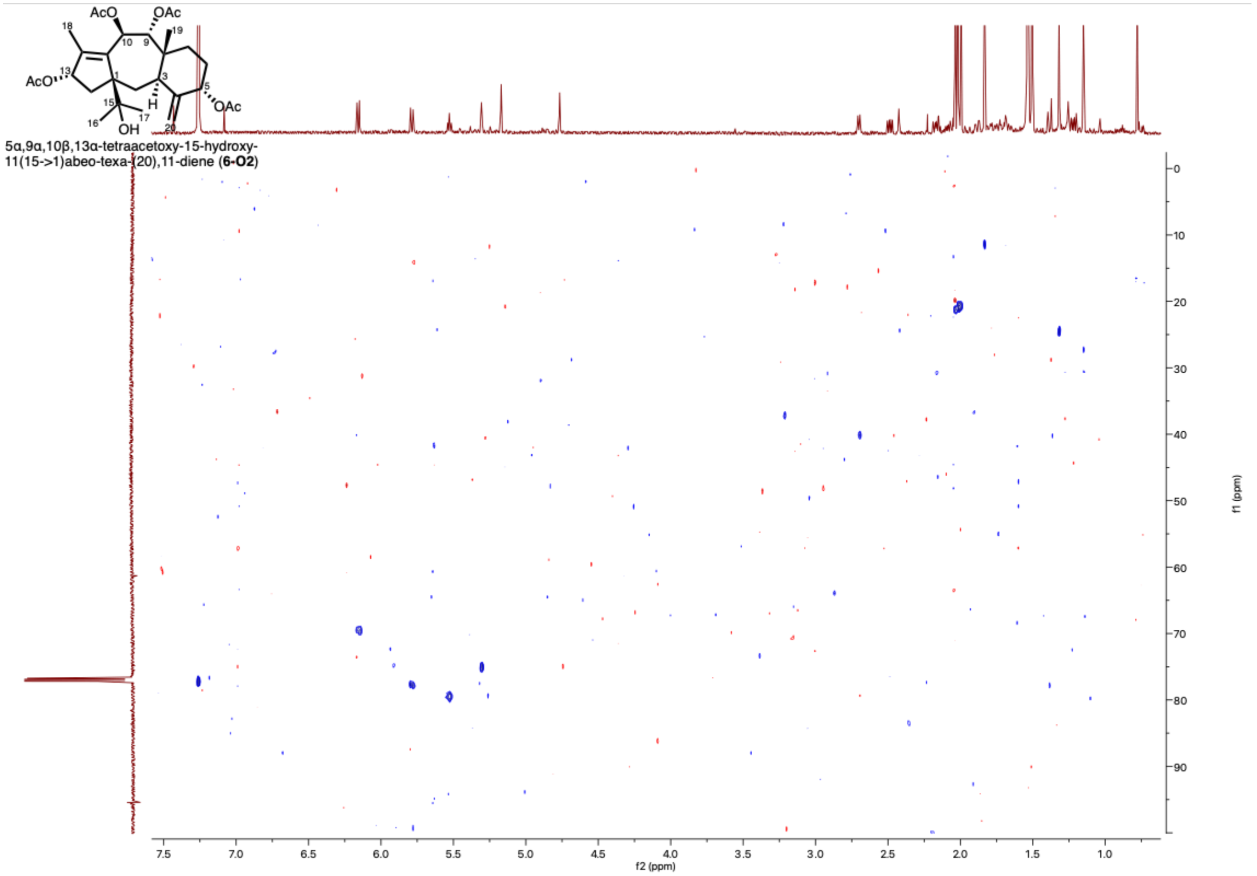


**Fig. S52. HSQC spectrum of 15-hydroxy-11(15→1)*abeo*-taxusin (6-O2) in $CDCl_3$.**

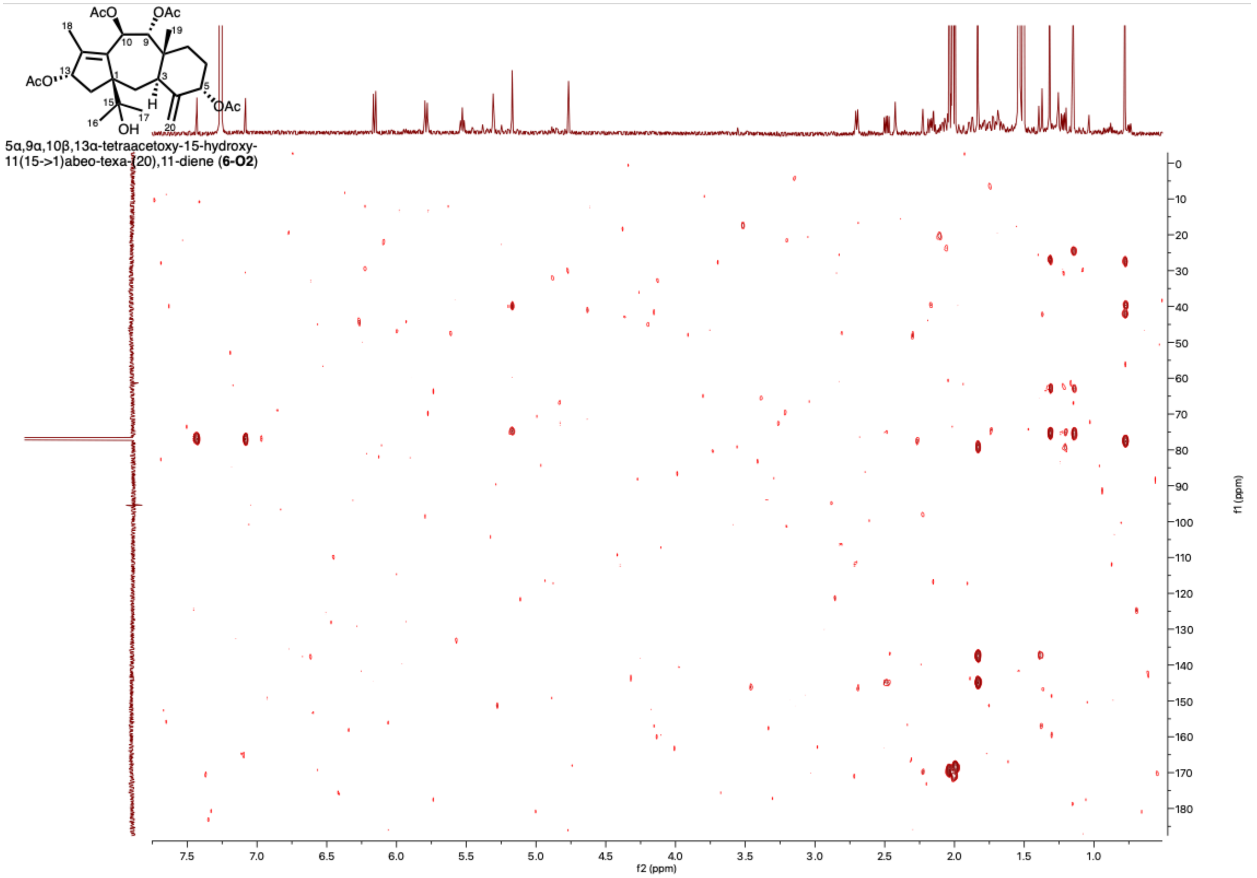


**Fig. S53. HMBC spectrum of 15-hydroxy-11(15→1)*abeo*-taxusin (6-O2) in $CDCl_3$.**

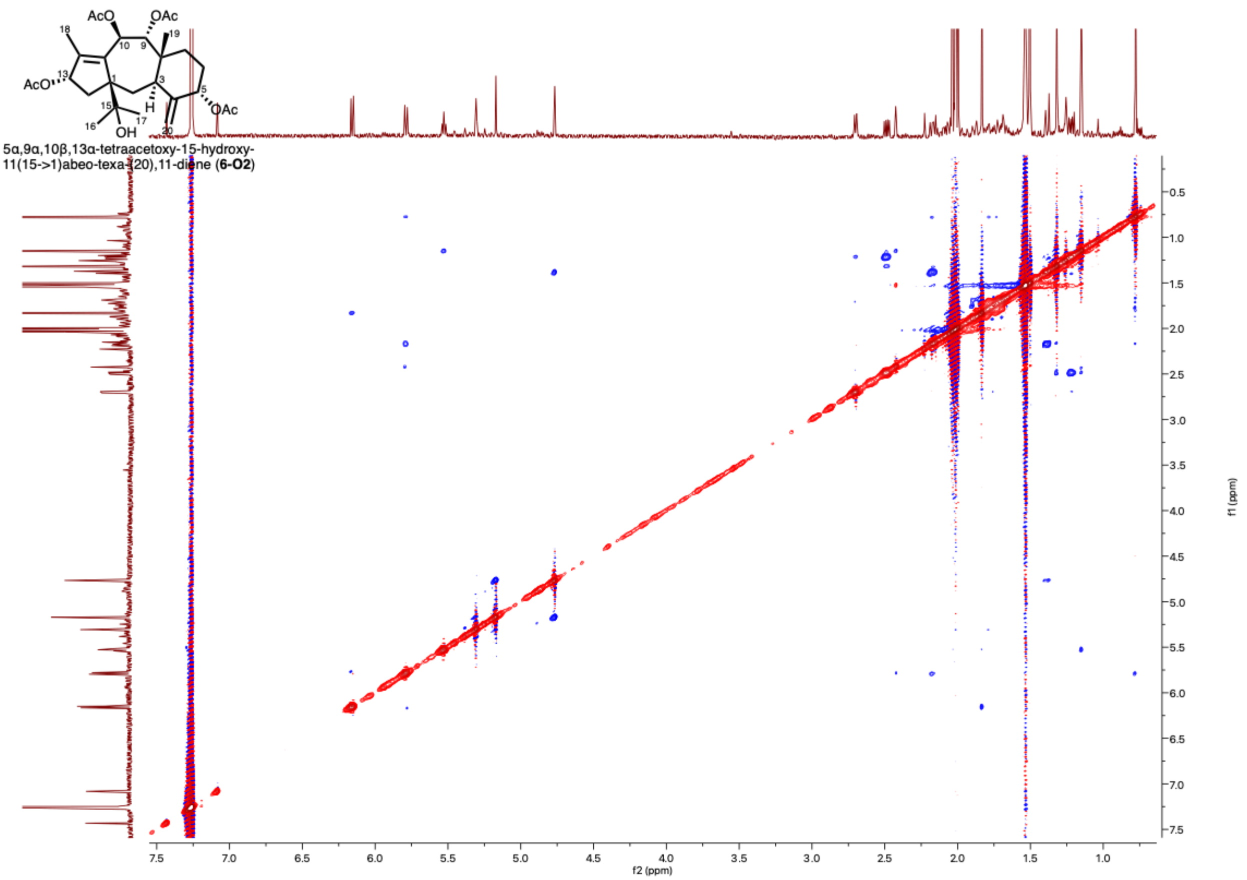


**Fig. S54. ROESY spectrum of 15-hydroxy-11(15→1)*abeo*-taxusin (6-O2) in $CDCl_3$.**

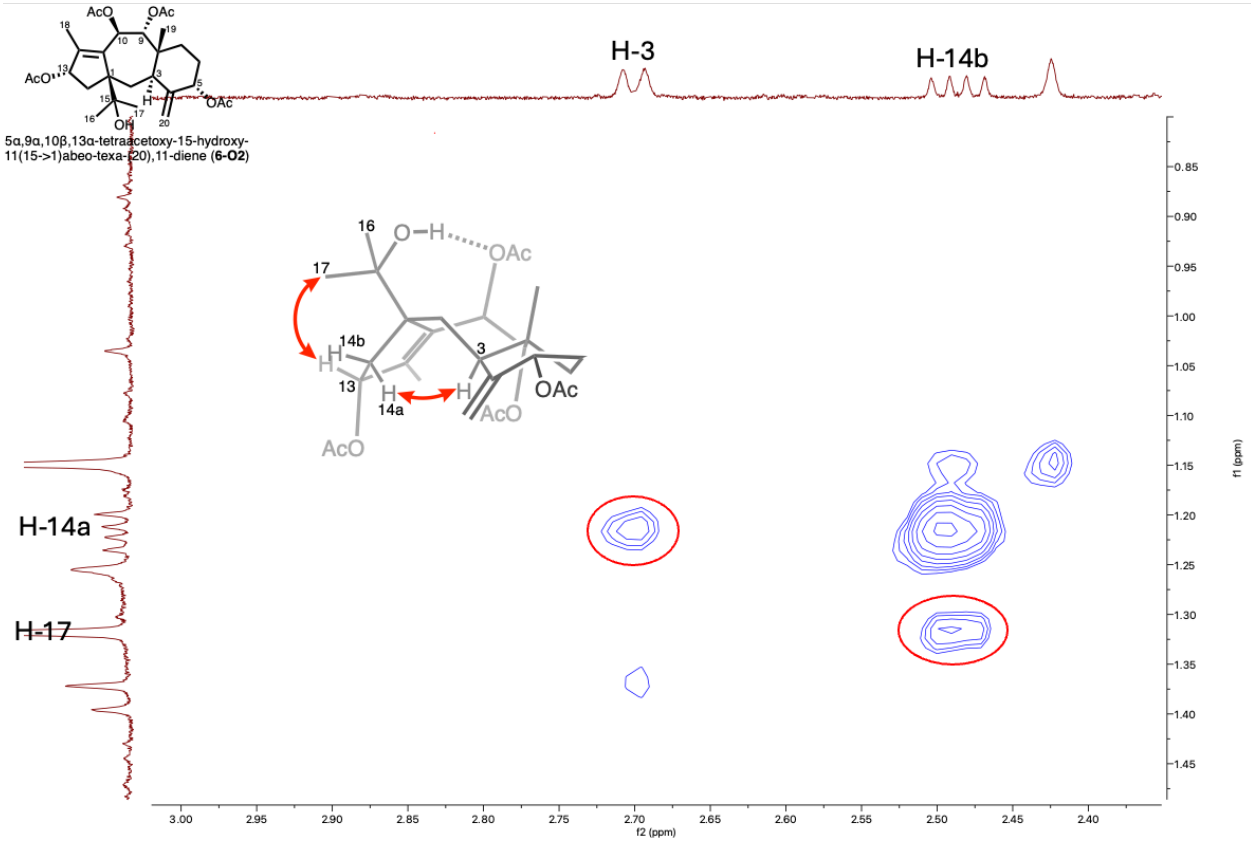


**Fig. S55. ROESY spectrum of 15-hydroxy-11(15→1)*abeo*-taxusin (6-O2) in $CDCl_3$ showing key ROESY correlations.**

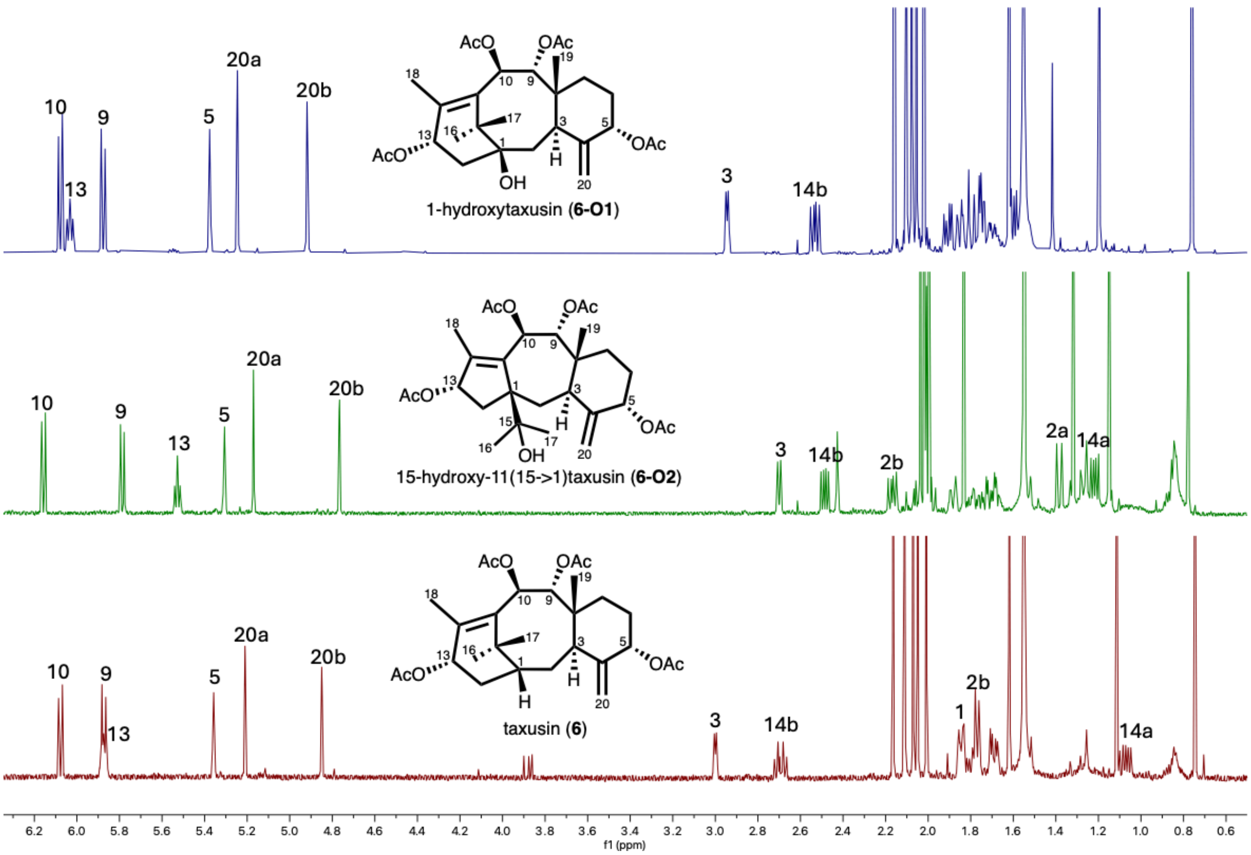


**Fig. S56.** $^1$**H-NMR spectra of taxusin (6), 1β-hydroxytaxusin (6-O1), and 15-hydroxy-11(15→1)*abeo*-taxusin (6-O2) in $CDCl_3$.**
